# Never cared for what they do. High structural stability of Guanine-quadruplexes in presence of strand-break damages

**DOI:** 10.1101/2021.12.01.470770

**Authors:** Tom Miclot, Cécilia Hognon, Emmanuelle Bignon, Alessio Terenzi, Stéphanie Grandemange, Giampaolo Barone, Antonio Monari

**Affiliations:** Department of Biological, Chemical and Pharmaceutical Sciences. University of Palermo, via delle Scienze 90126 Palermo, Italy; Université de Lorraine and CNRS, LPCT UMR 7019, F-54000 Nancy, France; Université de Lorraine and CNRS, CRAN UMR 7039, F-54000 Nancy, France; Université de Paris and CNRS, ITODYS, F-75006 Paris, France

**Keywords:** DNA strand breaks, G-quadruplexes, Ionizing radiations, Molecular dynamics simulations, Cancer resistance

## Abstract

DNA integrity is an important factor to assure genome stability and, more generally, cells and organisms’ viability. In presence of DNA damage, the normal cell cycle is perturbed while cells activate their repair processes. Although efficient, the repair system is not always able to ensure the complete restoration of gene integrity. In these cases, not only mutations may occur, but the accumulation of lesions can either lead to carcinogenesis or reach a threshold which induces apoptosis and the programmed cell death. Among the different types of DNA lesions, strand breaks produced by ionizing radiations are the most toxic, due to their inherently difficult repair, which may lead to genomic instability. In this article we show, by using classical molecular simulations techniques, that differently from the canonical double-helical B-DNA, guanine-quadruplex (G4) arrangements show a remarkable structural stability, even in presence of two strand breaks. Since G4-DNA are recognized for their regulatory roles in cell senescence and gene expression, also involving oncogene, their stability can be related to an evolutionary cellular response aimed at minimizing the effects of ionizing radiation.

## Introduction

G-quadruplexes (G4s) are guanine-rich nucleotide sequences that form a non-canonical DNA or RNA secondary structure, stabilized by monovalent cations, in which guanine tetrads, interacting through Hoogsteen hydrogen-bonds (H-bonds), are stacked together. They can be formed both intra-strand (i.e., produced by the folding of a single-stranded DNA fragment) or inter-strand G-G pairing. G4s may adopt various topologies, depending on the orientation of the glycosidic bond, giving rise to parallel, antiparallel, and hybrid arrangements [1]. The rigid tetrad cores are connected by nucleotide loops, whose length and flexibility may experience rather large variations. The structural study of G4 is gaining momentum since the latter have been recognized to have important and versatile biological implications. For example, DNA or RNA G4s may regulate viral infection cycles, becoming, as a consequence, potential therapeutic targets for antiviral drug candidates [2,3] For this reason, they are also of major interest in the context of the current pandemic caused by the infectious pathogen SARS-CoV-2, whose genome has been shown to present G4-compatible regions [4–8]. Interesting and recent reviews on the antiviral possibilities offered by G4 can be found in literature [9–11]. G4s are also involved in some neurological diseases, such as Alpha-thalassaemia or X-linked intellectual disability syndrome, in which they are positively or negatively involved in a cascade of gene expression regulations [12–14]. From a cellular point of view, G4s are involved in DNA replication pathways [15,16], as well as in gene expression, since they have been localized in oncogene and viral DNA promoter regions [17–20]. In addition, G4 are abundant in the chromosomes’ terminal sequences, the telomeres. In this respect they play a fundamental role in regulating the cellular life cycle controlling the replication-induced shortening of the telomeres and hence cellular programmed death via inhibition of the telomerase [21–24]. As a matter of fact, the disruption of this mechanisms is linked to the immortality phenotype of cancer cells which makes G4s ideal targets for cancer chemotherapeutic agents. Remarkably, it has been shown that G4s are involved in conferring *Deinococcus radiodurans* its extraordinary resistance to ionizing radiations [25,26].

Also because of their versatile and rather ubiquitous biological roles, several studies have been focusing on the effect of different DNA damages on the stability of G4s. In particular, oxidative damages have been particularly scrutinized, due to the fact that G4s are inherently composed of guanine-rich sequences and the latter is the most easily oxidized nucleotide, being its most common oxidation product 8-oxo-guanine (8-OxoG) [27–30]. Although G4s are clearly considered hotspots for oxidative DNA damages, they have shown strong structural resistance to this class of lesion [31], also depending on the amount of oxidative lesions and on their position in the DNA backbone [31–34].

In addition to base modification or deletion, oxidative stress [35,36] and ionizing radiation [37] are able to induce DNA strand-break damage. These lesions, in particular strand breaks in two different points of the same DNA strand, are among the most toxic DNA damages, leading to a very strong genome instability, usually resulting in cell death. This is essentially due to the difficulty in repairing the dispersed DNA fragments. Interestingly, radiation resistant bacteria present even specific DNA binding proteins that colocalize at the lesions foci favoring, in this way, the repair [38]. Ionizing radiation can result in two kind of strand-breaks: the canonical 5’- PO_4_- / 3’-OH (CA), and non-canonical 5’- OH / 3’-PO_4_^-^ (NC) [39,40], as shown in Figure 1. Both kinds of DNA terminations are already known in biological systems, since they are respectively produced by the activity of deoxyribonucleases I and II [41], respectively. However, from a more global cellular point of view, the two types of damages are not equivalent: CA strand breaks are also commonly produced by normal cellular processes, notably in replication and repair [42], and thus are more easily recognized by DNA ligase, which may catalyze the formation of a phosphodiester bond to repair the DNA damage [43,44]. Conversely, NC lesions are more hardly repaired within the cell, and are also usually produced by deoxyribonuclease II during programmed cell death pathways [45,46].

**Figure 1.**
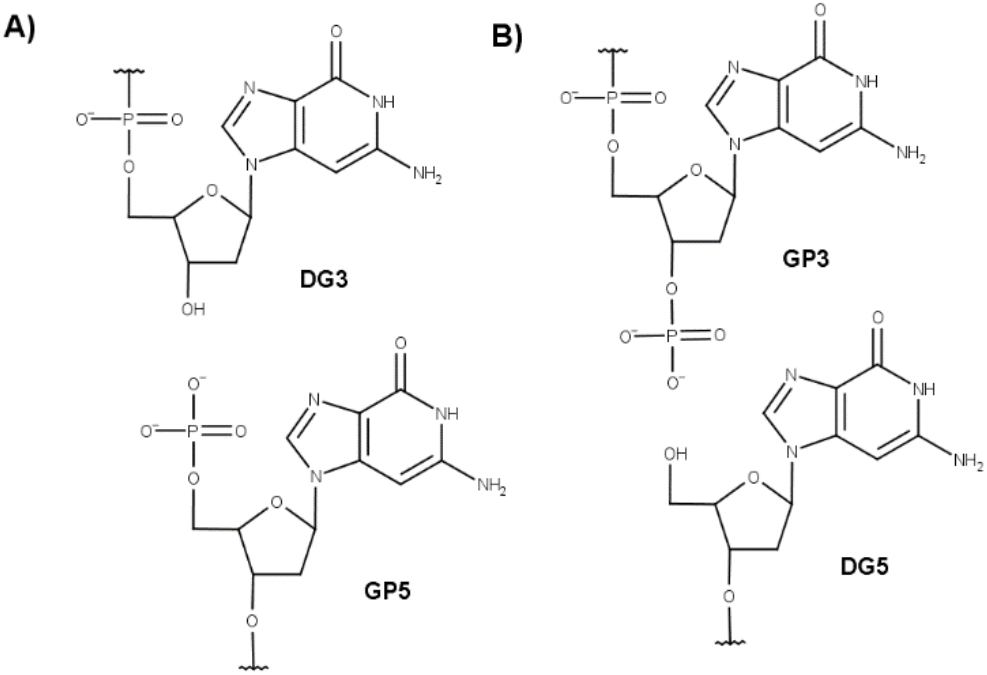
A) Canonical and non-canonical B) strand break damages occurring on the phosphodiester -O-P(O2)-O- bond of the nucleic acid.

However, some pathways allowing the reparation of NC strand breaks exist. As an example, mammalian polynucleotide kinase adds a phosphate group at the 5’ position of non-canonical terminations while replacing the -PO_4_^-^ moiety at the 3’ position with a hydroxyl group, hence permitting the further action of DNA ligase [47,48]. In addition, some organisms have developed proper mechanisms for repairing non-canonical strand breaks. One example is the repair pathway involving RNA ligase RtcB in Escherichia coli, or the one involving HD-Pnk in *Deinococcus radiodurans* [49,50].

Previous studies have highlighted the impact of G4s on strand break formation. Indeed, when G4s cannot be unfolded by helicases then DNA replication is stopped at their location and DNA breaks may occur [51,52]. They are also known to play an important role in radio-resistance and DNA damage response [25,53]. Despite their interesting features, the impact of the occurrence of strand breaks lesions in G4s from a structural and atomistic point of view is only poorly documented. Recently, Kumari et al. [53] have experimentally demonstrated the resistance of G4s to ionizing radiation and their presence in coding DNA sequence (CDS).

Starting from these considerations, in this contribution we rationalize the structural effects of the presence of strand breaks in G4-forming DNA sequences containing both CA and NC lesions. To this aim we have used extensive state-of-the-art molecular modelling and simulation techniques, in particular molecular dynamic (MD) simulations, to check how the number and position of strand breaks affect the structure of a parallel G4 structure, extracted from the human telomeric sequence (h-telo). Our results show that G4 are extremely resistant to strand-breaks, an occurrence which may be correlated to a possible protective role exerted in conditions of high ionizing or oxidative stress.

## Results and Discussion

The parallel h-telo G4 DNA has been used as our model, due to its presence in cells, where it protects telomeres by acting as a telomerase inhibitor [54]. In addition to the undamaged quadruplex used as a reference, 27 different structures harboring strand breaks have been taken into account. For both CA and NC forms, 6 single-breaks and 6 double-breaks have been introduced into the tetrads. Breaks have also been introduced into the loops of the CA form at 3 different positions, as shown in Table 1 and Figure 2. This choice allowed us to rationalize the role of the position of the strand breaks on the structural stability of G4s. Note that, the chosen strand break pattern follows a similar scheme used by us to study the impact of 8-oxoG on the structural stability of G4s [55].

**Figure 2.**
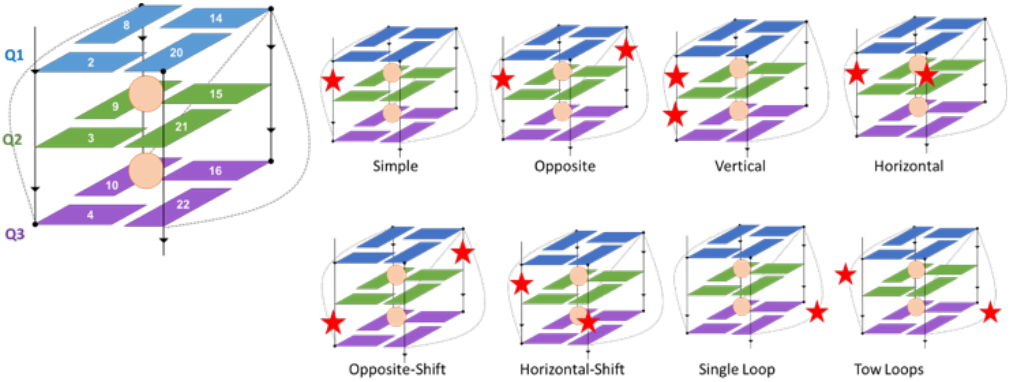
Position of the tetrads in the studied h-telo G4 and the relative orientation of the strand breaks damages (displayed as red stars) in peripheral loops or tetrad-connecting backbone. The orange dots represent the cations.

**Table 1.**
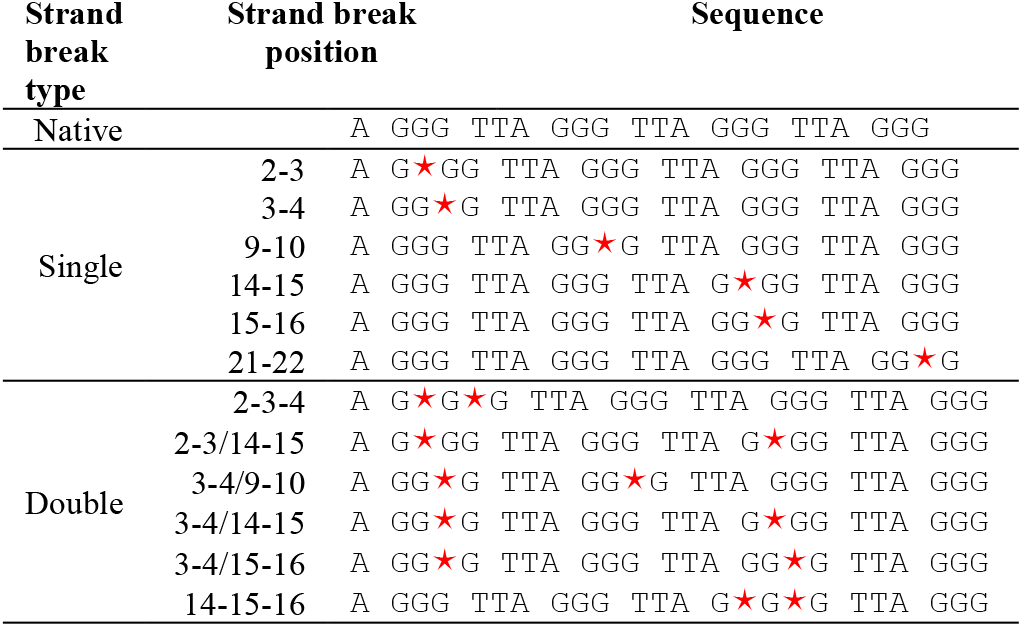
Schematic representation of the positions of the considered strand breaks, represented by the symbol *. CA and NC strand breaks occur on the two O-P(O2)-O sides of the same phosphate group,

The presence of strand breaks inevitably induces some structural variations impacting the G4, which may be more or less marked, highly localized or conversely more global. Monitoring the evolution of the main structural parameters allows to account for the impact of the type of lesion and its position on the specific G4. Namely, we focused on the distance of the centers of mass of the tetrads, their twist angle and the angles formed between the guanines belonging to the same quartet. In addition, the RMSD of the guanines forming the tetrads highlights the conservation of the quartet arrangement and the global preservation of the G4 conformation.

As shown in Electronic Supplementary Information (ESI, Figure S1-S56) and in Figure 3, RMSD clearly highlights the conservation of the quartet arrangement of the G4, and hence the stability of this conformation. This is particularly visible on the 2D-RMSD maps plotted for each structure (Figure 3 and ESI Figure S1-S56). Indeed, the structural deviation of the quartets remains relatively small throughout the simulation, rarely exceeding 3 Å. However, when considering the whole nucleic acid structure, we observe much larger structural variations, even for the native G4. This result is due to the peripheral loops, whose flexibility is clearly enhanced by the presence of strand breaks, and constitutes further evidence of the coexistence of a rigid core with flexible loops in G4 arrangements. Interestingly, as shown in Figure 3 for specific strand break positions, this effect is similar and is produced analogously for both CA and NC structures. Remarkably, the stability of the G4 arrangement is preserved also in presence of multiple strand breaks, differently for what is observed for other lesions, such as abasic sites [55]. This is of great importance since ionizing radiation deposit their energy in a limited spatial area, hence usually leading to DNA cluster lesions [56,57].

**Figure 3.**
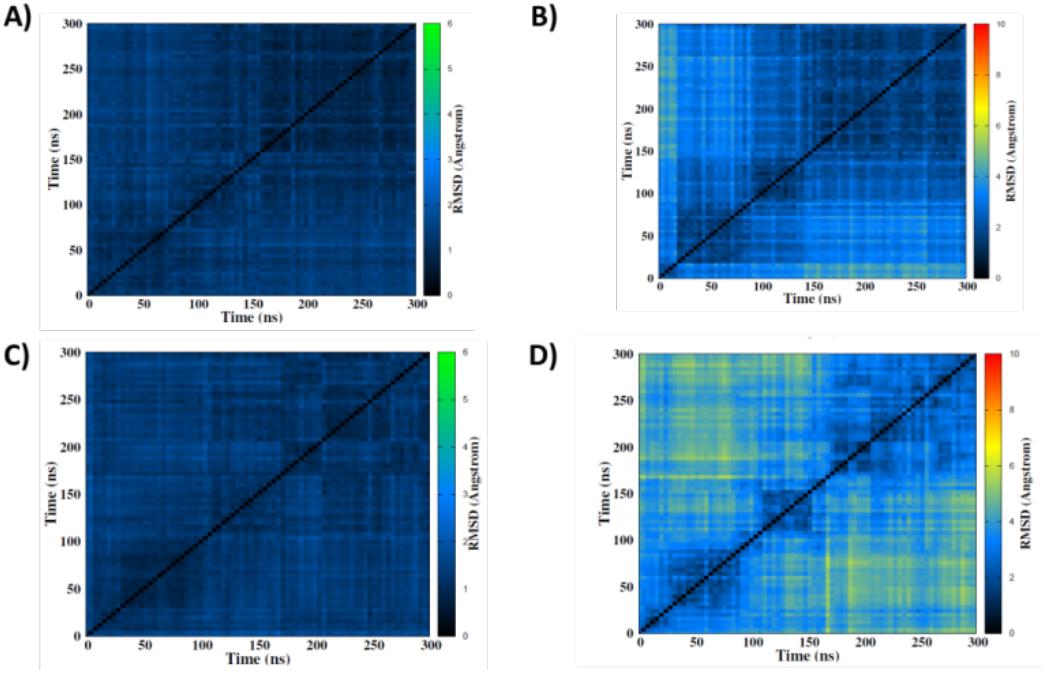
2D-RMSD maps of simulations of the native structure CA (top) and the 14-15-16 NC (bottom), including the tetrads only (A and C) or the whole DNA (B and D).

The behavior of G4 is in striking contrast with the one of canonical double-helical B-DNA structures. Indeed, in the latter case a break in the backbone induces a high instability of the genome, which is mainly due to the dispersion of the broken DNA fragments, especially in presence of double-strand breaks. Conversely, G4s appear to be much more resilient and strand breaks do not alter their arrangement. Once again, this feature can be correlated to the biological role of G4s, and the protective role they can play in presence of high oxidative stress. The global stability of G4s to strand-breaks is also resonating with their resilience to the presence of oxidative damages, which we have recently determined [31]. However, it has to be pointed out that the guanine core may be much more sensitive to the damages. Indeed, the introduction of an abasic site [55] may lead to the G4 disruption or to a complex structural reorganization necessary to maintain its folding.

Our results are also coherent with those reported by Kumari et al. [53] who experimentally demonstrated the resistance of G4s to strand breaks. Firstly, their in vitro experiments highlight the formation of stable intra- and intermolecular G4s after exposure to ionizing radiations. Also, their cell irradiation experiments suggest that G4-forming regions also exhibit high ionizing radiation resistance. Furthermore, their experimental results point to the fact that strand breaks occur mainly into G4-connecting loops. In fact, the introduction of a strand-break lesion into the loops causes them to open. Such a situation undoubtedly leads to a modification of the global arrangement of the G4 DNA. The phenomenon is particularly well highlighted by our 2D-RMSD maps (see ESI, Figure S1-S56). However, as the guanine core is not affected, the structural properties of tetrads remain unchanged. Interestingly, from our simulations we may infer that strand-breaks located into the connecting loops show an even higher structural resistance compared to those directly connecting guanines forming the internal core.

To provide a deeper analysis of the effects of strand breaks formation on G4 structures, we have also considered more local deformations. As a matter of fact, Hoogsteen base-pairing of guanines and the interaction with alkali metal ions are the most crucial factor behind the formation of G4s [58,59], while the involvement of the backbone should be considered minor in dictating their formation.

The time series of the angles formed between the guanines on the tetrads still show a global stability (see ESI, Figure S57 and Table S1). More precisely, the values are all globally centered around the values explored by the native undamaged structure: i.e., ca. 90° for the adjacent and ca. 180° for the opposite guanines, the fluctuations being only indicative of slightly minor changes in their arrangements. If we focus on the respective distance between the center-of-mass of the tetrads, we may also observe only slight fluctuations compared to the native structure, in agreement to the global stability revealed by the RMSD analysis.

Instead, larger fluctuations can be observed for the twist angles (see Table S1 and Figure 4). Indeed, the presence of strand breaks induces a slight enlargement of the distribution of the twist angles, which may be related to an increased flexibility. This is particularly evident for the twist angle between the two terminal tetrads, while the variations of the twists involving the central tetrad are less important. In addition to the enlargement of the distribution, we can notice also some deviations on the value of its maximum which may lead to deviation between 5 and 10° from the undamaged structure, hence pointing to a slightly, albeit non-negligible, structural reorganization. Obviously, this deviation is maximal when the lesion is on the backbone directly connecting two tetrads, while less pronounced when the loops are involved. Finally, and despite the global stability observed and discussed, we should point out that only one trajectory makes a significant exception. This involves one replica for the CA damaged at the 14-15 position (Figure S10). In this case we can observe a rearrangement of the first peripheral quartet which is accompanied by a leakage of a K^+^ cation, leading to the expulsion of the guanine G8 into a loop region and further the complete destabilization of the peripheral tetrad. In the past, we have already observed that the loss of the cation is an important phenomenon in the destabilization of the G4 structure [31,55]. Although it points out the crucial role played by the metal cations in dictating the stability of the G4 arrangements, this case remains isolated since it is only observed once in all our simulations and can then be considered as a rare event.

**Figure 4.**
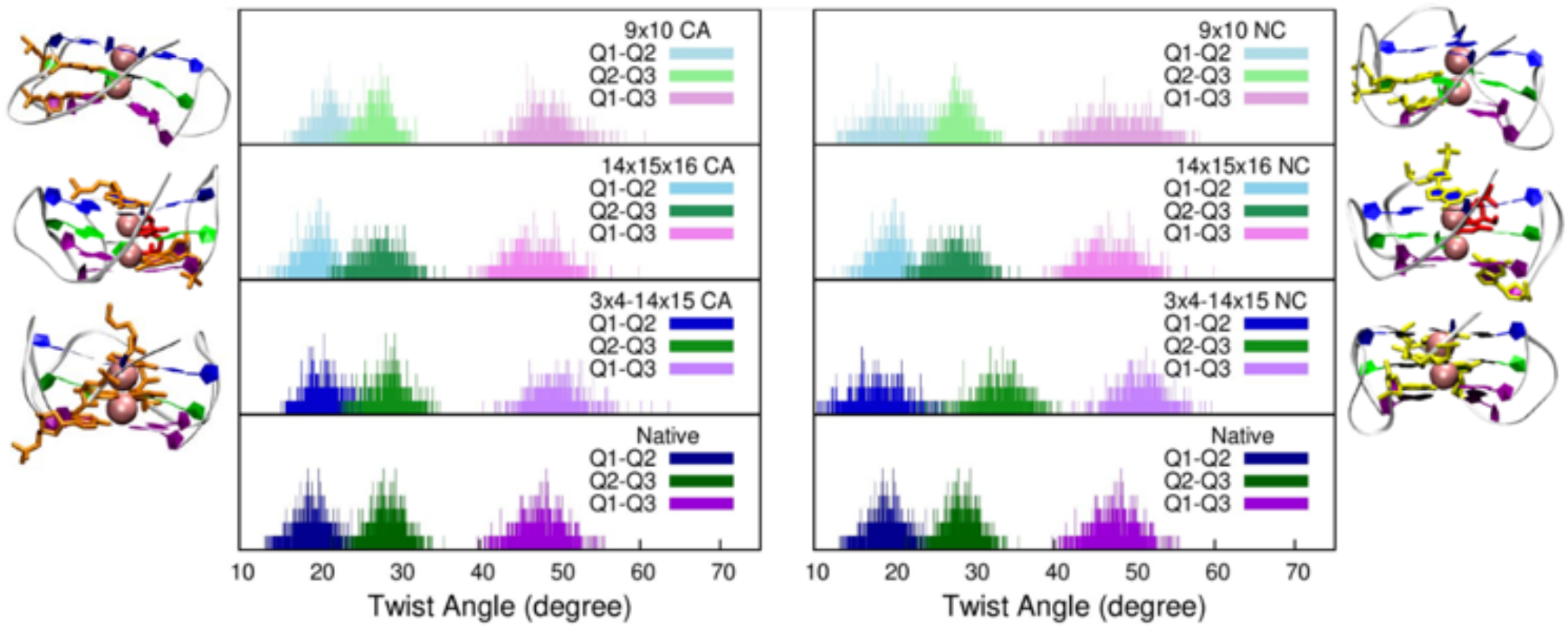
Distribution of the twist angle of representative DNA damaged structures, compared to those of the native structure, shows a shift in the values of this angle

## Conclusion

Strand breaks are important and toxic classes of DNA lesions, mostly produced by exposure to ionizing radiations. Double-breaks in a DNA strand are associated to very high cytotoxicity and are difficult to repair. In this contribution, we aim to rationalize the impact of strand breaks on the stability and persistence of intra-strand G4 architectures, considering the important regulatory role played by G4-DNA structures at the cellular level. Although double-helical B-DNA strand-breaks are usually correlated to a strong structural destabilization and consequent genome dispersion, our results have consistently shown that G4s experience only negligible structural deformations in 26 out of 27 cases of the considered strand breaks, and maintain their global folding and shape. The introduction of a strand break is typically accompanied by a slight increase of flexibility of the connecting loops, and by a slight change of the twist angle, especially when the break is directly located between guanines belonging to different tetrads. Only one strand-break occurrence, i.e. when the strand is broken between the 14^th^ and 15^th^ nucleotides, has led to unfolding of the G4 in the second replica. This is due to the deformation of one tetrad and the subsequent expulsion of the stabilizing K^+^ cation, and can be considered as a rare event. Our results, which are coherent with the report of Kumari et al [40], confirm the high stability of the G4 and their inherent resistance to strand-break damages. From a molecular point of view this can be attributed to the combined effect of stabilizing intra-tetrad Hoogsteen H-bonds (conferring rigidity to the tetrad core) with inter-tetrad π-stacking interactions. From a cellular point of view, this occurrence can be related to the protective and regulatory role played by G4s in regulating gene expression and cellular senescence. As already noticed in presence of oxidative lesions, preserving a G4 arrangement in presence of high stress conditions has the effect of limiting the expression of oncogenes, inhibiting the telomerase and avoiding the emergence of an immortal phenotype, which could lead to carcinogenesis.

It is important to point out that our MD simulations consistently used a pre-folded G4 on top of which strand breaks have been created. Hence, our results are unequivocal concerning the stability of the quadruplexes. However, it cannot be inferred whether the presence of strand breaks could lead to the inhibition of the folding itself, hence limiting the actual G4 presence in cells. The propensity to G4 folding of damaged DNA, which could also strongly depend on the length of the fragments, and hence on the radiation intensity, will require resorting to enhanced sampling procedures and will be addressed in forthcoming contributions.

## Computational Section

### Force field for non-standard nucleotides

Prior to modelling the formation of strand breaks, specific force field parameters for the two sides of the cleaved backbone needed to be generated. To this aim we chose the AMBER ff99bsc1 force field [60,61] [45] and specific modifications to the guanine and adenine force field have been performed to obtain CA and NC ends (see ESI). The geometry of each of the new residues has been optimized at the B3LYP/6-311+G(d,p) level of theory, with Gaussian 09 [62]. Restrained electrostatic potential (RESP) charges have been obtained at HF/6-31G* and converted into amber format with the antechamber utilities.

### Molecular dynamics simulations

ll the setups have been generated using the AMBER16 suite of programs [63]. The initial G4 structure of h-telo was obtained from the pdb database (PDB:1KF1) [64], and the strand breaks were manually created at specific sequence positions (see Table 1 and ESI). Then the initial systems were solvated in an octahedral TIP3P water box [65] with a 12 Å buffer; and electroneutrality was provided by the addition of K^+^ ions. Note that the central K^+^ ions present in the crystal structure have been kept. Hydrogen mass repartitioning [66] was applied to allow the integration of the Newton equations of motion using a 4 fs time step in combination with the RATTLE and SHAKE algorithms [67]. All the MD simulations were performed with the NAMD software [68,69] until reaching a simulation time of 300ns in the NPT ensemble maintained with a Langevin thermostat and barostat [70]. Each of the simulation was preceded by 1000 minimization steps and by 36 ns of equilibration. All simulations have been performed on two independent replicas to increase the global sampling. VMD [71] was used to visualize and to analyze the MD trajectories. G-quadruplex structural parameters have been calculated using the script developed by Tsvetkov et al. [72].

## Acknowledgements

Support from the Universities of Lorraine, Palermo and Paris, as well as from the French CNRS are gratefully acknowledged. T.M. thanks Universita degli Studi di Palermo for funding his doctoral program. EB thanks the French “Ministère de l’Education Supérieure, de la Recherche Scientifique et de l’Innovation” (MESRI) for funding her postdoctoral fellowship through the special GAVO program. AM thanks ANR (Agence Nationale de la Recherche) and CGI (Commissariat à l’Investissement d’Avenir) for their financial support of this work through Labex SEAM (Science and Engineering for Advanced Materials and devices) ANR 11 LABX 086, ANR 11 IDEX 05 02. The support of the IdEx “Université Paris 2019” ANR-18-IDEX-0001 and of the Platform PSM3B is gratefully acknowledged.

## Entry for the Table of Contents

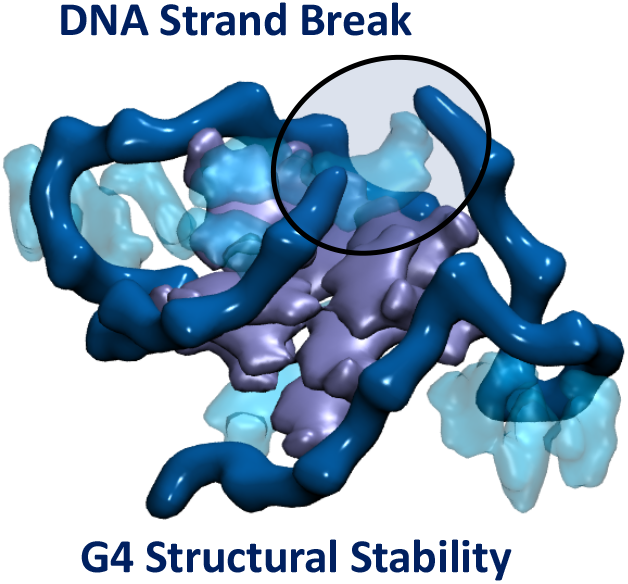

By using molecular long-range molecular dynamic and simulations we have shown that DNA G-quadruplexes show a remarkable stability to the presence of strand break. Indeed, the G-quadruplex structure is maintained and only some flexibility of the peripheral loops can be underlined. These results may also be correlated to the protective role of G-quadruplexes.

Institute and/or researcher Twitter usernames: @AntonioMonari, @baronegiampaolo, @Al3XI0, @DrEmmaBignon

