## Supplementary Information for "Never cared for what they do. High structural stability of Guanine-quadruplexes in presence of strand-break damages"

Electronic Supplementary Information for

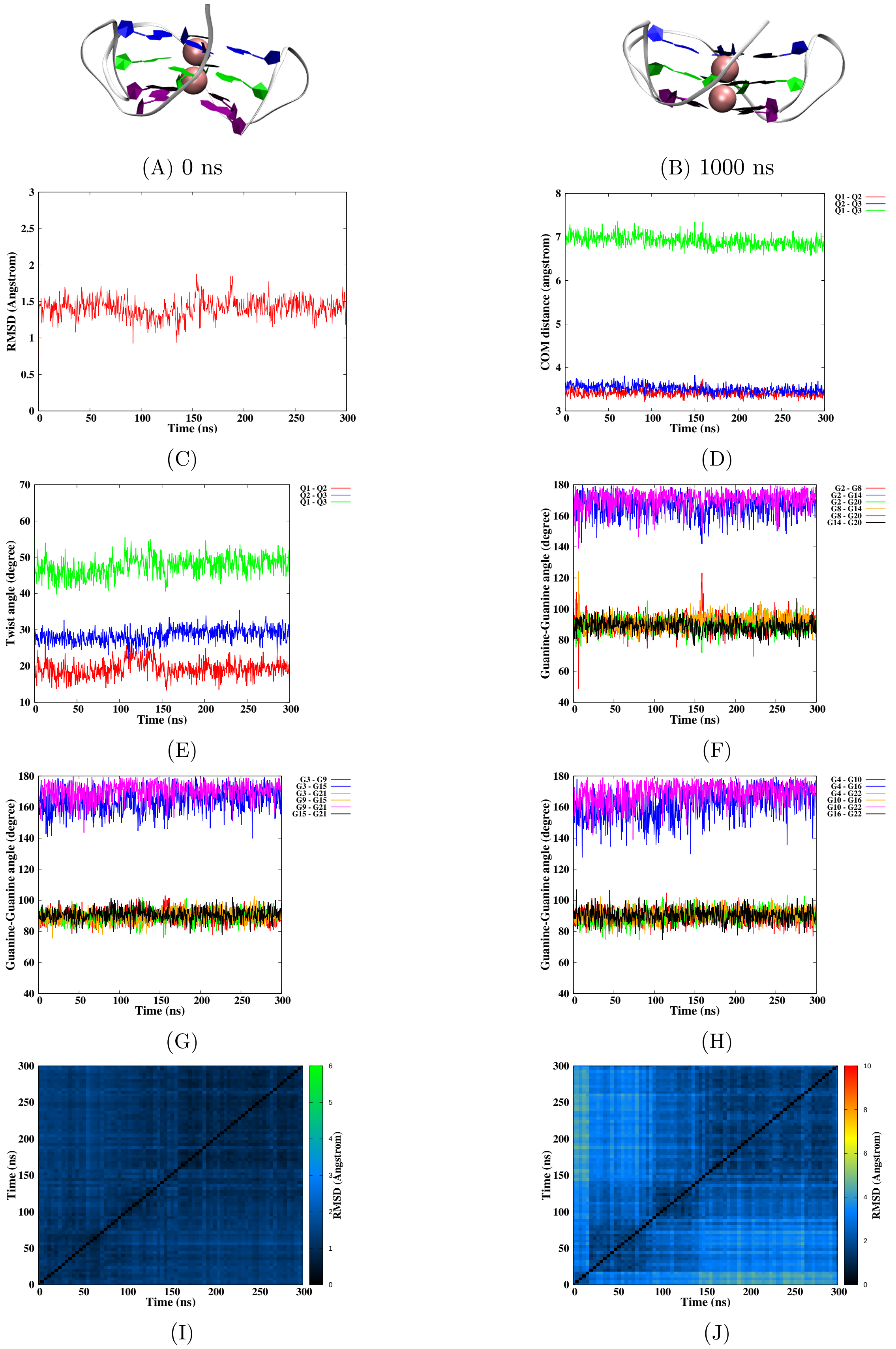


Figure S1 – Simulation of the native G-quadruplex DNA, run 1. Snapshots of the initial and final MD simulation (A-B). RMSD of guanines forming the tetrads (C), distance between the tetrads (D), twist angle (E), angle between guanines for the first (F), second (G) and the third (H) tetrad, 2D-RMSD of guanines forming the tetrads (I) and the whole DNA (J).

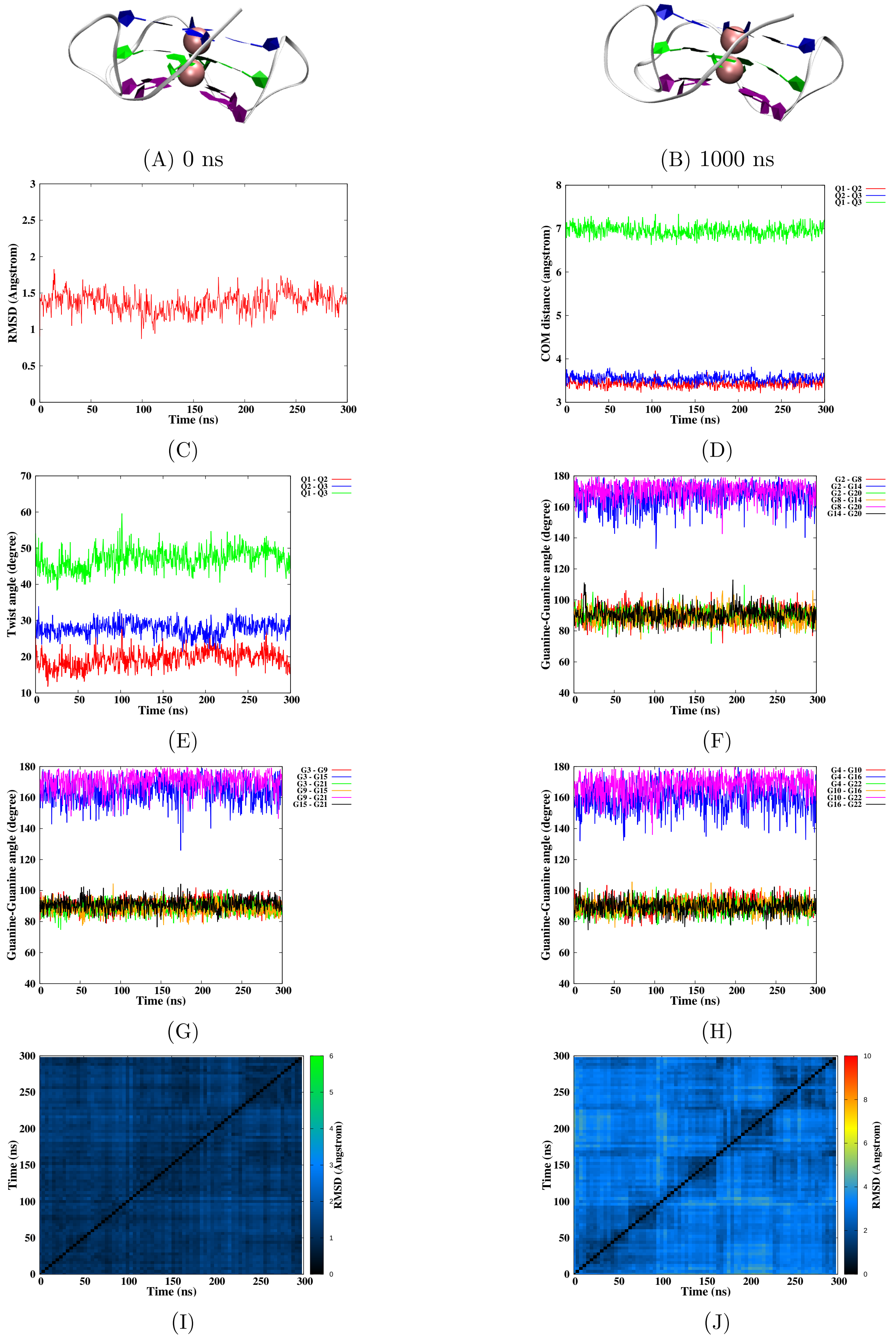


Figure S2 – Simulation of the native G-quadruplex DNA, run 2. Snapshots of the initial and final MD simulation (A-B). RMSD of guanines forming the tetrads (C), distance between the tetrads (D), twist angle (E), angle between guanines for the first (F), second (G) and the third (H) tetrad, 2D-RMSD of guanines forming the tetrads (I) and the whole DNA (J).

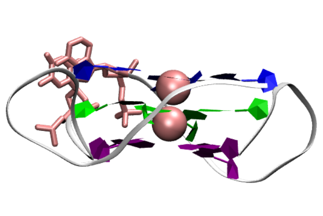

(A) 0 ns

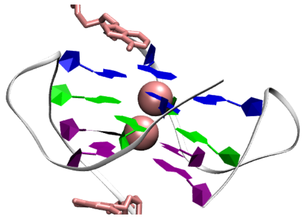

(B) 1000 ns

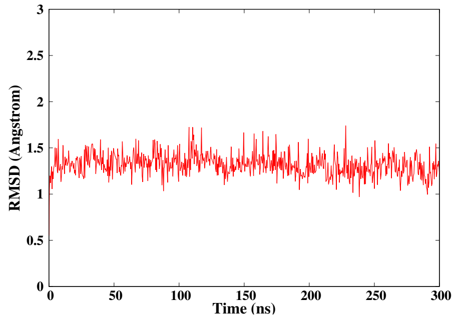


(C)

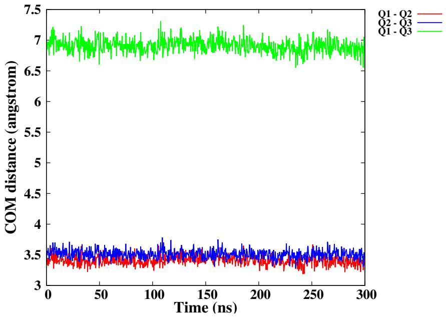


(D)

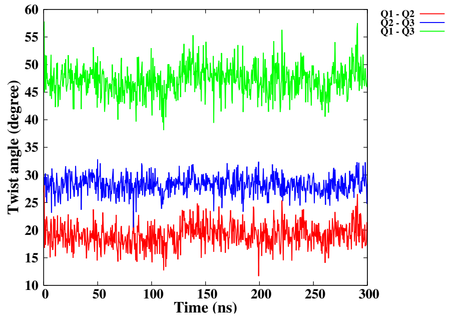


(E)

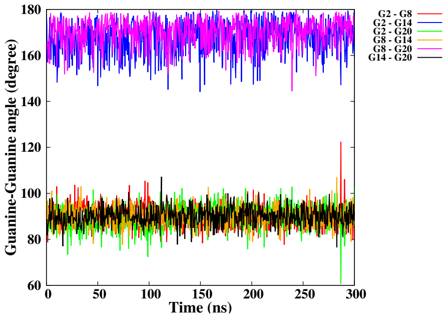


(F)

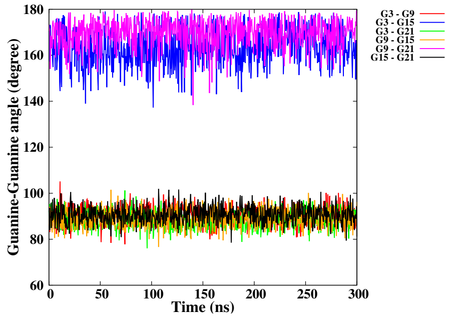


(G)

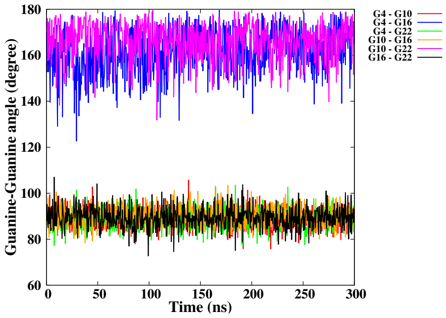


(H)

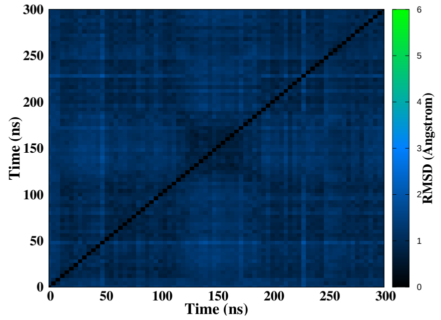


(I)

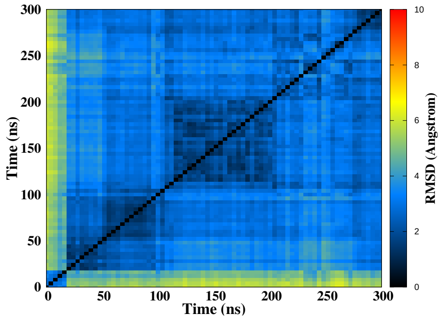


(J)

Figure S3 – Simulation of the CA damage at position 12-13 **(in loop)**, run 1. Snapshots of the initial and final MD simulation (A-B). RMSD of guanines forming the tetrads (C), distance between the tetrads (D), twist angle (E), angle between guanines for the first (F), second (G) and the third (H) tetrad, 2D-RMSD of guanines forming the tetrads (I) and the whole DNA (J).

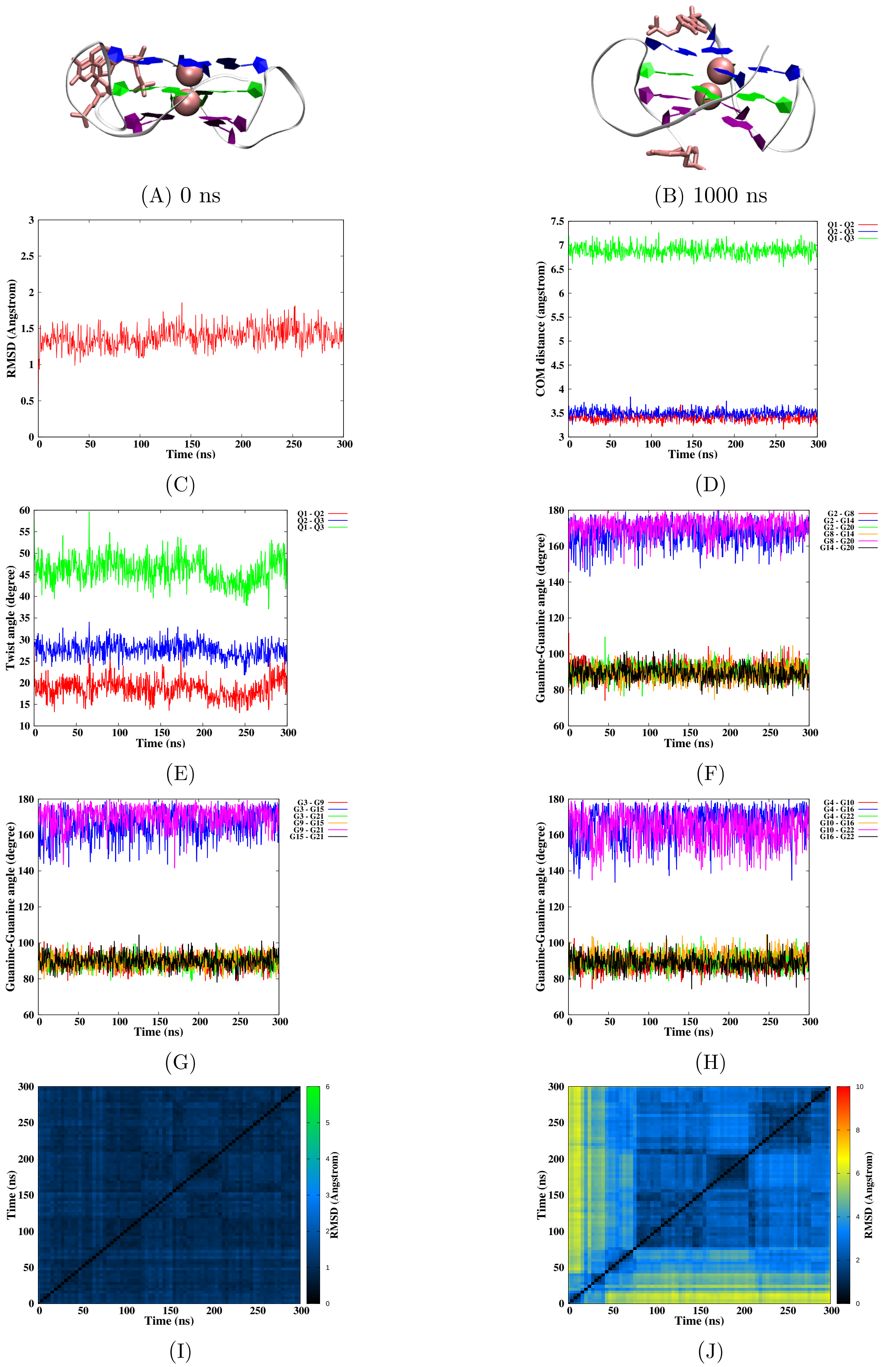


Figure S4 – Simulation of the CA damage at position 12-13 (in loop), run 2. Snapshots of the initial and final MD simulation (A-B). RMSD of guanines forming the tetrads (C), distance between the tetrads (D), twist angle (E), angle between guanines for the first (F), second (G) and the third (H) tetrad, 2D-RMSD of guanines forming the tetrads (I) and the whole DNA (J).

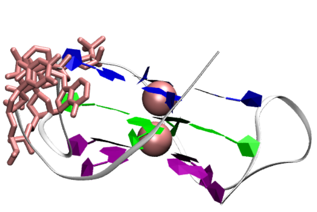

(A) 0 ns

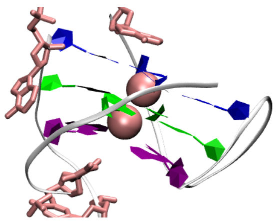

(B) 1000 ns

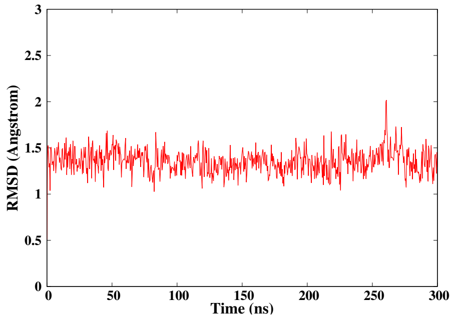


(C)

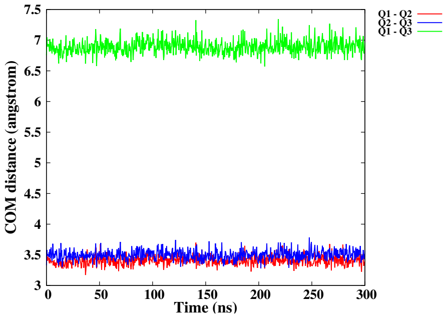


(D)

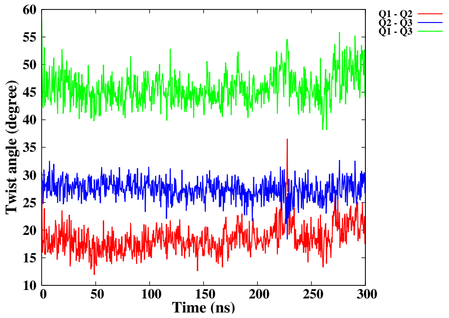


(E)

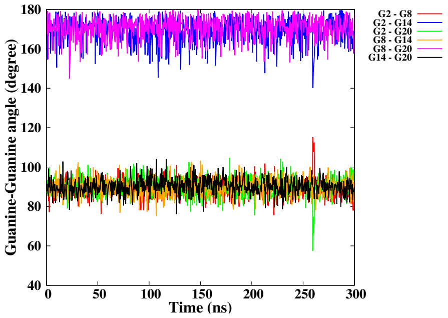


(F)

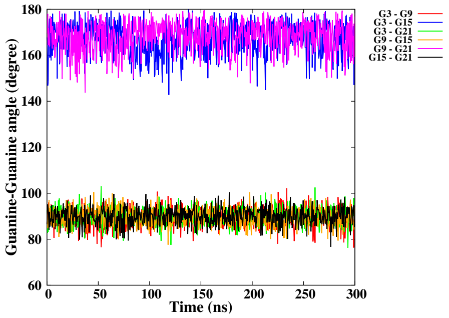


(G)

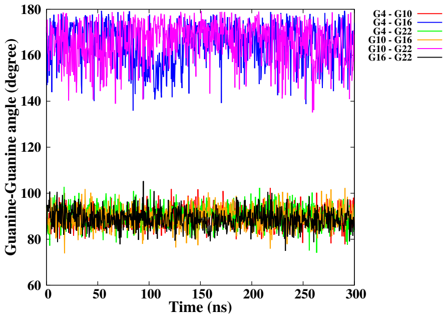


(H)

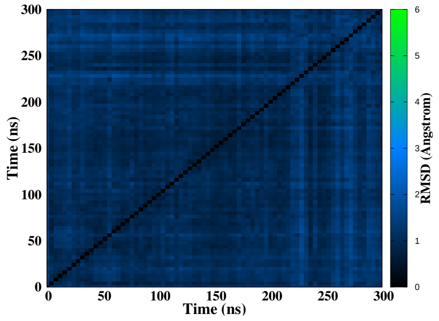


(I)

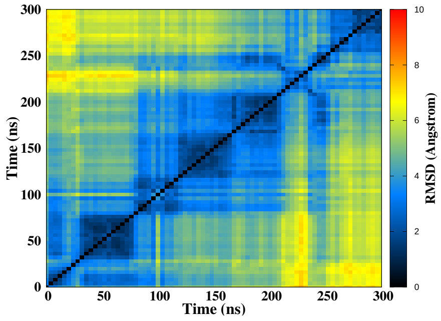


(J)

Figure S5 – Simulation of the CA damage at position 6-7/12-13 (in loop), run 1. Snapshots of the initial and final MD simulation (A-B). RMSD of guanines forming the tetrads (C), distance between the tetrads (D), twist angle (E), angle between guanines for the first (F), second (G) and the third (H) tetrad, 2D-RMSD of guanines forming the tetrads (I) and the whole DNA (J).

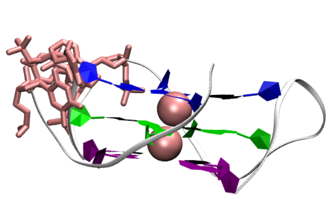

(A) 0 ns

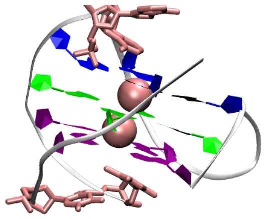

(B) 1000 ns

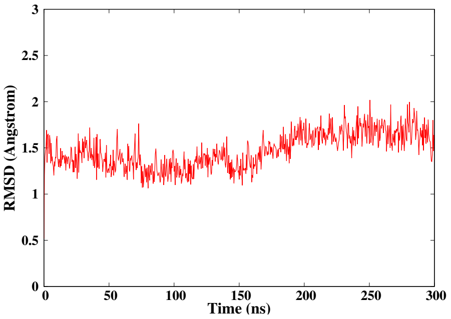


(C)

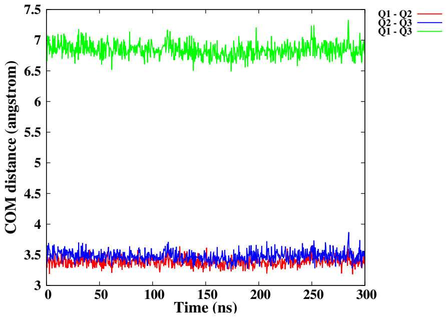


(D)

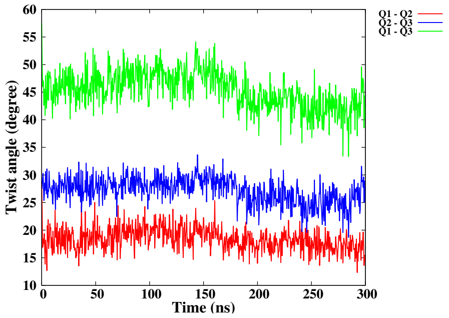


(E)

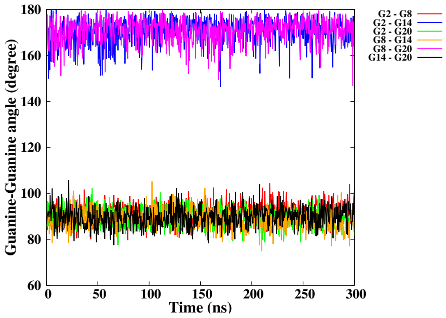


(F)

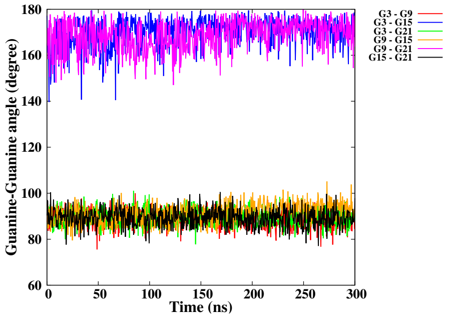


(G)

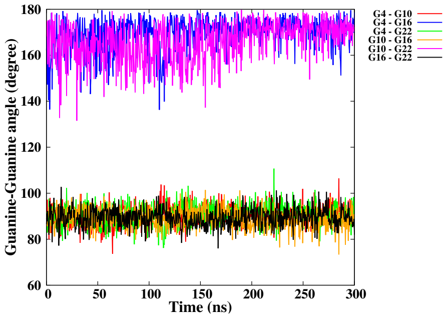


(H)

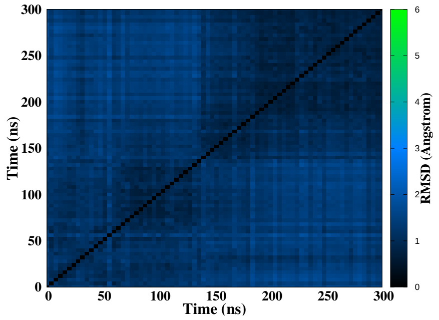


(I)

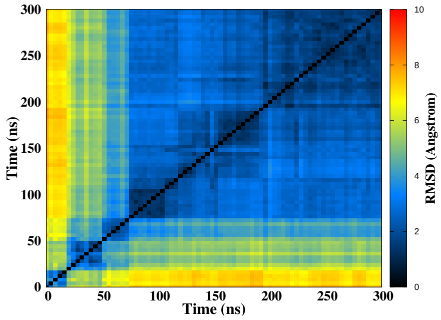


(J)

Figure S6 – Simulation of the CA damage at position 6-7/12-13 (in loop), run 2. Snapshots of the initial and final MD simulation (A-B). RMSD of guanines forming the tetrads (C), distance between the tetrads (D), twist angle (E), angle between guanines for the first (F), second (G) and the third (H) tetrad, 2D-RMSD of guanines forming the tetrads (I) and the whole DNA (J).

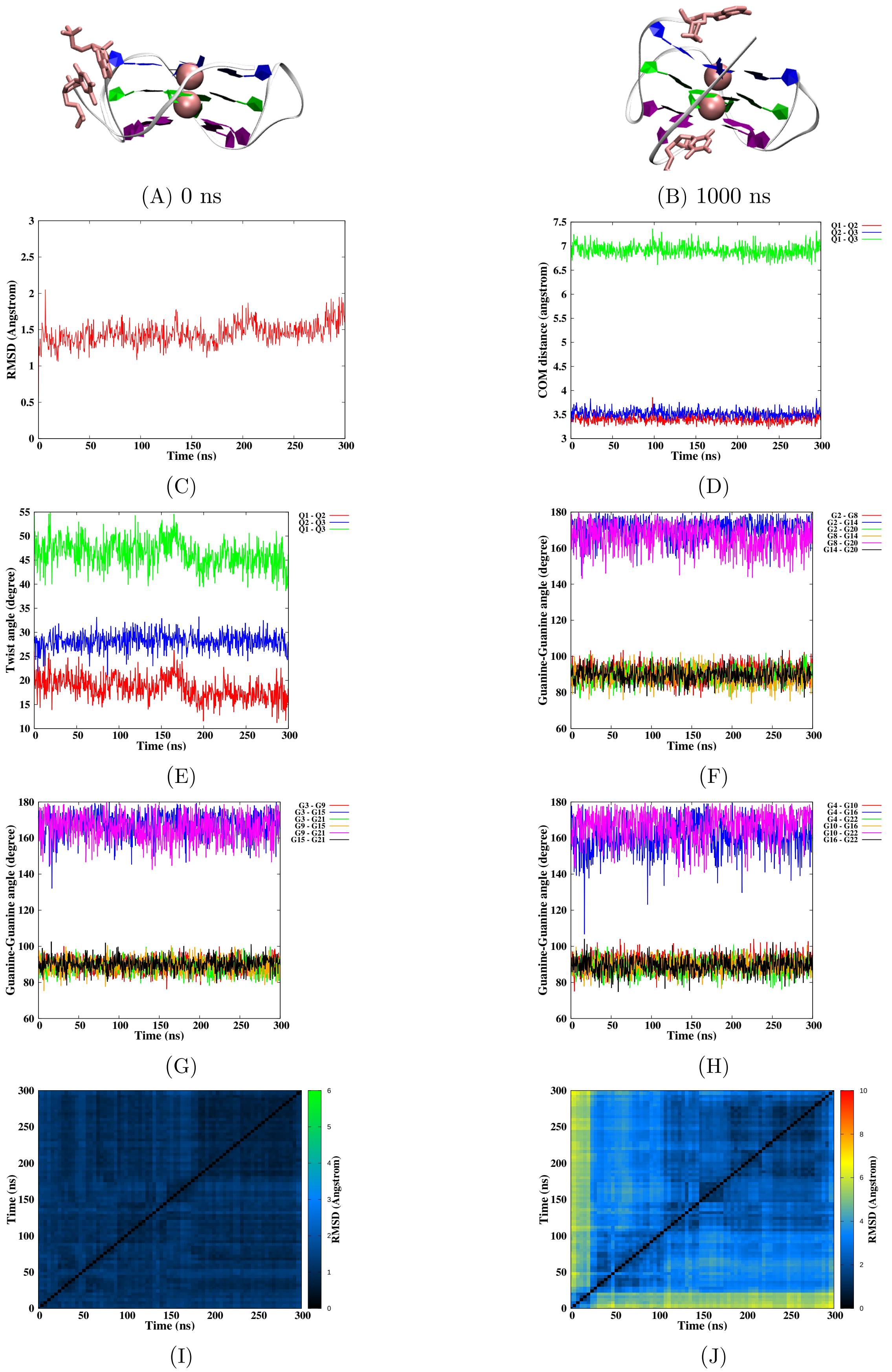

Figure S7 – Simulation of the CA damage at position 6-7 (in loop), run 1. Snapshots of the initial and final MD simulation (A-B). RMSD of guanines forming the tetrads (C), distance between the tetrads (D), twist angle (E), angle between guanines for the first (F), second (G) and the third (H) tetrad, 2D-RMSD of guanines forming the tetrads (I) and the whole DNA (J).

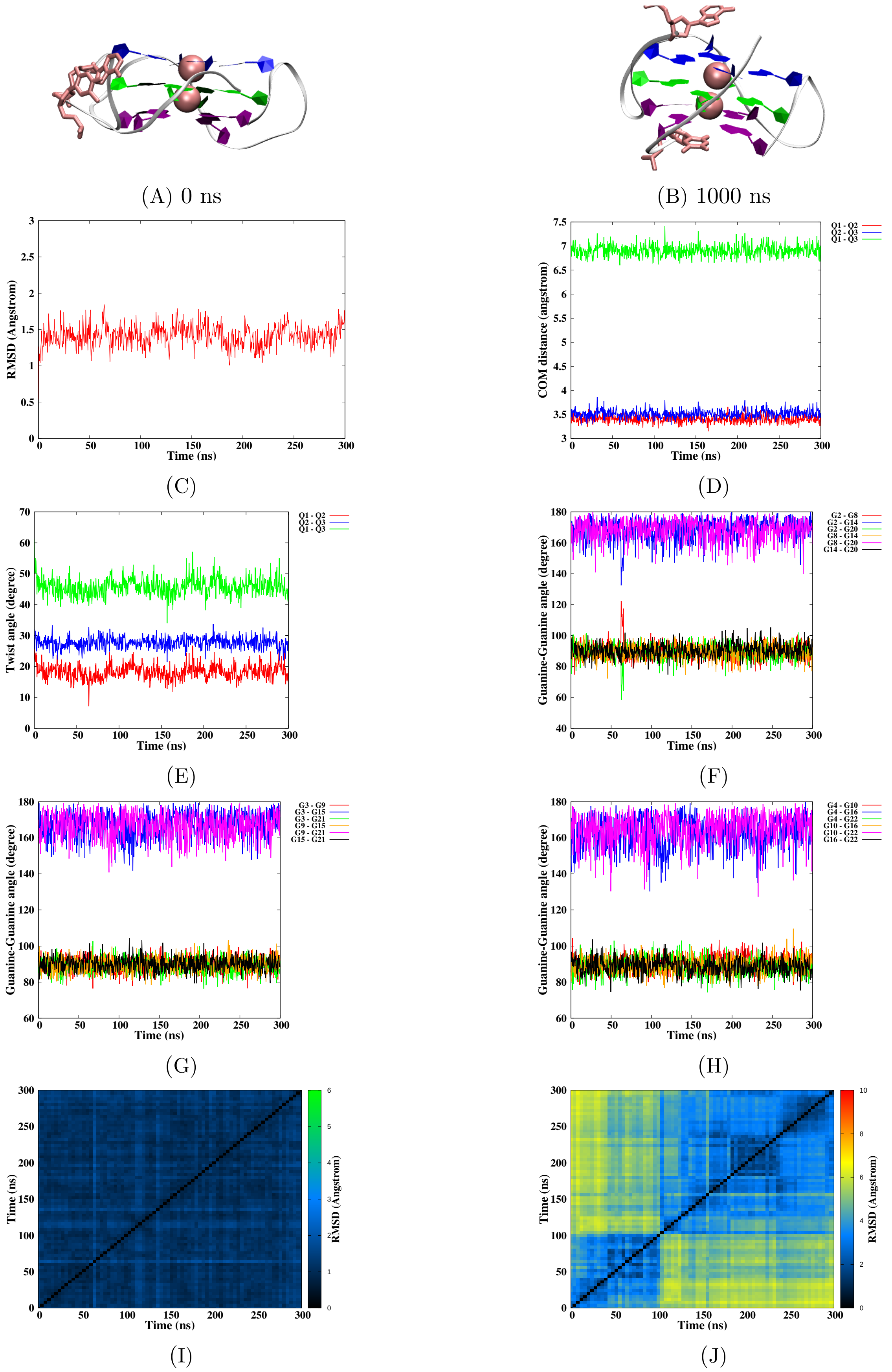


Figure S8 – Simulation of the CA damage at position 6-7 (in loop), run 2. Snapshots of the initial and final MD simulation (A-B). RMSD of guanines forming the tetrads (C), distance between the tetrads (D), twist angle (E), angle between guanines for the first (F), second (G) and the third (H) tetrad, 2D-RMSD of guanines forming the tetrads (I) and the whole DNA (J).

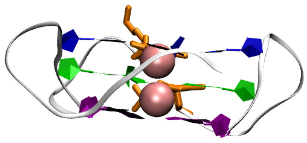

(A) 0 ns

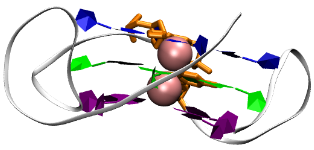

(B) 1000 ns

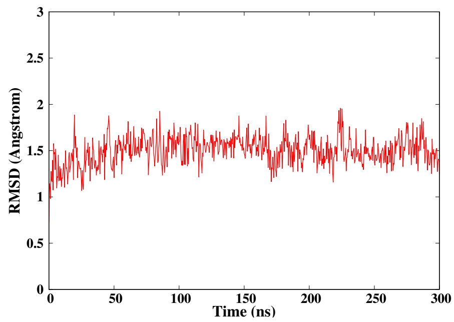


(C)

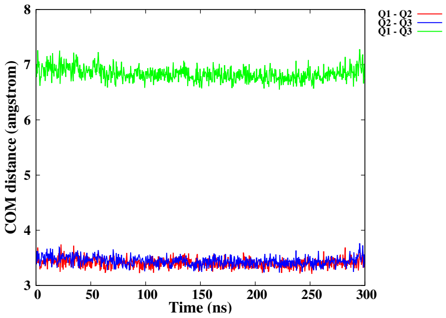


(D)

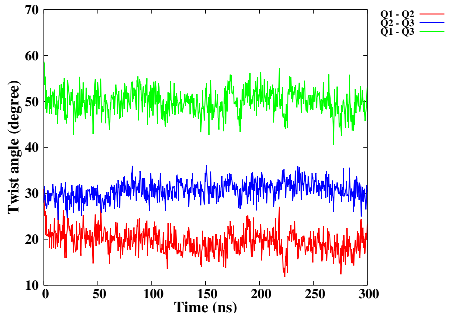


(E)

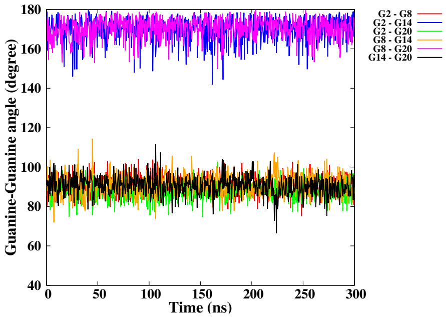


(F)

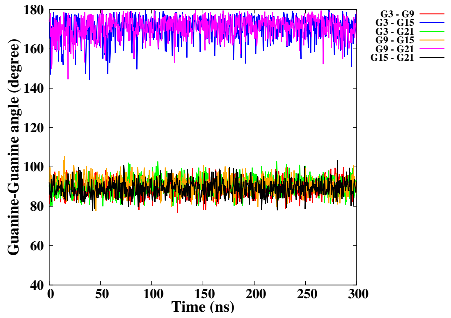


(G)

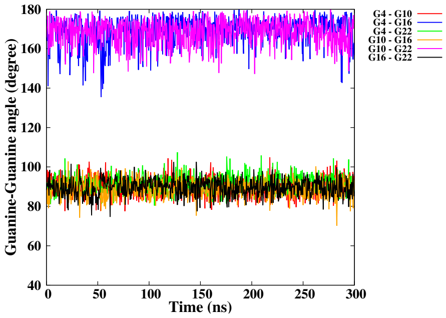


(H)

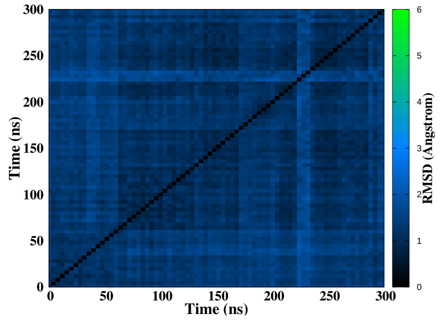


(I)

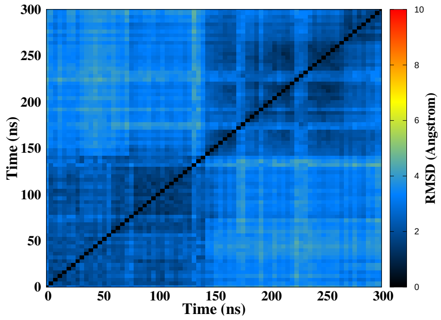


(J)

Figure S9 – Simulation of the CA damage at position 14-15, run 1. Snapshots of the initial and final MD simulation (A-B). RMSD of guanines forming the tetrads (C), distance between the tetrads (D), twist angle (E), angle between guanines for the first (F), second (G) and the third (H) tetrad, 2D-RMSD of guanines forming the tetrads (I) and the whole DNA (J).

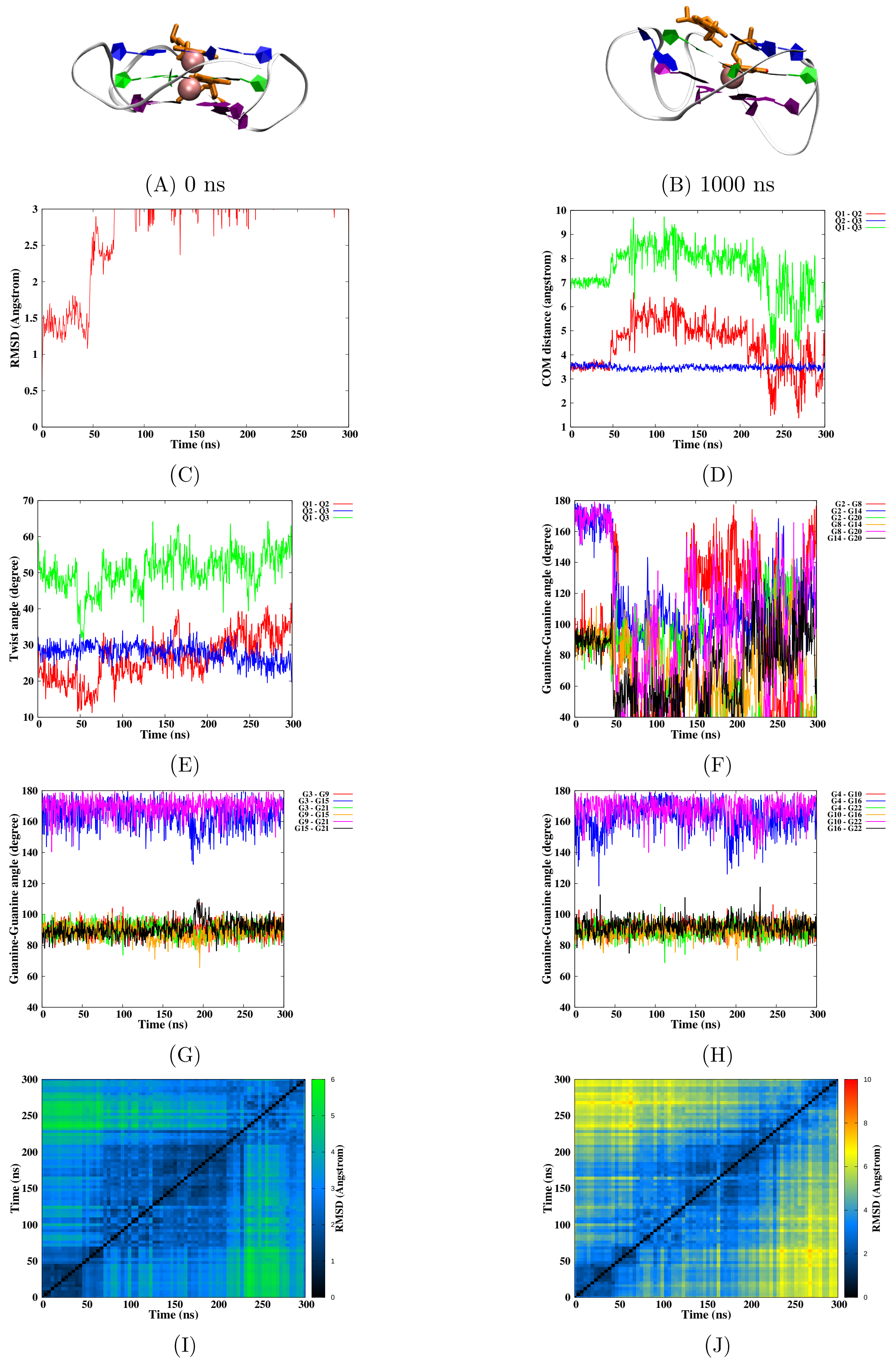


Figure S10 – Simulation of the CA damage at position 14-15, run 2. Snapshots of the initial and final MD simulation (A-B). RMSD of guanines forming the tetrads (C), distance between the tetrads (D), twist angle (E), angle between guanines for the first (F), second (G) and the third (H) tetrad, 2D-RMSD of guanines forming the tetrads (I) and the whole DNA (J).

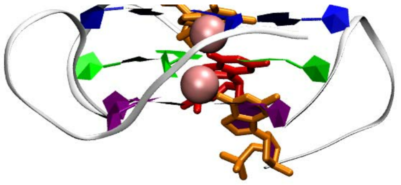

(A) 0 ns

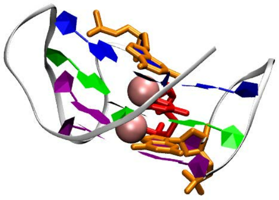

(B) 1000 ns

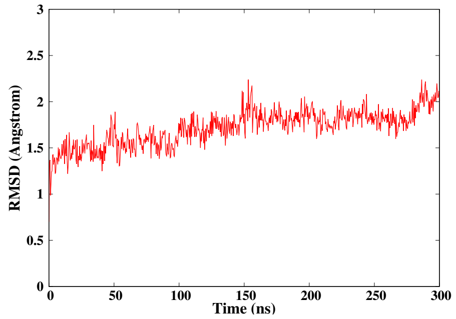


(C)

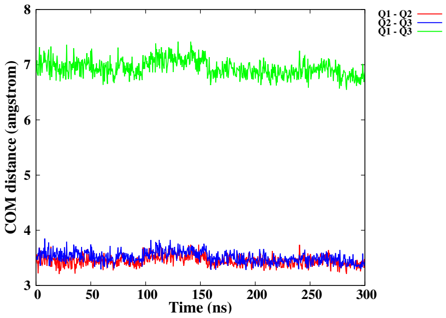


(D)

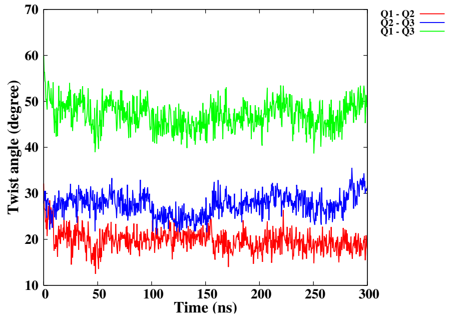


(E)

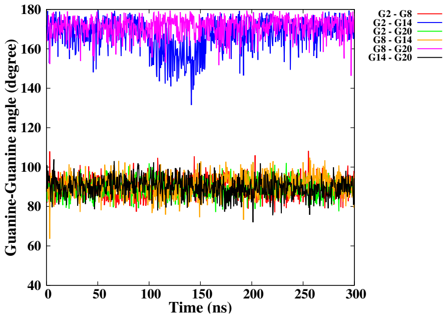


(F)

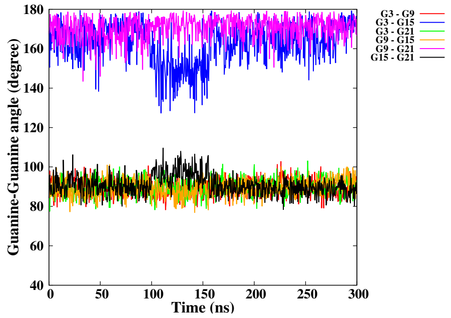


(G)

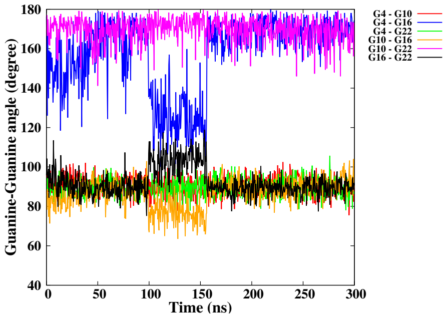


(H)

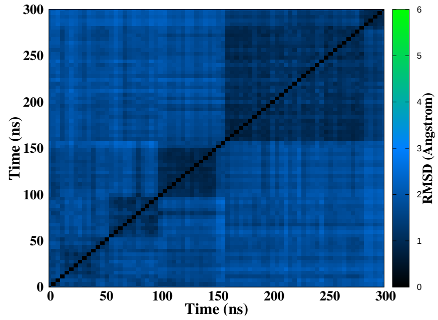


(I)

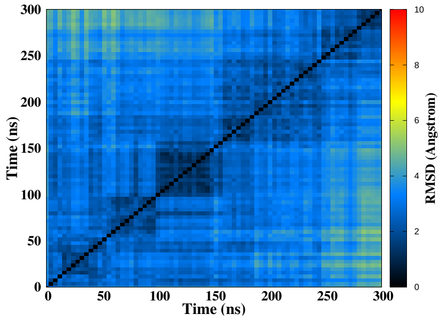


(J)

Figure S11 – Simulation of the CA damage at position 14-15-16, run 1. Snapshots of the initial and final MD simulation (A-B). RMSD of guanines forming the tetrads (C), distance between the tetrads (D), twist angle (E), angle between guanines for the first (F), second (G) and the third (H) tetrad, 2D-RMSD of guanines forming the tetrads (I) and the whole DNA (J).

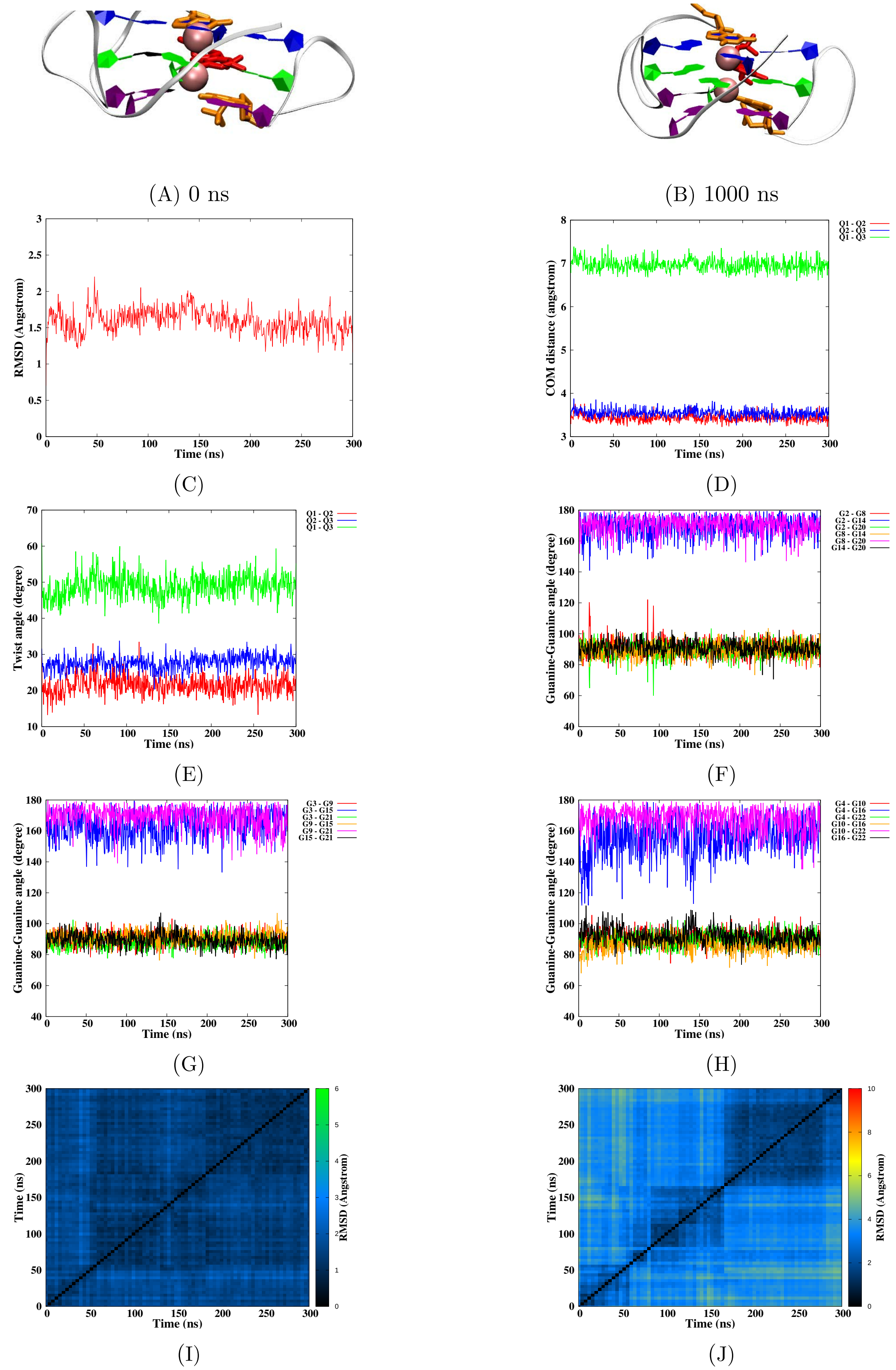


Figure S12 – Simulation of the CA damage at position 14-15-16, run 2. Snapshots of the initial and final MD simulation (A-B). RMSD of guanines forming the tetrads (C), distance between the tetrads (D), twist angle (E), angle between guanines for the first (F), second (G) and the third (H) tetrad, 2D-RMSD of guanines forming the tetrads (I) and the whole DNA (J).

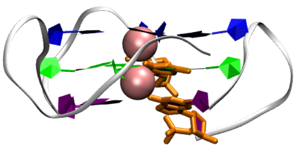

(A) 0 ns

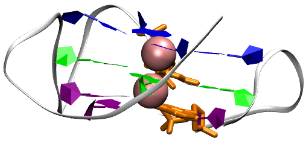

(B) 1000 ns

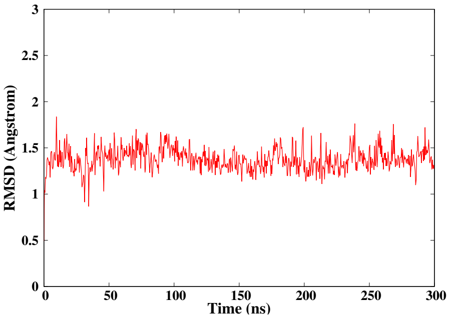


(C)

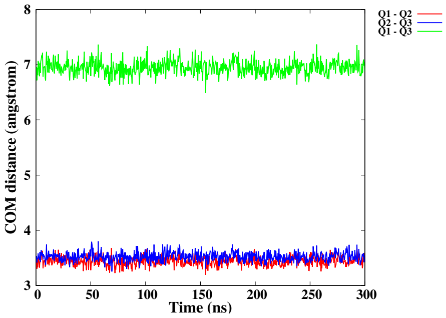


(D)

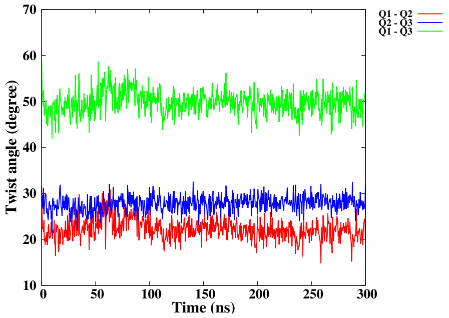


(E)

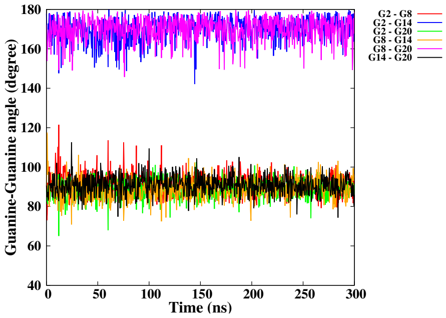


(F)

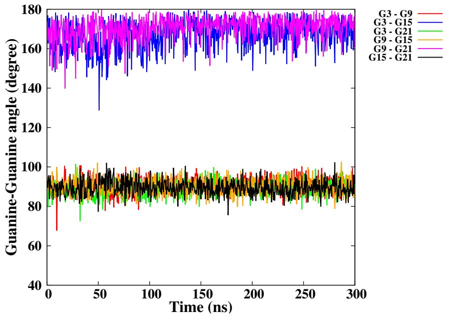


(G)

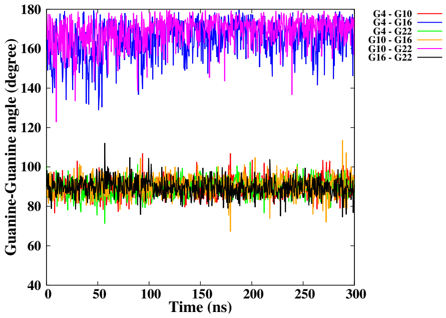


(H)

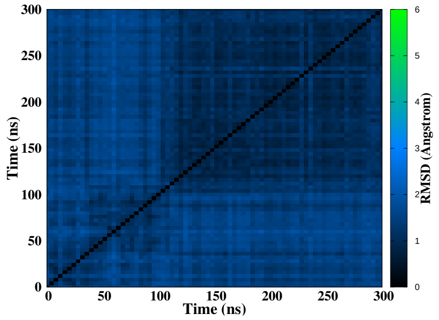


(I)

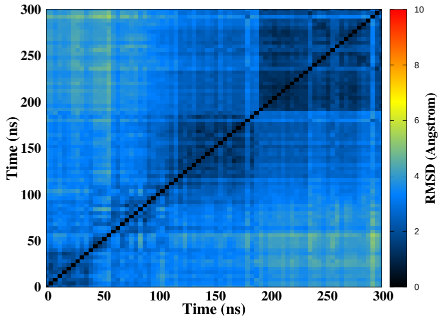


(J)

Figure S13 – Simulation of the CA damage at position 15-16, run 1. Snapshots of the initial and final MD simulation (A-B). RMSD of guanines forming the tetrads (C), distance between the tetrads (D), twist angle (E), angle between guanines for the first (F), second (G) and the third (H) tetrad, 2D-RMSD of guanines forming the tetrads (I) and the whole DNA (J).

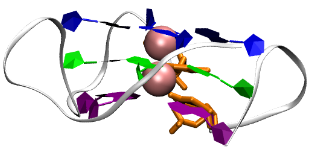

(A) 0 ns

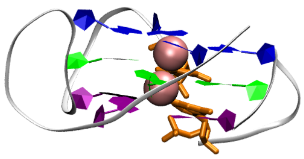

(B) 1000 ns

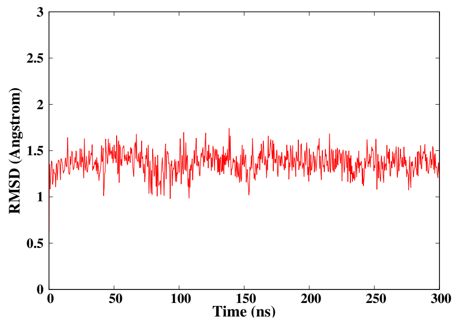


(C)

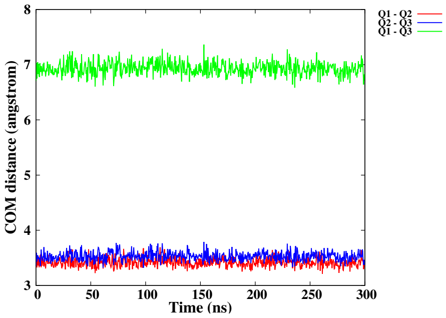


(D)

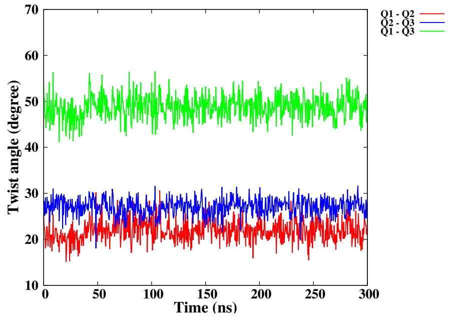


(E)

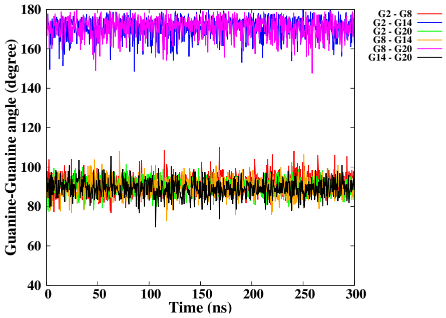


(F)

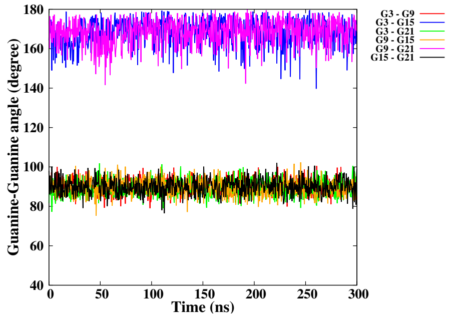


(G)

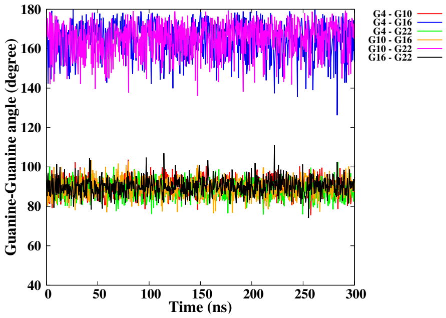


(H)

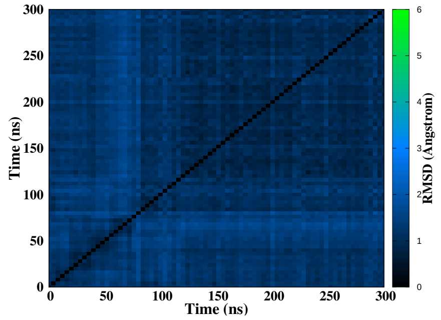


(I)

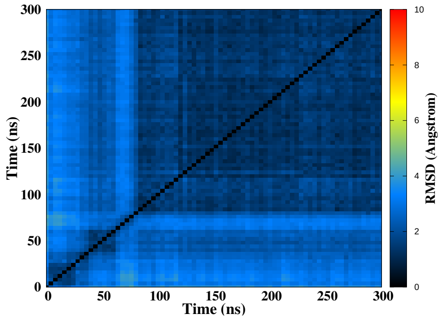


(J)

Figure S14 – Simulation of the CA damage at position 15-16, run 2. Snapshots of the initial and final MD simulation (A-B). RMSD of guanines forming the tetrads (C), distance between the tetrads (D), twist angle (E), angle between guanines for the first (F), second (G) and the third (H) tetrad, 2D-RMSD of guanines forming the tetrads (I) and the whole DNA (J).

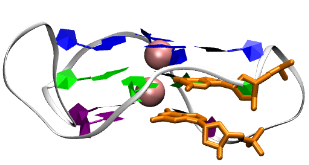

(A) 0 ns

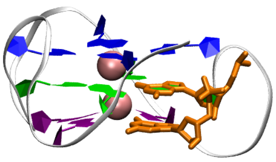

(B) 1000 ns

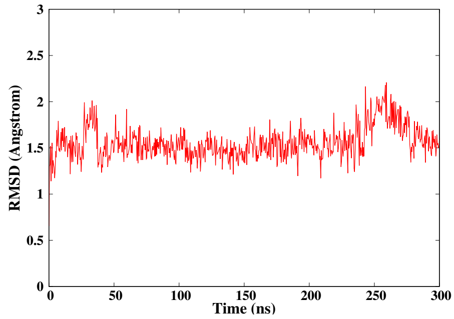


(C)

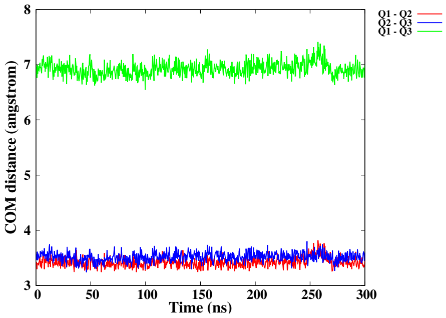


(D)

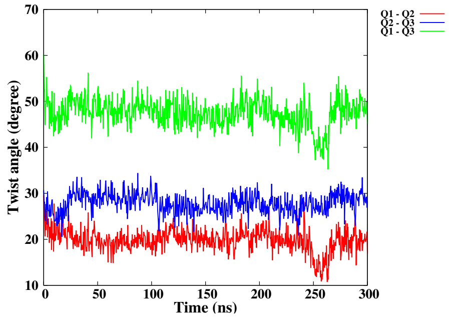


(E)

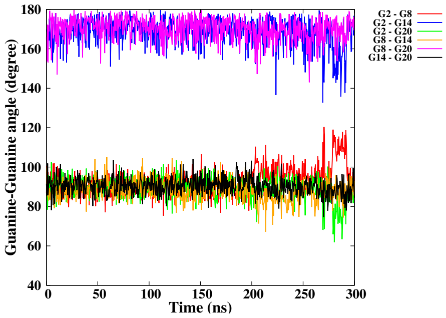


(F)

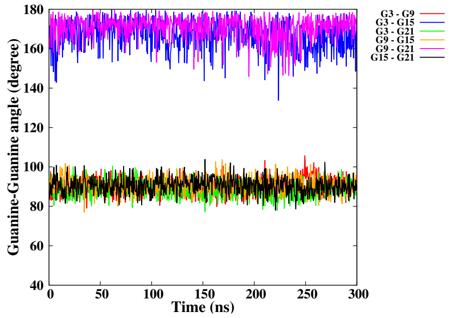


(G)

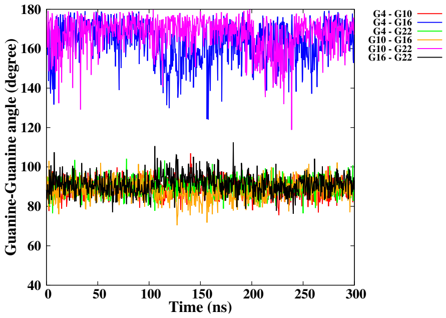


(H)

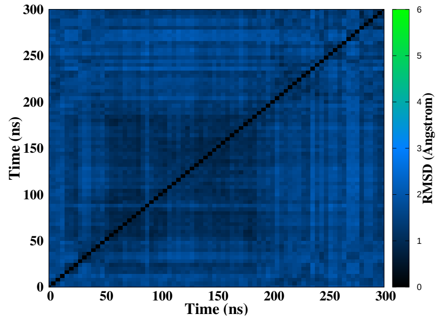


(I)

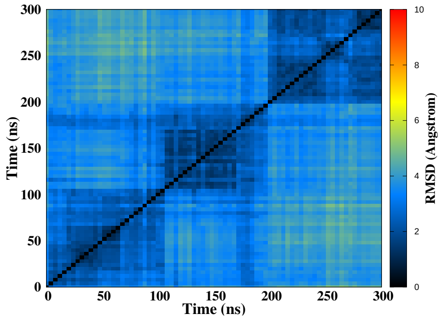


(J)

Figure S15 – Simulation of the CA damage at position 21-22, run 1. Snapshots of the initial and final MD simulation (A-B). RMSD of guanines forming the tetrads (C), distance between the tetrads (D), twist angle (E), angle between guanines for the first (F), second (G) and the third (H) tetrad, 2D-RMSD of guanines forming the tetrads (I) and the whole DNA (J).

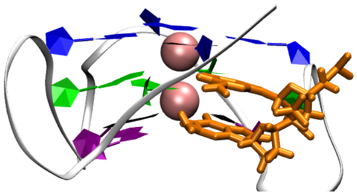

(A) 0 ns

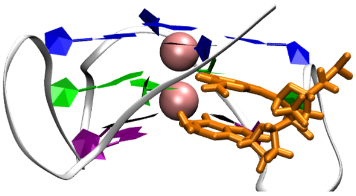

(B) 1000 ns

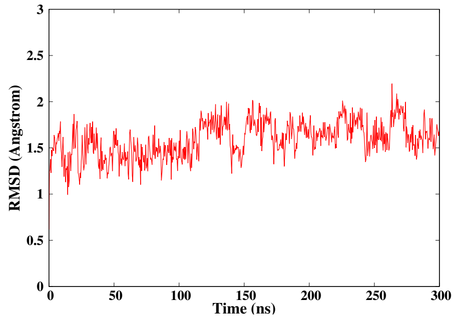


(C)

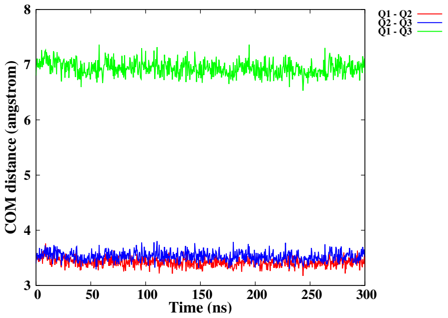


(D)

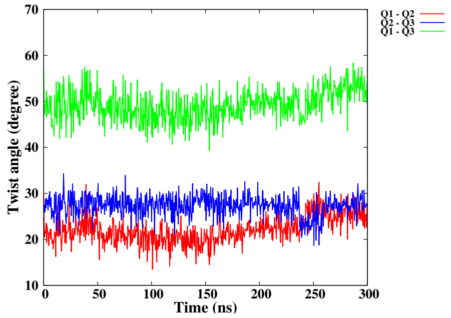


(E)

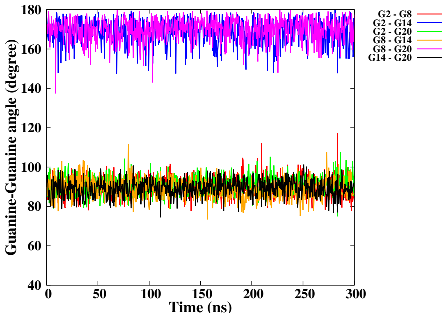


(F)

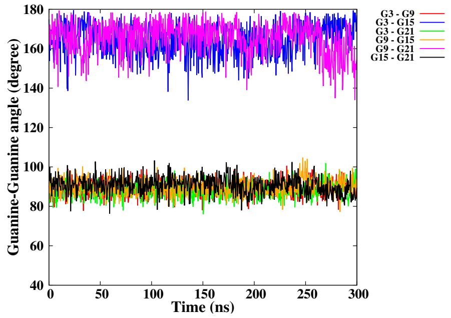


(G)

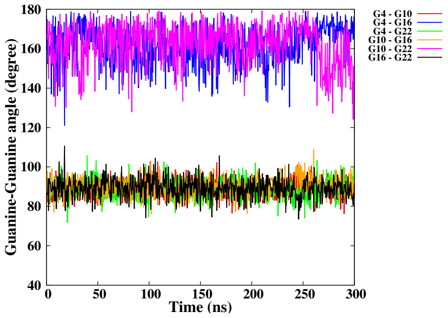


(H)

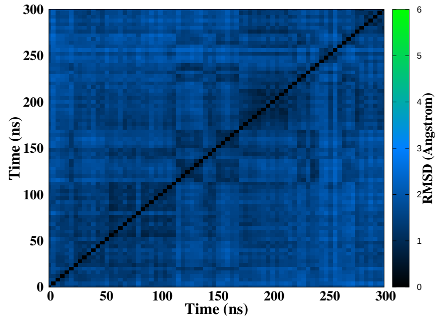


(I)

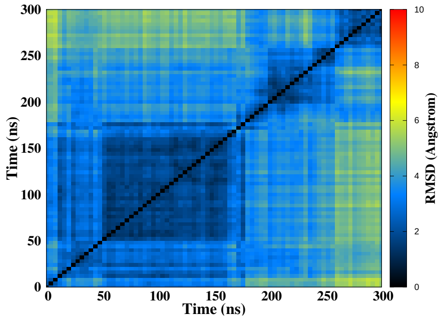


(J)

Figure S16 – Simulation of the CA damage at position 21-22, run 2. Snapshots of the initial and final MD simulation (A-B). RMSD of guanines forming the tetrads (C), distance between the tetrads (D), twist angle (E), angle between guanines for the first (F), second (G) and the third (H) tetrad, 2D-RMSD of guanines forming the tetrads (I) and the whole DNA (J).

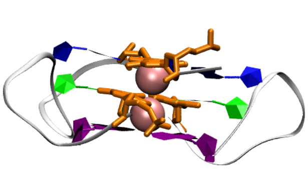

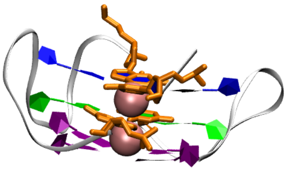

(A) 0 ns (B) 1000 ns

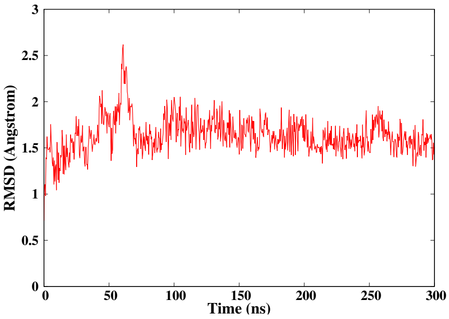


(C)

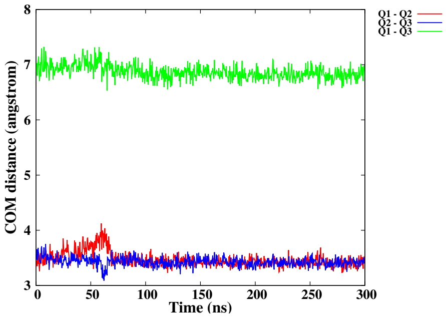


(D)

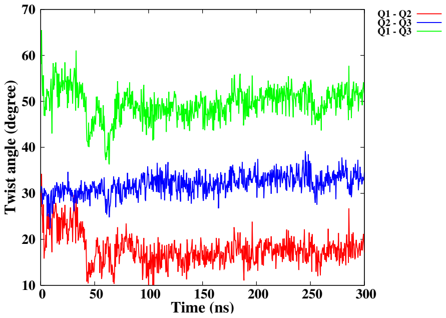


(E)

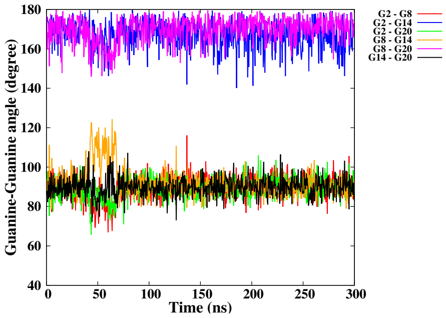


(F)

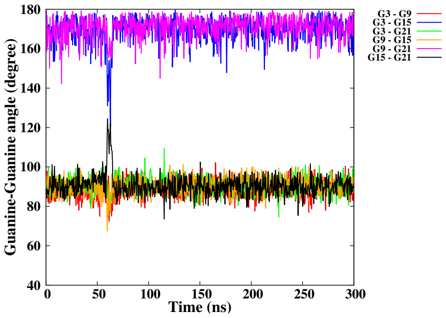


(G)

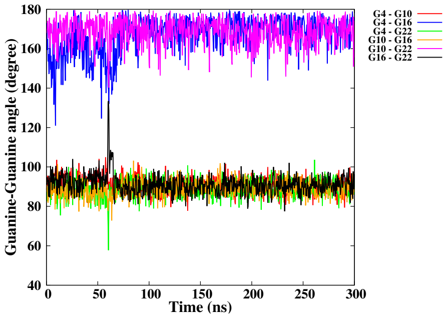


(H)

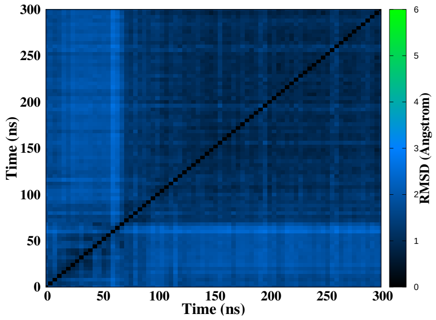


(I)

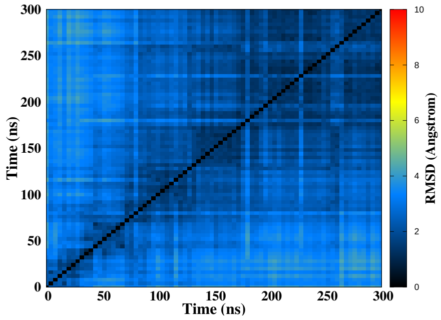


(J)

Figure S17 – Simulation of the CA damage at position 2-3/14-15, run 1. Snapshots of the initial and final MD simulation (A-B). RMSD of guanines forming the tetrads (C), distance between the tetrads (D), twist angle (E), angle between guanines for the first (F), second (G) and the third (H) tetrad, 2D-RMSD of guanines forming the tetrads (I) and the whole DNA (J).

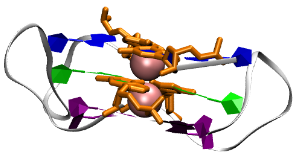

(A) 0 ns

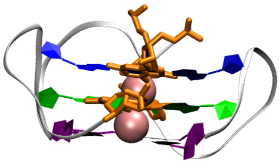

(B) 1000 ns

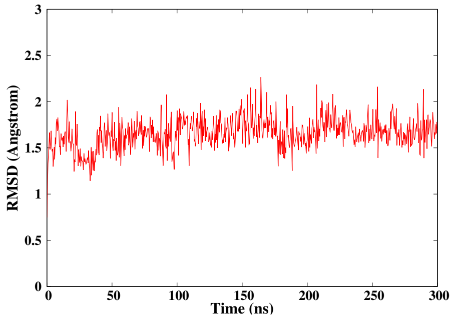


(C)

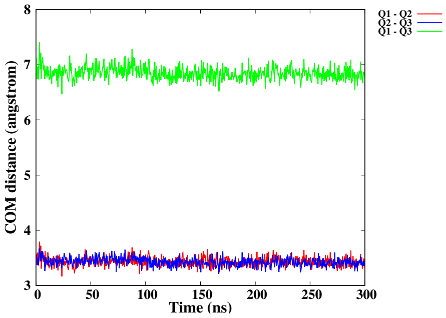


(D)

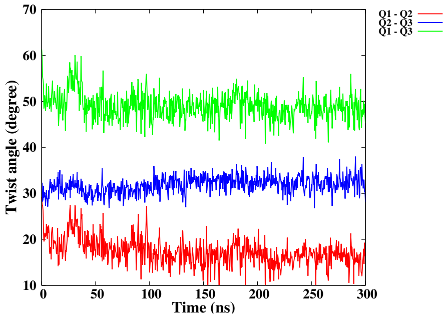


(E)

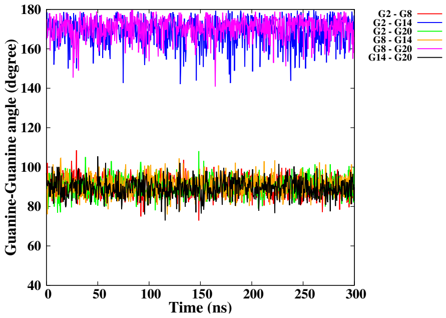


(F)

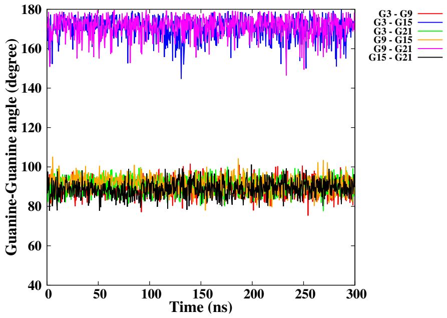


(G)

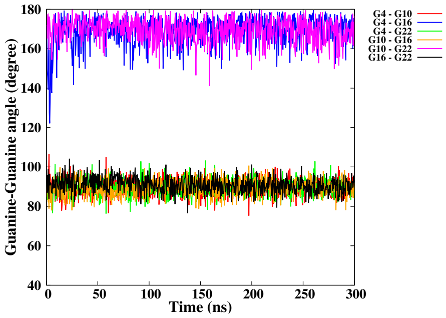


(H)

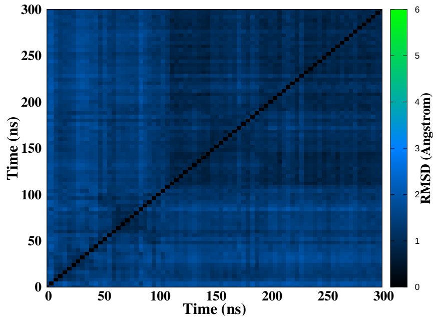


(I)

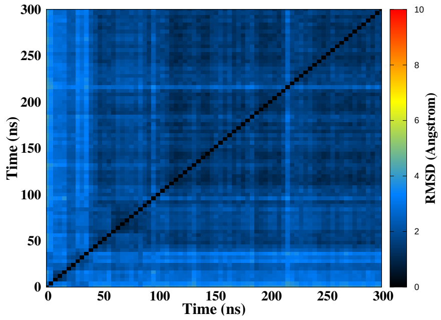


(J)

Figure S18 – Simulation of the CA damage at position 2-3/14-15, run 2. Snapshots of the initial and final MD simulation (A-B). RMSD of guanines forming the tetrads (C), distance between the tetrads (D), twist angle (E), angle between guanines for the first (F), second (G) and the third (H) tetrad, 2D-RMSD of guanines forming the tetrads (I) and the whole DNA (J).

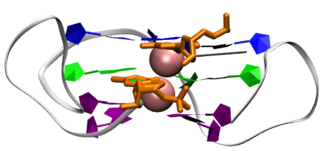

(A) 0 ns

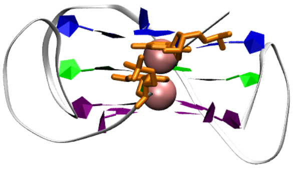

(B) 1000 ns

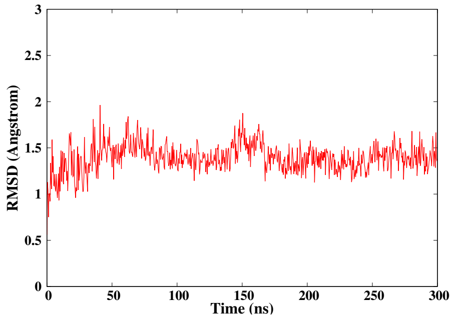


(C)

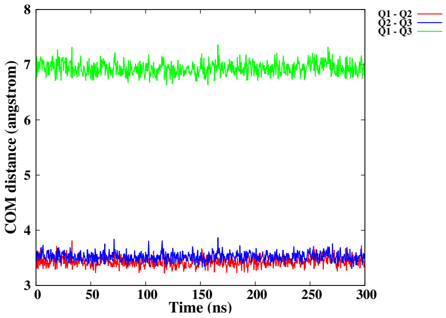


(D)

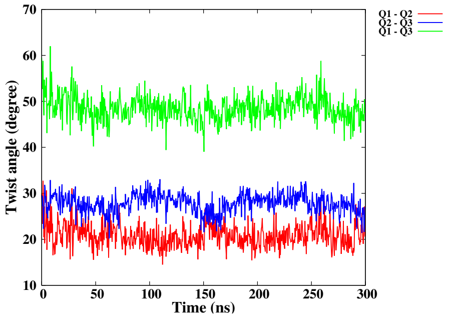


(E)

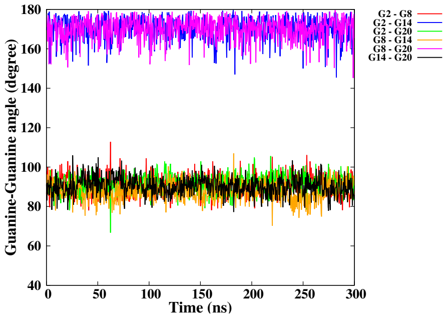


(F)

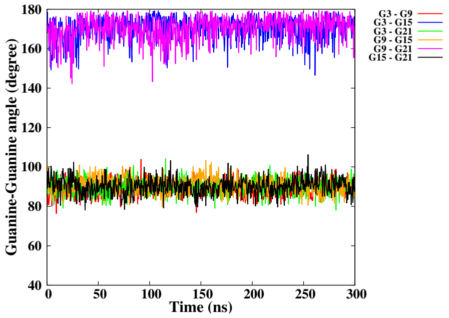


(G)

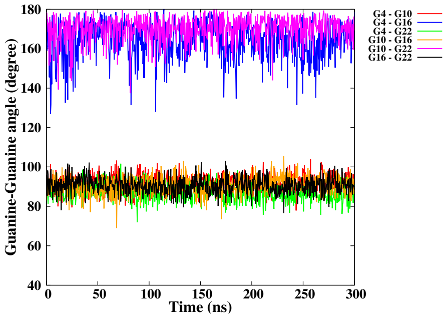


(H)

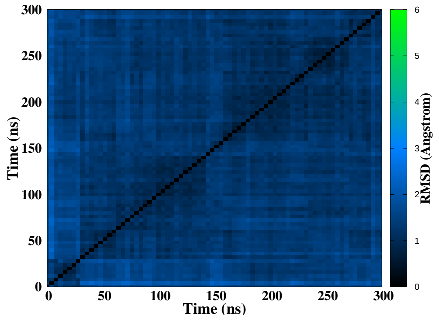


(I)

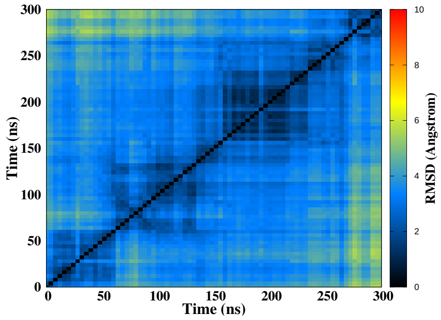


(J)

Figure S19 – Simulation of the CA damage at position 2-3, run 1. Snapshots of the initial and final MD simulation (A-B). RMSD of guanines forming the tetrads (C), distance between the tetrads (D), twist angle (E), angle between guanines for the first (F), second (G) and the third (H) tetrad, 2D-RMSD of guanines forming the tetrads (I) and the whole DNA (J).

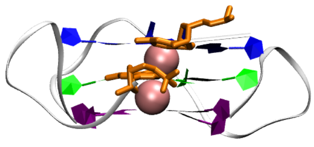

(A) 0 ns

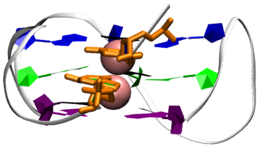

(B) 1000 ns

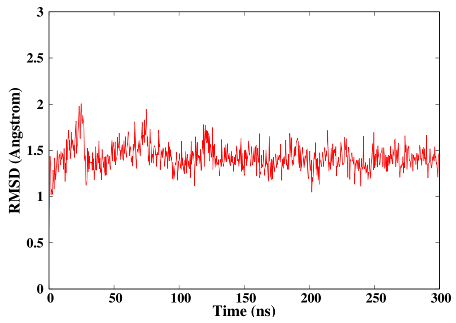


(C)

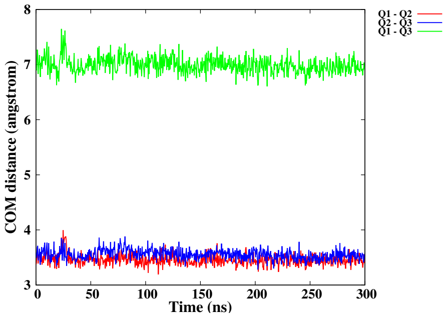


(D)

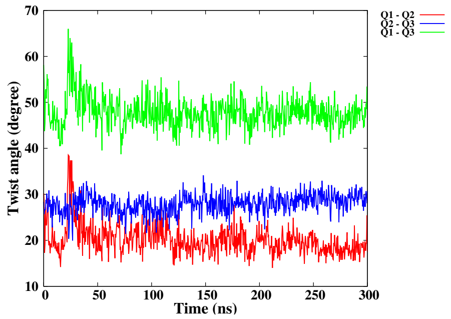


(E)

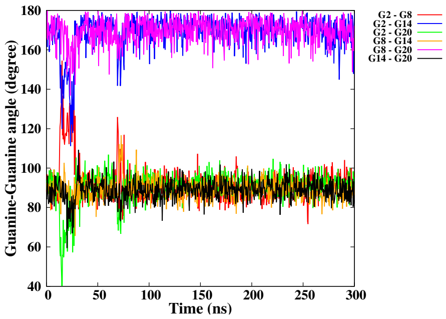


(F)

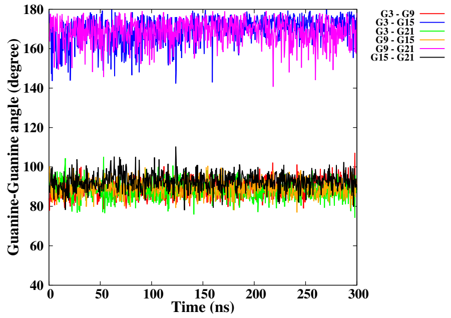


(G)

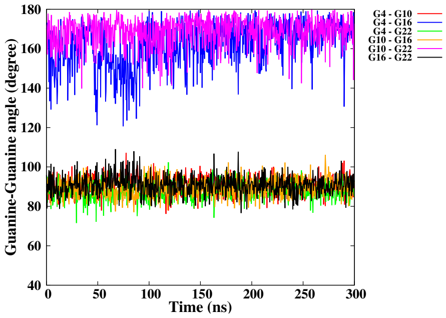


(H)

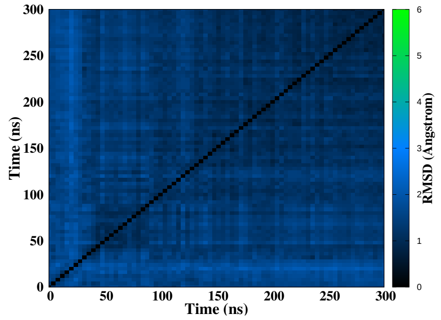


(I)

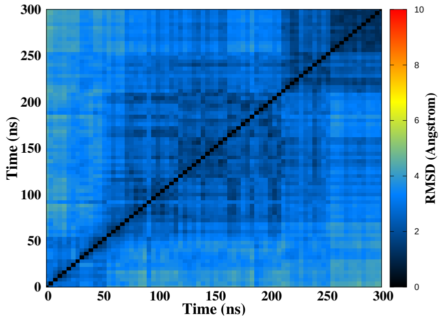


(J)

Figure S20 – Simulation of the CA damage at position 2-3, run 2. Snapshots of the initial and final MD simulation (A-B). RMSD of guanines forming the tetrads (C), distance between the tetrads (D), twist angle (E), angle between guanines for the first (F), second (G) and the third (H) tetrad, 2D-RMSD of guanines forming the tetrads (I) and the whole DNA (J).

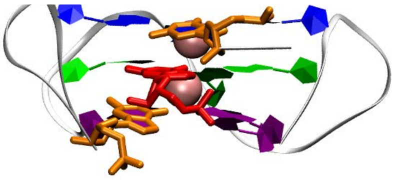

(A) 0 ns

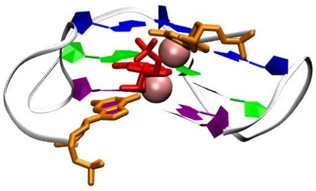

(B) 1000 ns

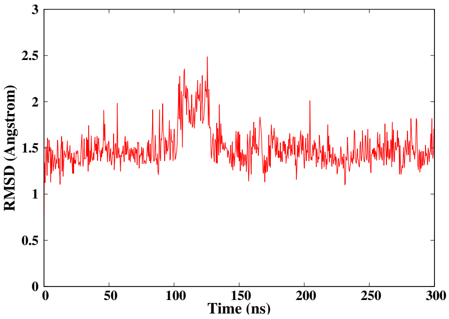


(C)

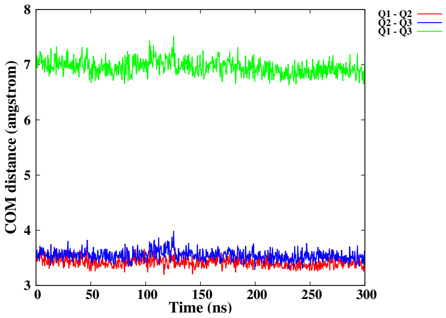


(D)

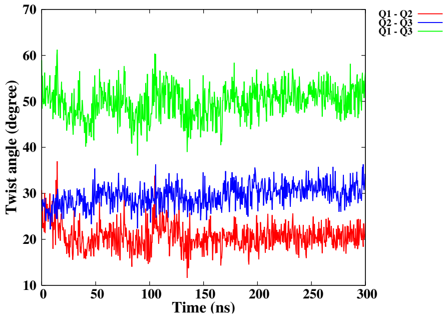


(E)

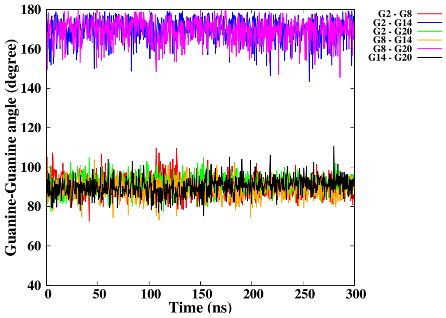


(F)

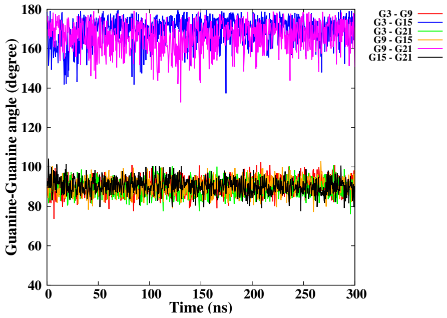


(G)

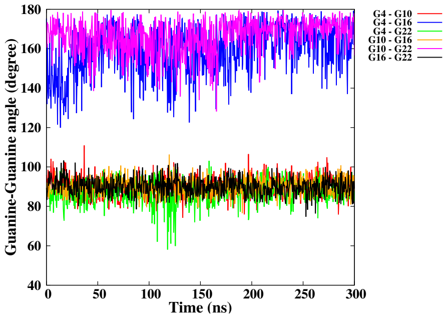


(H)

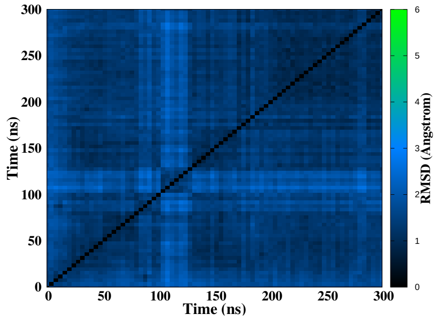


(I)

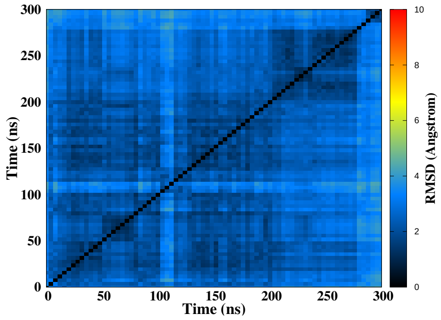


(J)

Figure S21 – Simulation of the CA damage at position 2-3-4, run 1. Snapshots of the initial and final MD simulation (A-B). RMSD of guanines forming the tetrads (C), distance between the tetrads (D), twist angle (E), angle between guanines for the first (F), second (G) and the third (H) tetrad, 2D-RMSD of guanines forming the tetrads (I) and the whole DNA (J).

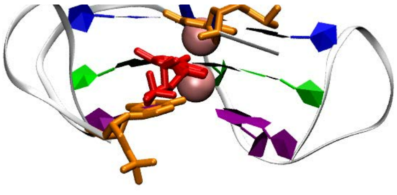

(A) 0 ns

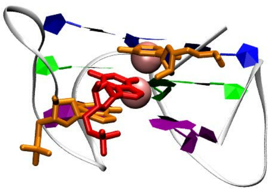

(B) 1000 ns

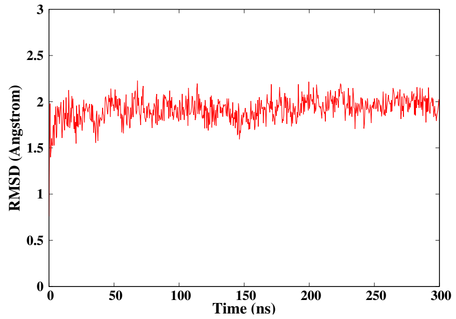


(C)

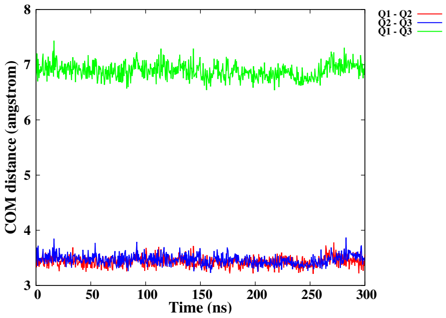


(D)

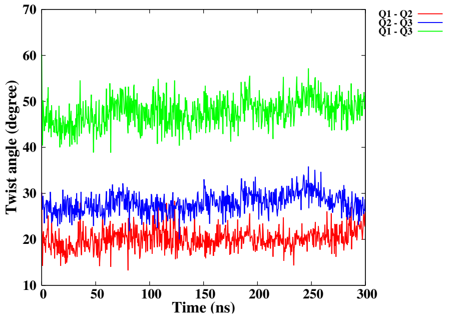


(E)

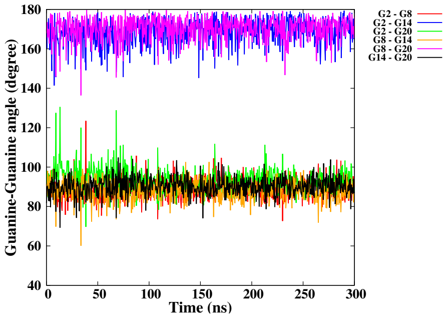


(F)

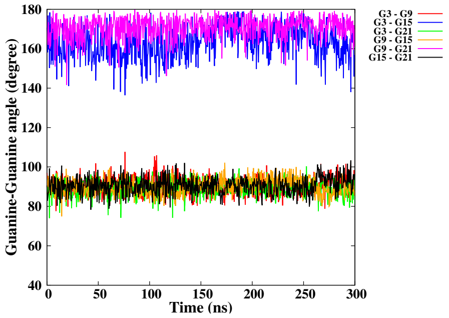


(G)

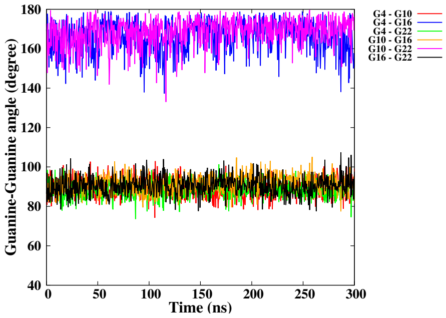


(H)

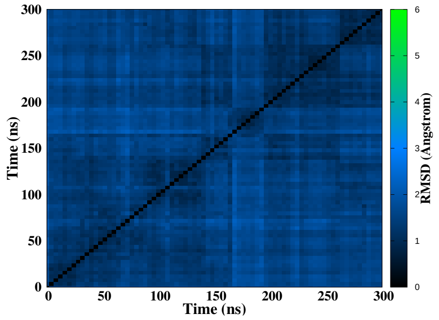


(I)

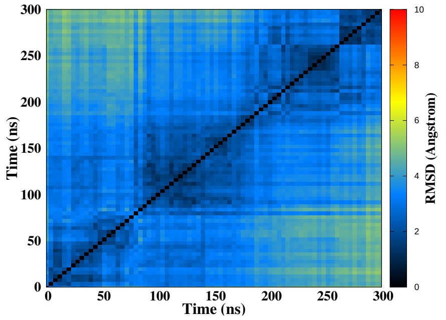


(J)

Figure S22 – Simulation of the CA damage at position 2-3-4, run 2. Snapshots of the initial and final MD simulation (A-B). RMSD of guanines forming the tetrads (C), distance between the tetrads (D), twist angle (E), angle between guanines for the first (F), second (G) and the third (H) tetrad, 2D-RMSD of guanines forming the tetrads (I) and the whole DNA (J).

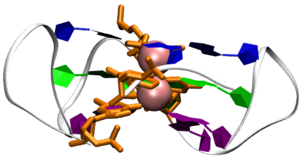

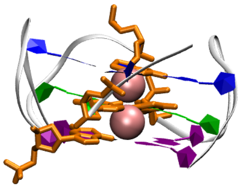

(A) 0 ns (B) 1000 ns

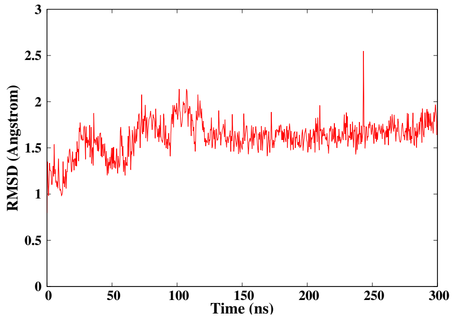


(C)

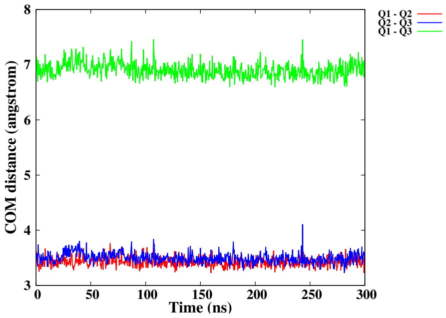


(D)

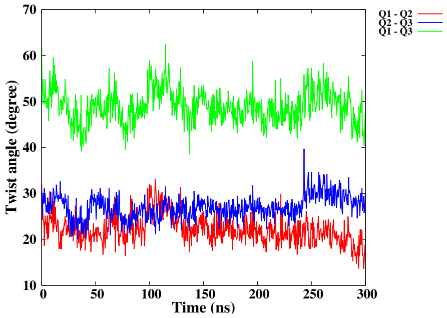


(E)

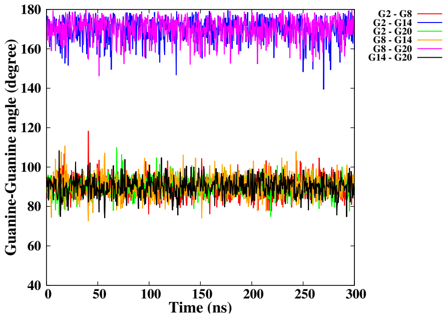


(F)

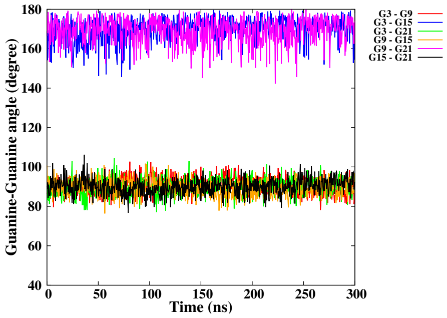


(G)

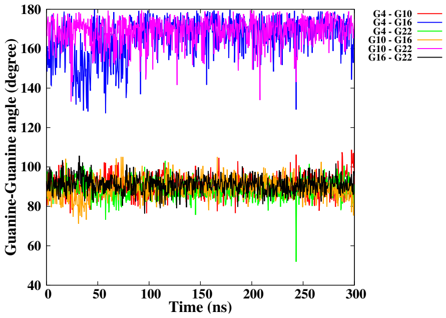


(H)

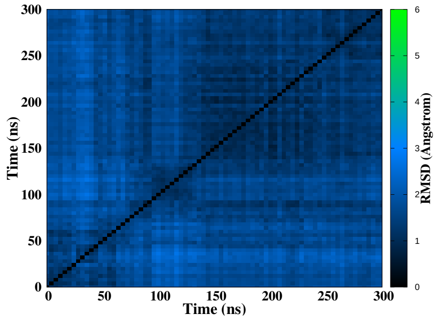


(I)

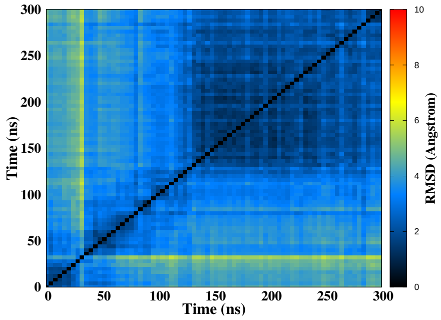


(J)

Figure S23 – Simulation of the CA damage at position 3-4/14-15, run 1. Snapshots of the initial and final MD simulation (A-B). RMSD of guanines forming the tetrads (C), distance between the tetrads (D), twist angle (E), angle between guanines for the first (F), second (G) and the third (H) tetrad, 2D-RMSD of guanines forming the tetrads (I) and the whole DNA (J).

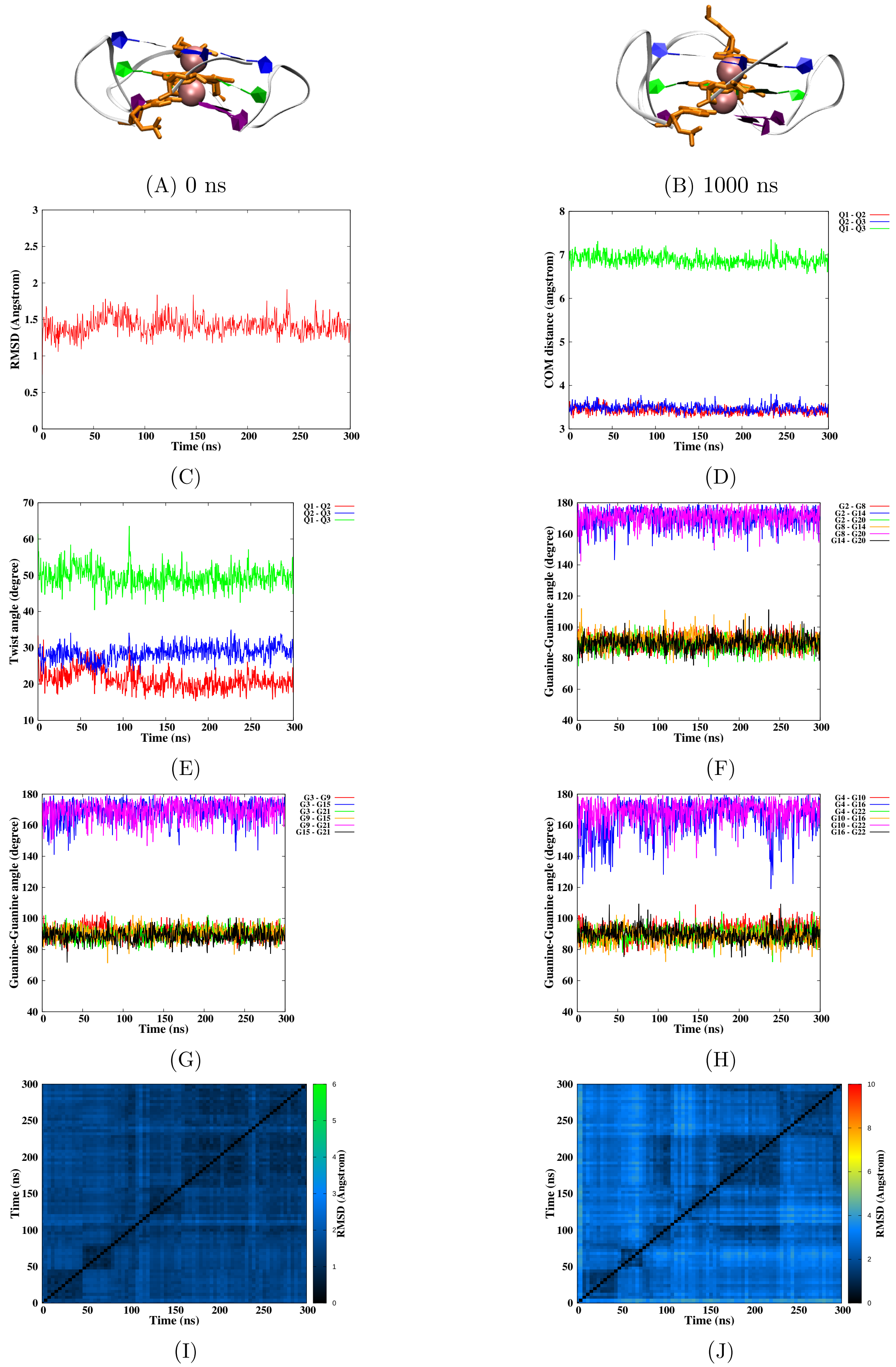


Figure S24 – Simulation of the CA damage at position 3-4/14-15, run 2. Snapshots of the initial and final MD simulation (A-B). RMSD of guanines forming the tetrads (C), distance between the tetrads (D), twist angle (E), angle between guanines for the first (F), second (G) and the third (H) tetrad, 2D-RMSD of guanines forming the tetrads (I) and the whole DNA (J).

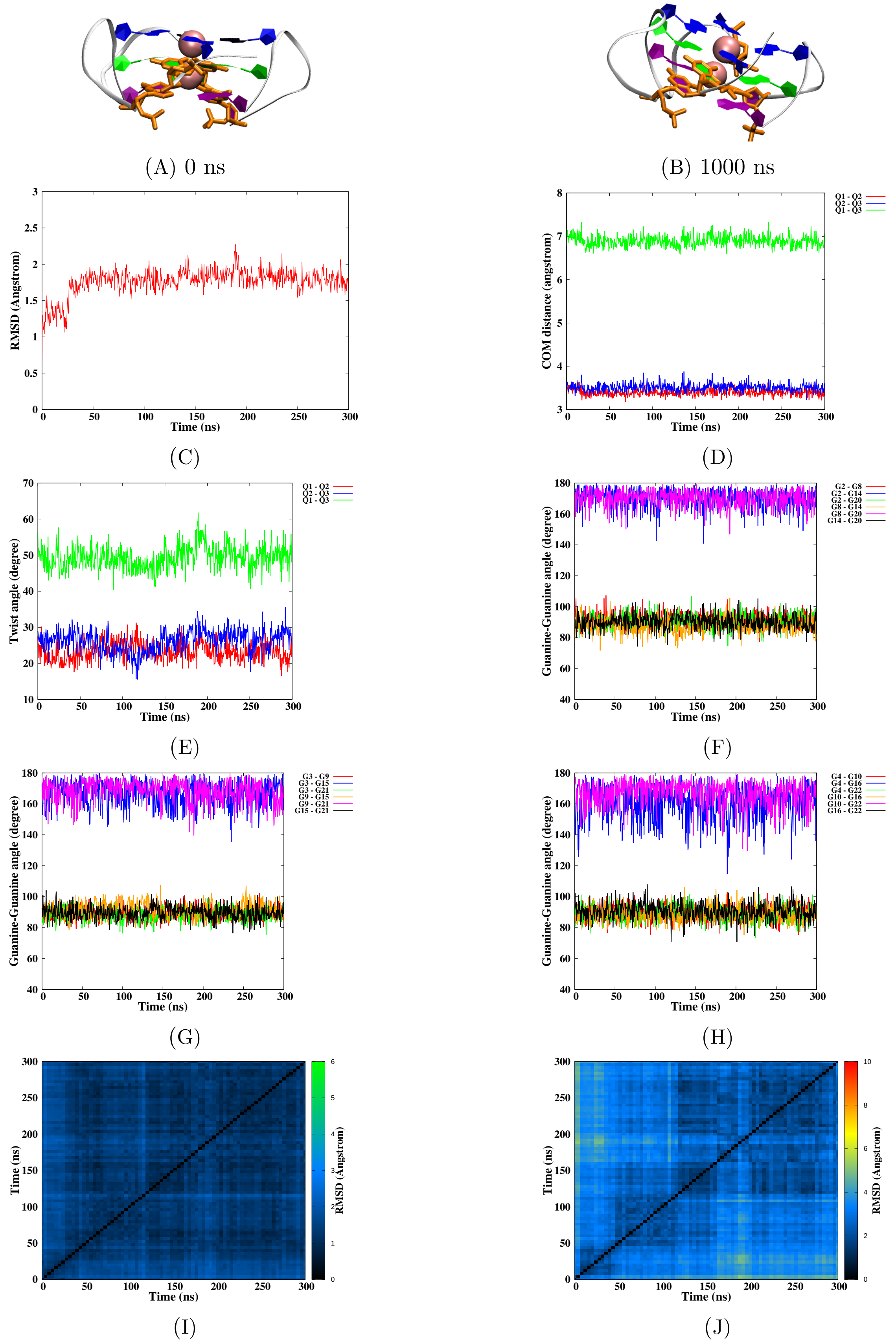


Figure S25 – Simulation of the CA damage at position 3-4/15-16, run 1. Snapshots of the initial and final MD simulation (A-B). RMSD of guanines forming the tetrads (C), distance between the tetrads (D), twist angle (E), angle between guanines for the first (F), second (G) and the third (H) tetrad, 2D-RMSD of guanines forming the tetrads (I) and the whole DNA (J).

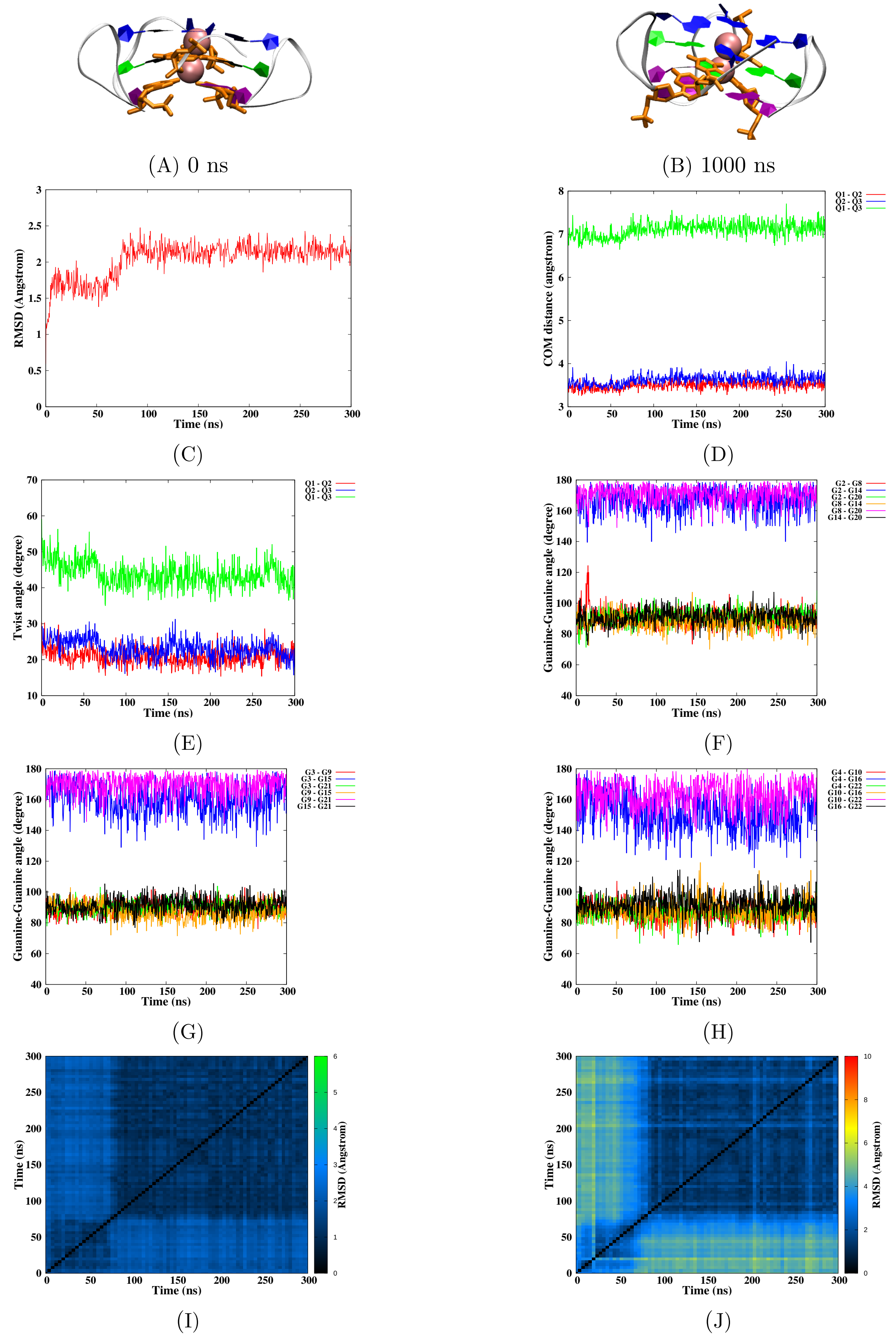


Figure S26 – Simulation of the CA damage at position 3-4/15-16, run 2. Snapshots of the initial and final MD simulation (A-B). RMSD of guanines forming the tetrads (C), distance between the tetrads (D), twist angle (E), angle between guanines for the first (F), second (G) and the third (H) tetrad, 2D-RMSD of guanines forming the tetrads (I) and the whole DNA (J).

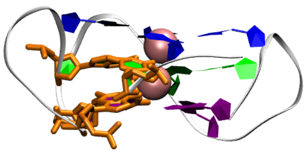

(A) 0 ns

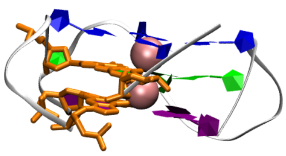

(B) 1000 ns

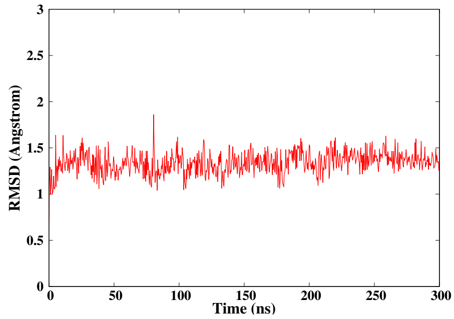


(C)

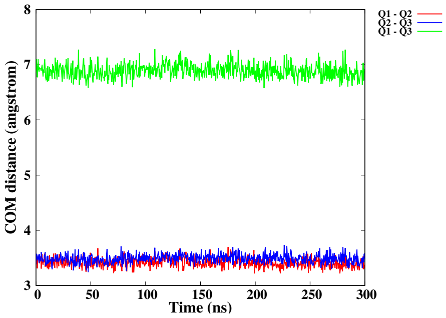


(D)

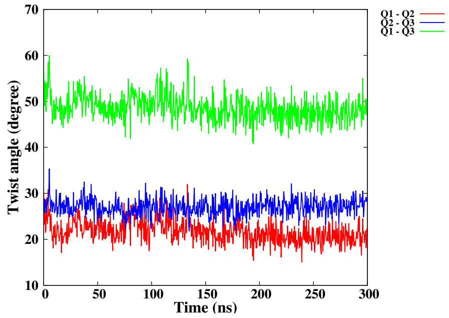


(E)

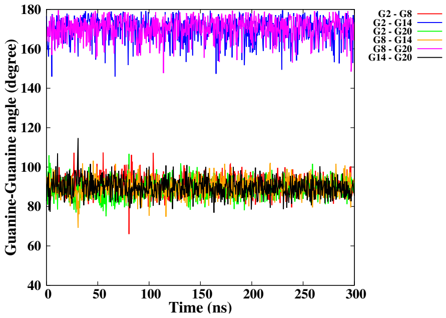


(F)

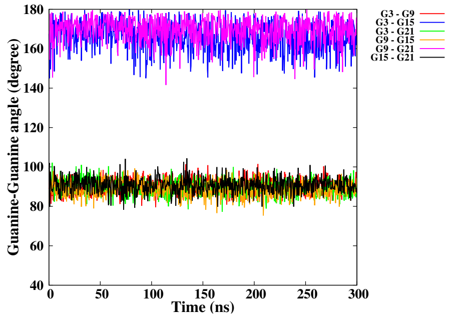


(G)

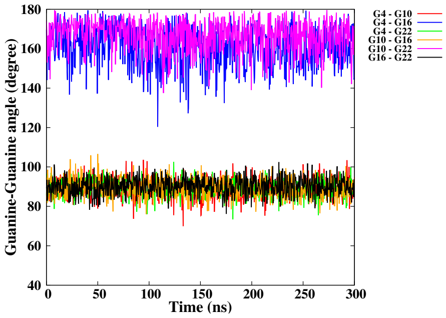


(H)

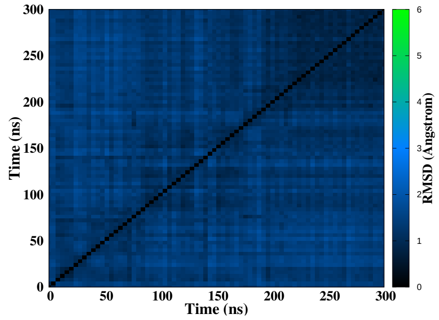


(I)

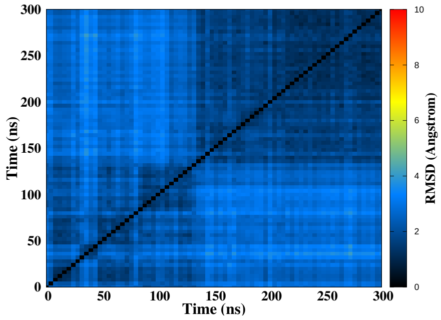


(J)

Figure S27 – Simulation of the CA damage at position 3-4/9-10, run 1. Snapshots of the initial and final MD simulation (A-B). RMSD of guanines forming the tetrads (C), distance between the tetrads (D), twist angle (E), angle between guanines for the first (F), second (G) and the third (H) tetrad, 2D-RMSD of guanines forming the tetrads (I) and the whole DNA (J).

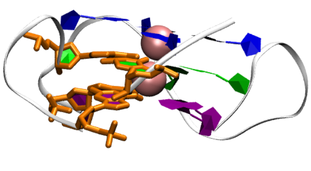

(A) 0 ns

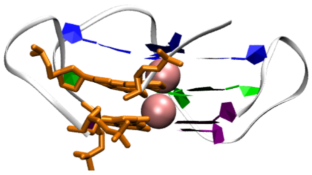

(B) 1000 ns

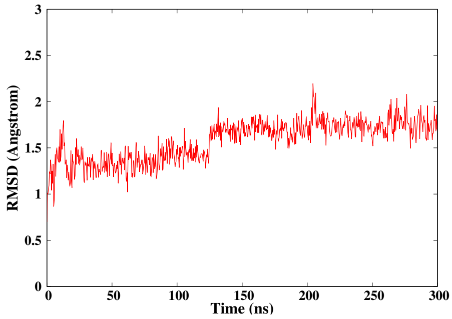


(C)

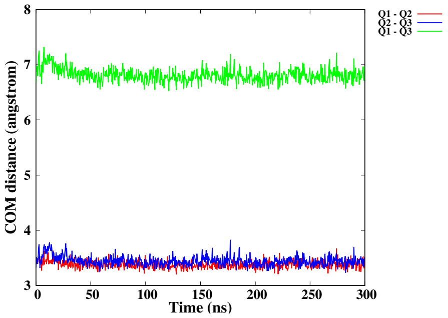


(D)

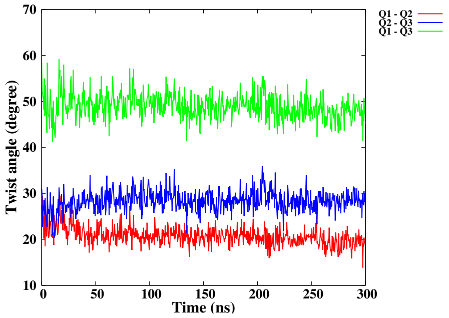


(E)

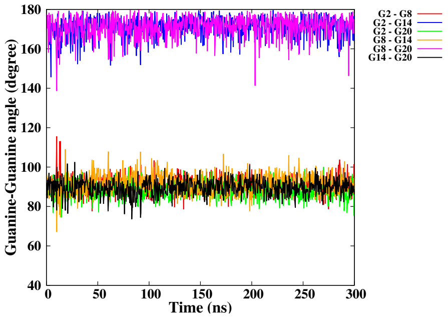


(F)

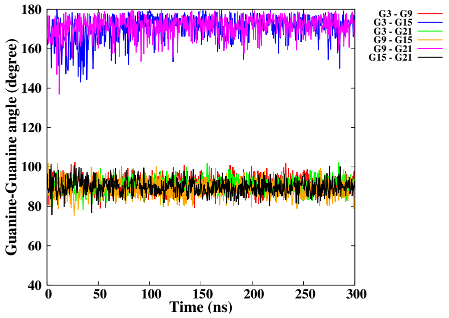


(G)

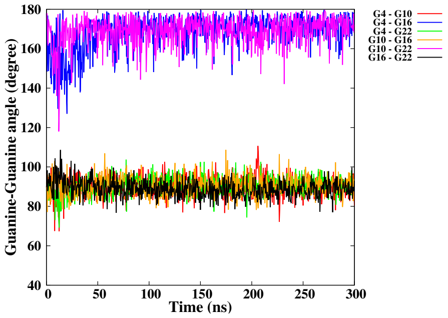


(H)

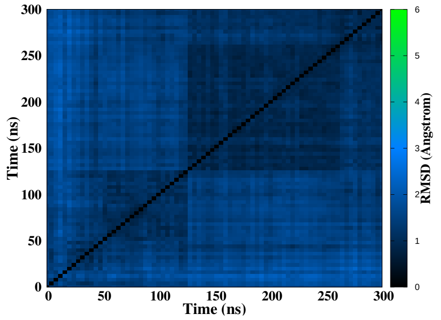


(I)

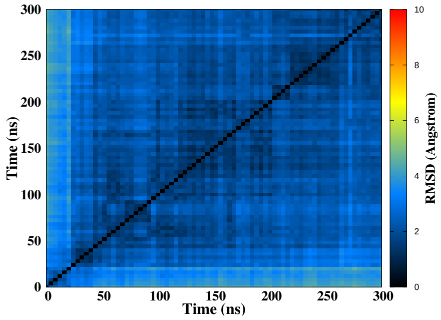


(J)

Figure S28 – Simulation of the CA damage at position 3-4/9-10, run 2. Snapshots of the initial and final MD simulation (A-B). RMSD of guanines forming the tetrads (C), distance between the tetrads (D), twist angle (E), angle between guanines for the first (F), second (G) and the third (H) tetrad, 2D-RMSD of guanines forming the tetrads (I) and the whole DNA (J).

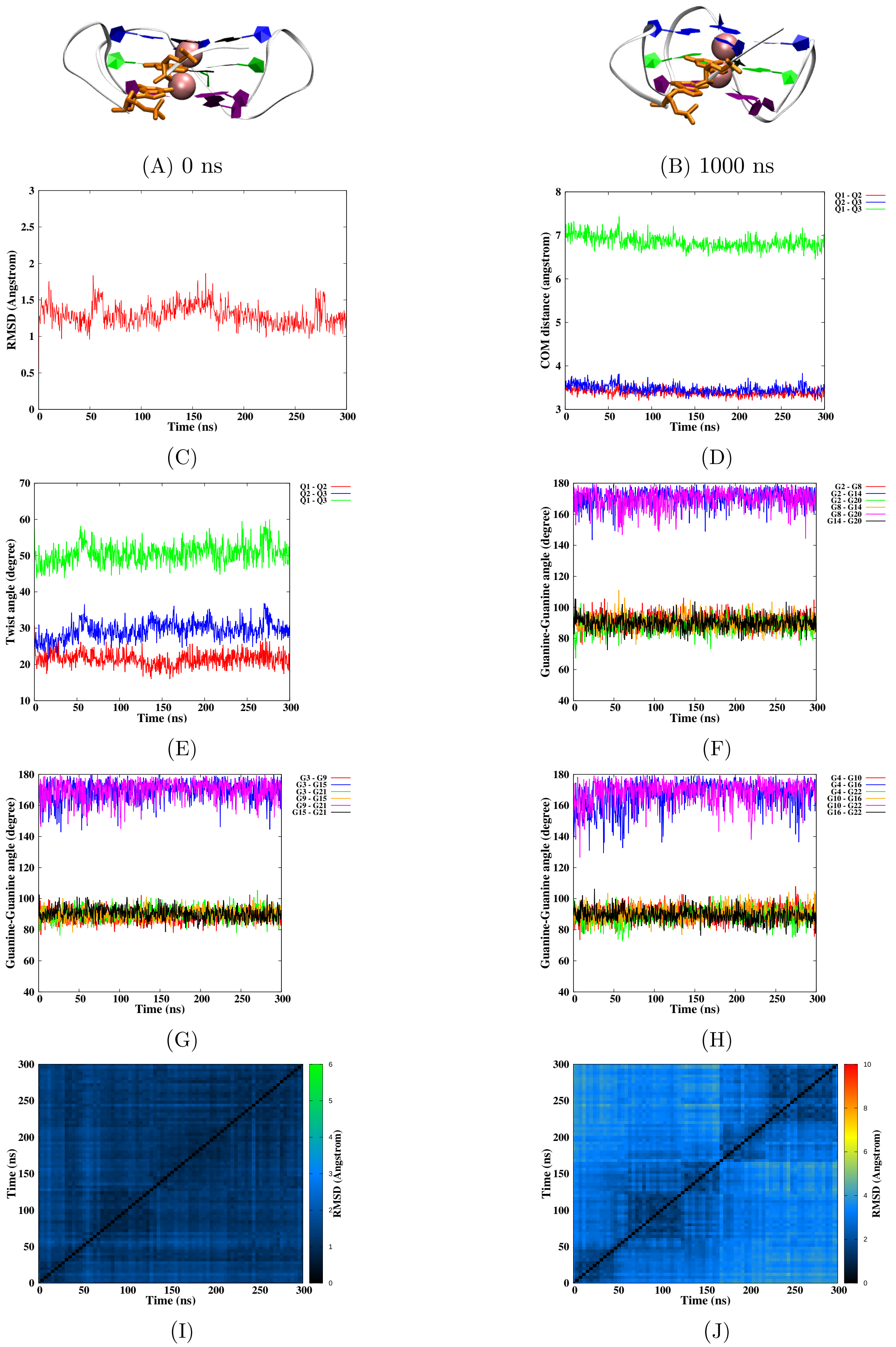


Figure S29 – Simulation of the CA damage at position 3-4, run 1. Snapshots of the initial and final MD simulation (A-B). RMSD of guanines forming the tetrads (C), distance between the tetrads (D), twist angle (E), angle between guanines for the first (F), second (G) and the third (H) tetrad, 2D-RMSD of guanines forming the tetrads (I) and the whole DNA (J).

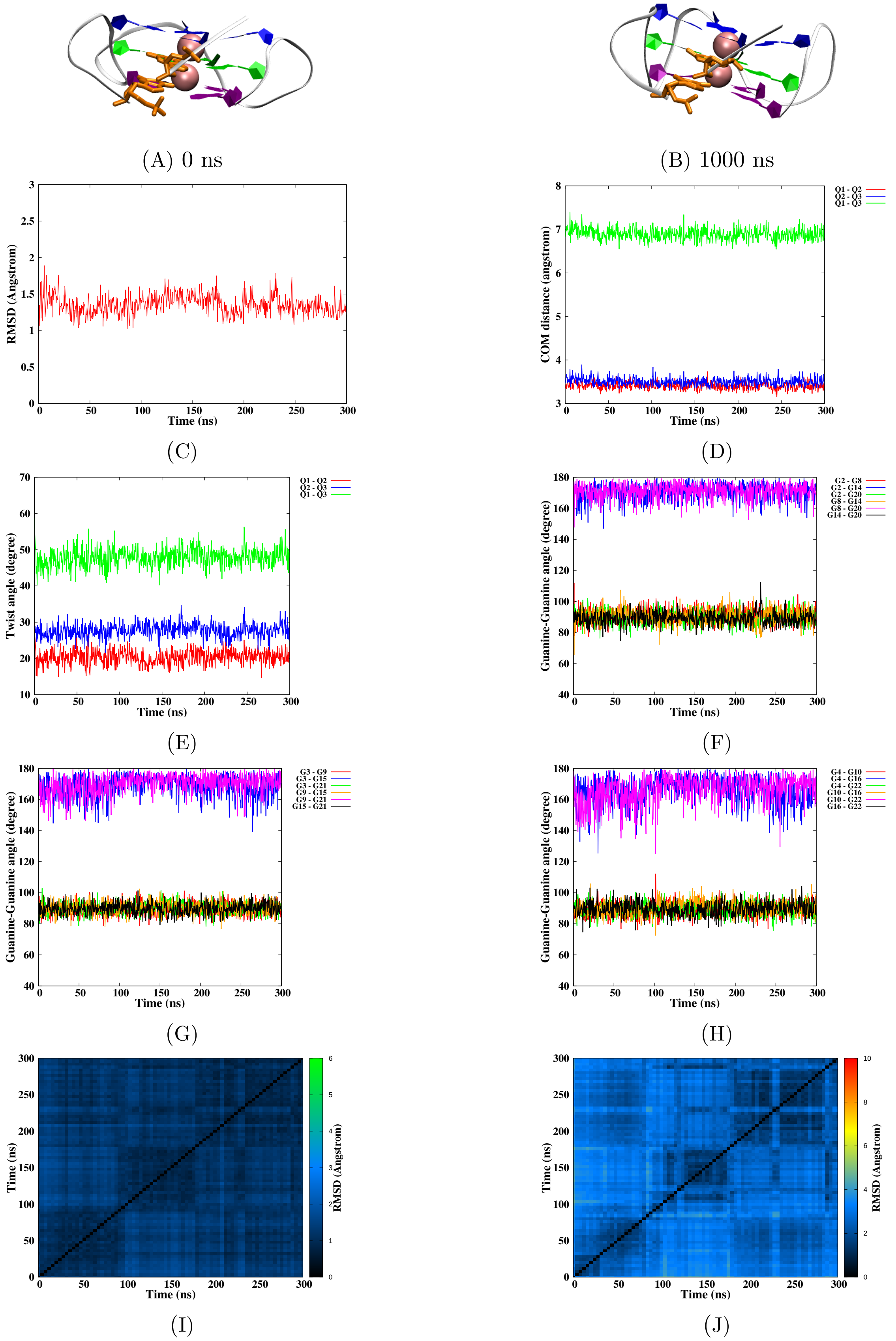


Figure S30 – Simulation of the CA damage at position 3-4, run 2. Snapshots of the initial and final MD simulation (A-B). RMSD of guanines forming the tetrads (C), distance between the tetrads (D), twist angle (E), angle between guanines for the first (F), second (G) and the third (H) tetrad, 2D-RMSD of guanines forming the tetrads (I) and the whole DNA (J).

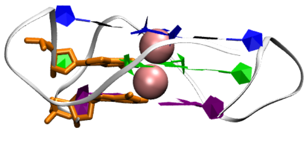

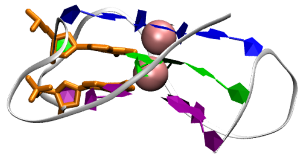

(A) 0 ns

(B) 1000 ns

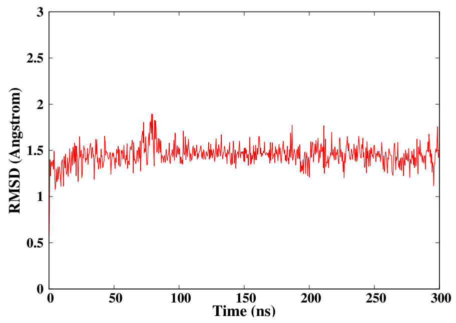


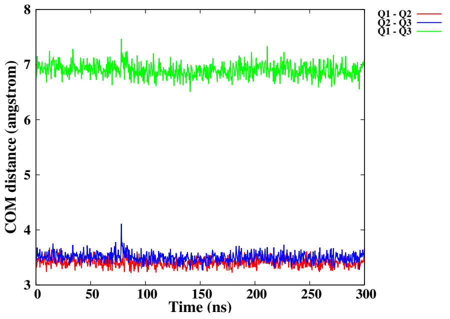


(C)

(D)

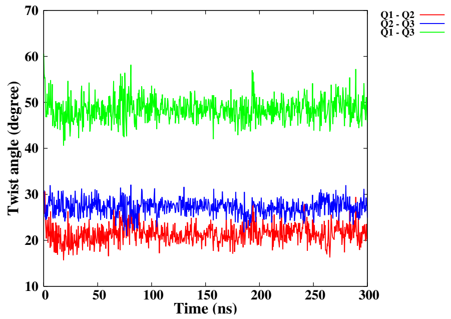


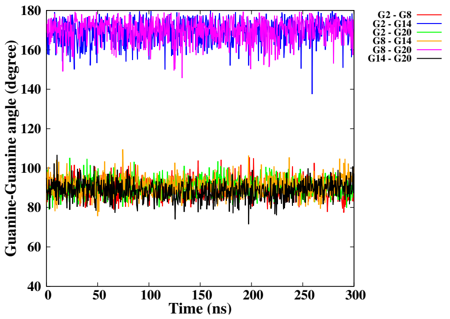


(E)

(F)

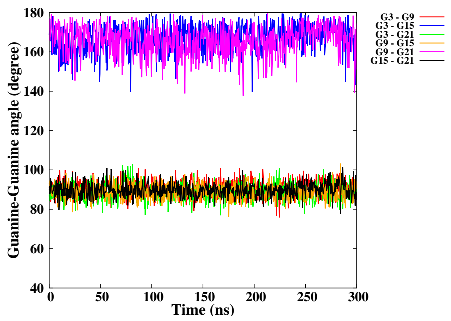


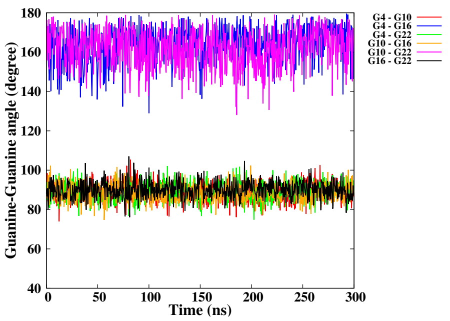


(G)

(H)

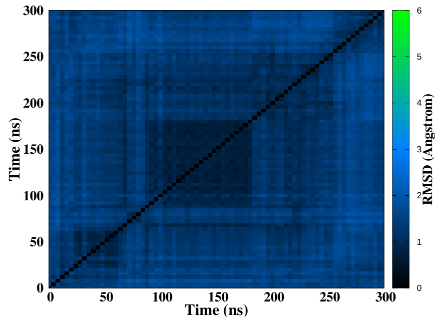


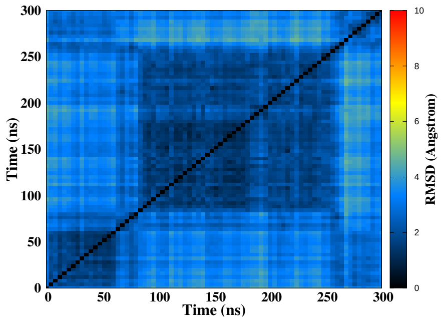


(I)

(J)

Figure S31 – Simulation of the CA damage at position 9-10, run 1. Snapshots of the initial and final MD simulation (A-B). RMSD of guanines forming the tetrads (C), distance between the tetrads (D), twist angle (E), angle between guanines for the first (F), second (G) and the third (H) tetrad, 2D-RMSD of guanines forming the tetrads (I) and the whole DNA (J).

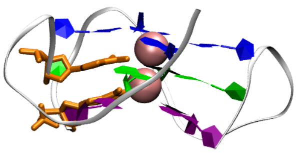

(A) 0 ns

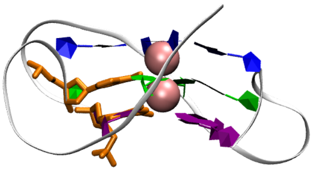

(B) 1000 ns

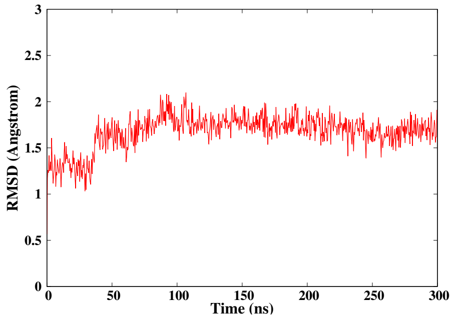


(C)

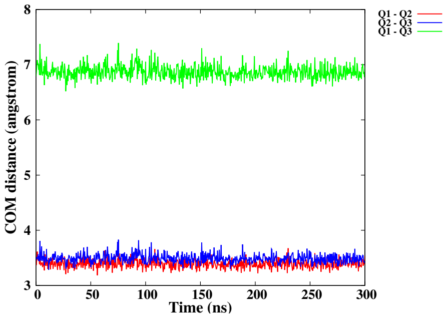


(D)

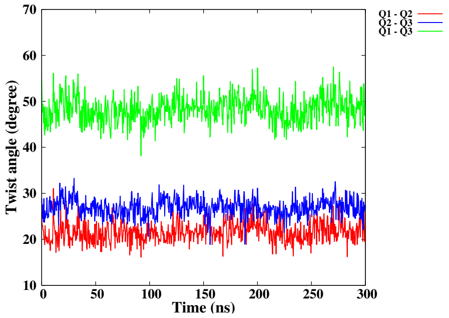


(E)

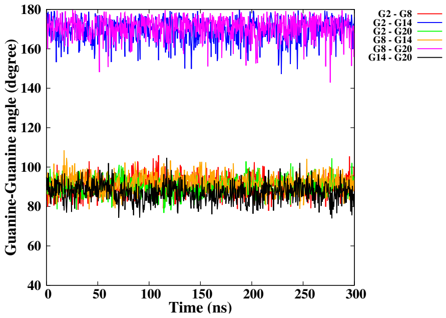


(F)

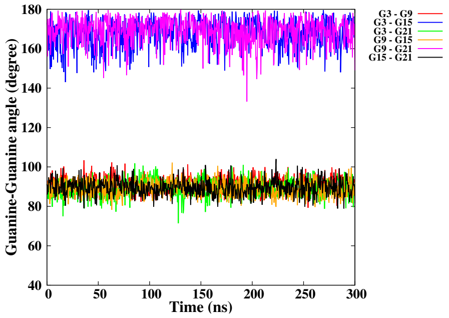


(G)

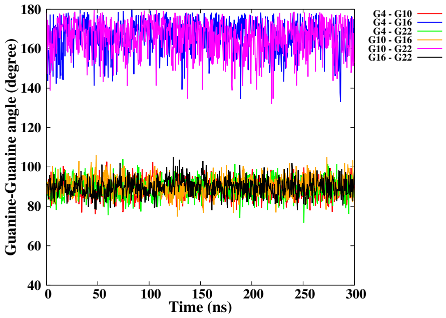


(H)

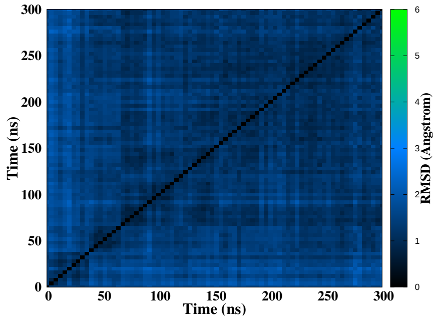


(I)

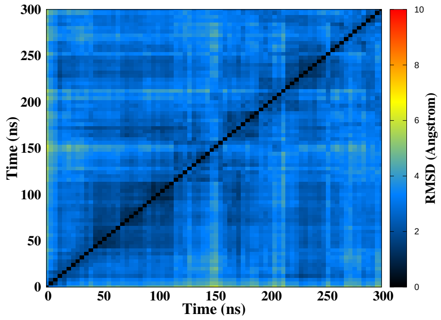


(J)

Figure S32 – Simulation of the CA damage at position 9-10, run 2. Snapshots of the initial and final MD simulation (A-B). RMSD of guanines forming the tetrads (C), distance between the tetrads (D), twist angle (E), angle between guanines for the first (F), second (G) and the third (H) tetrad, 2D-RMSD of guanines forming the tetrads (I) and the whole DNA (J).

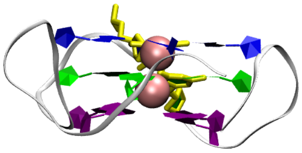

(A) 0 ns

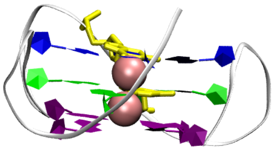

(B) 1000 ns

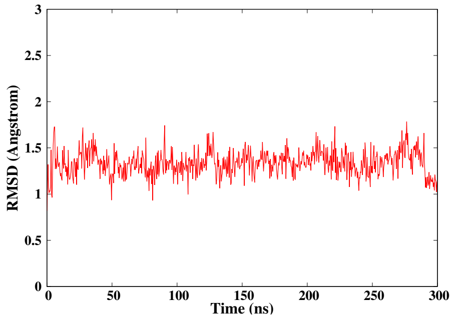


(C)

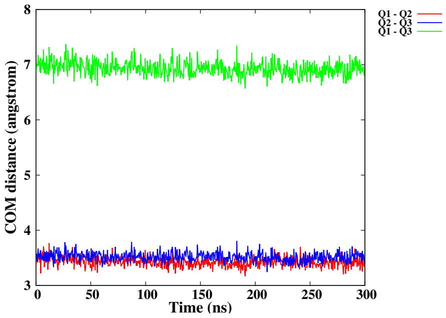


(D)

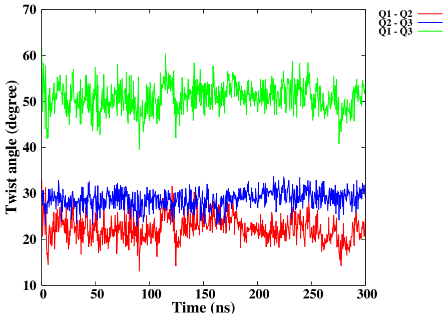


(E)

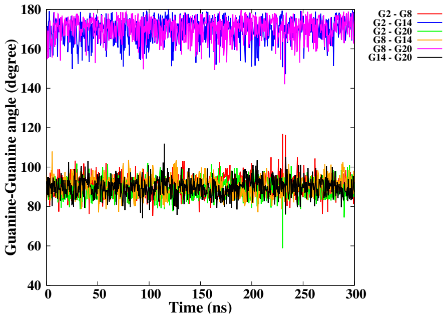


(F)

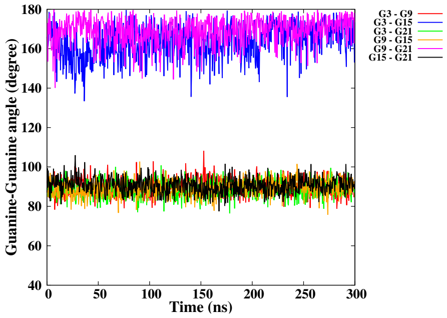


(G)

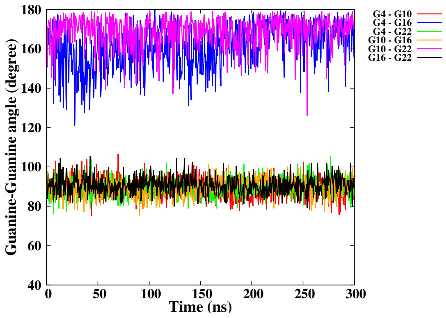


(H)

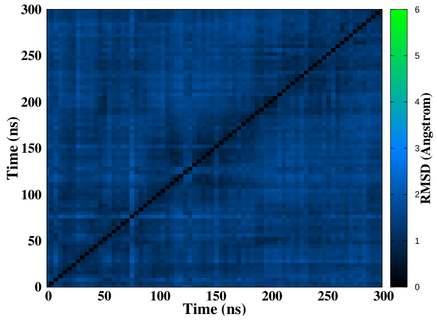


(I)

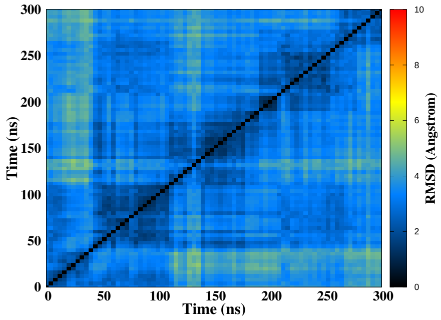


(J)

Figure S33 – Simulation of the NC damage at position 14-15, run 1. Snapshots of the initial and final MD simulation (A-B). RMSD of guanines forming the tetrads (C), distance between the tetrads (D), twist angle (E), angle between guanines for the first (F), second (G) and the third (H) tetrad, 2D-RMSD of guanines forming the tetrads (I) and the whole DNA (J).

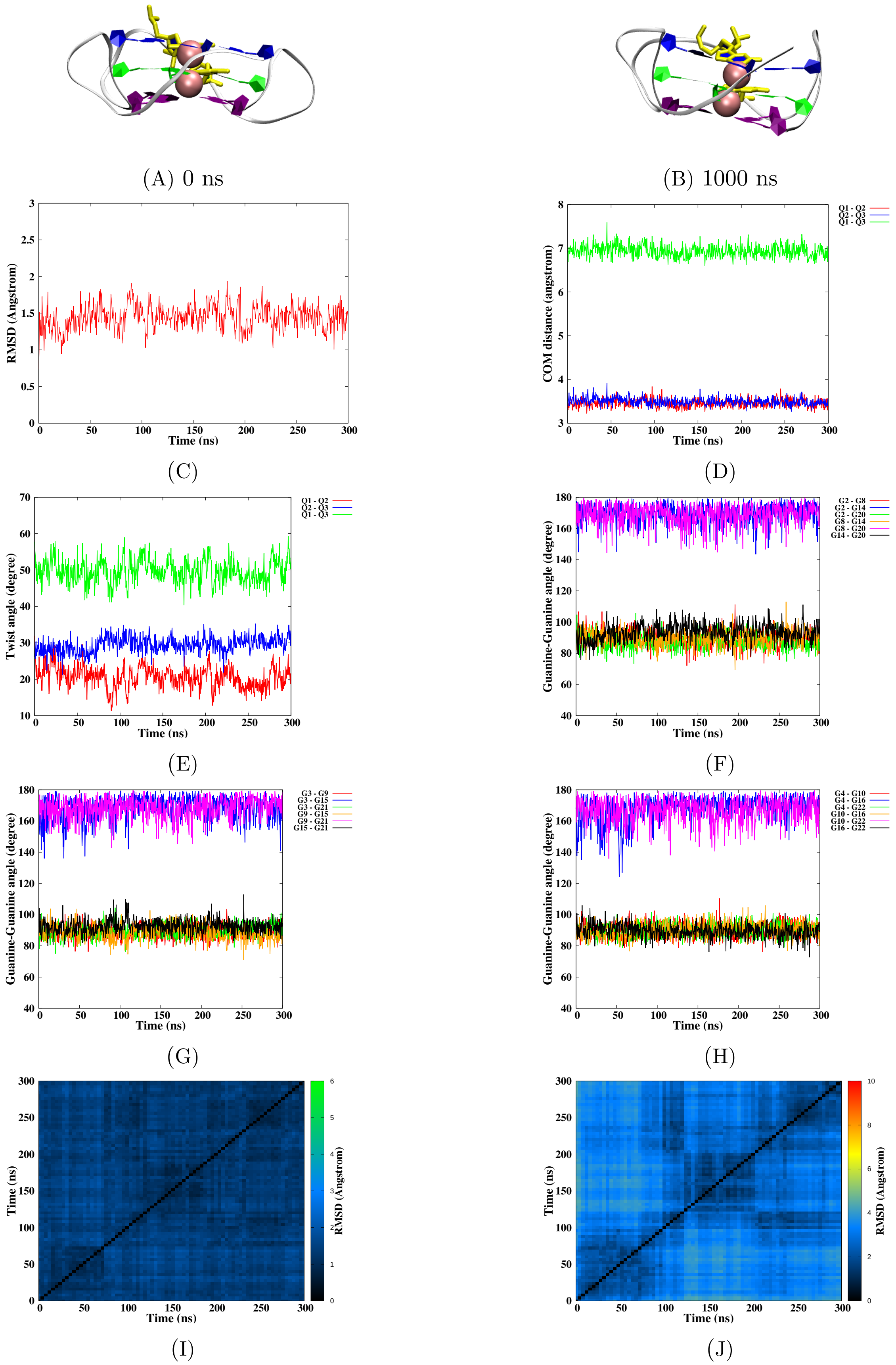


Figure S34 – Simulation of the NC damage at position 14-15, run 2. Snapshots of the initial and final MD simulation (A-B). RMSD of guanines forming the tetrads (C), distance between the tetrads (D), twist angle (E), angle between guanines for the first (F), second (G) and the third (H) tetrad, 2D-RMSD of guanines forming the tetrads (I) and the whole DNA (J).

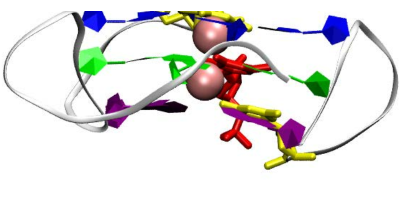

(A) 0 ns

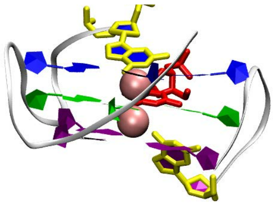

(B) 1000 ns

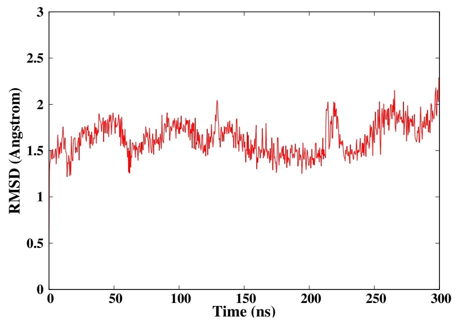


(C)

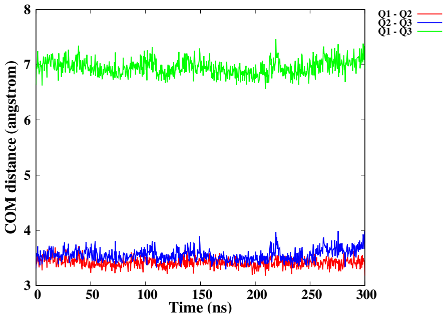


(D)

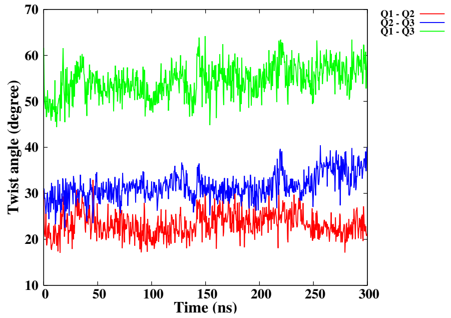


(E)

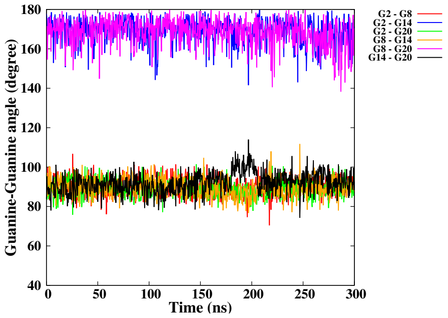


(F)

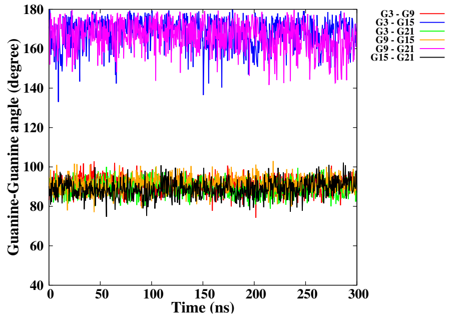


(G)

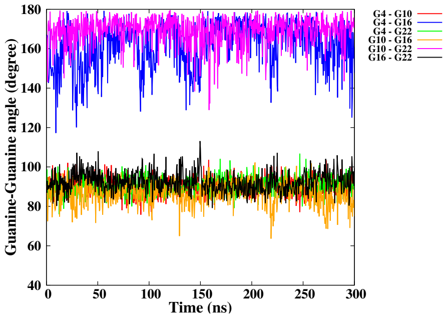


(H)

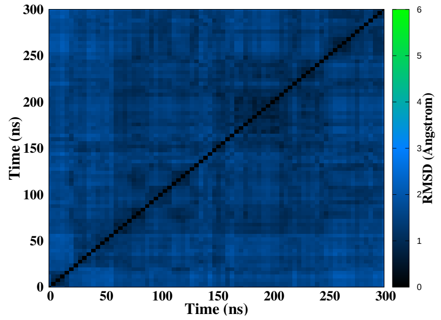


(I)

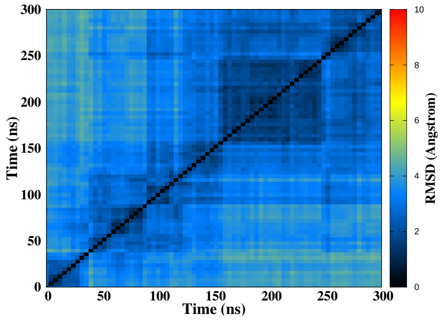


(J)

Figure S35 – Simulation of the NC damage at position 14-15-16, run 1. Snapshots of the initial and final MD simulation (A-B). RMSD of guanines forming the tetrads (C), distance between the tetrads (D), twist angle (E), angle between guanines for the first (F), second (G) and the third (H) tetrad, 2D-RMSD of guanines forming the tetrads (I) and the whole DNA (J).

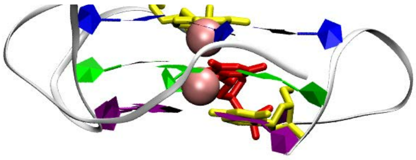

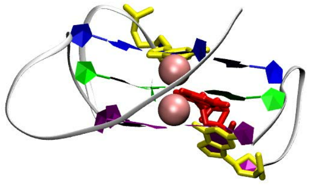

(A) 0 ns

(B) 1000 ns

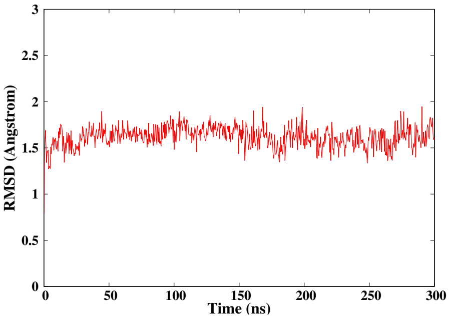


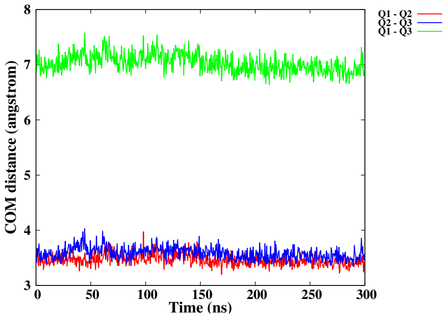


(C)

(D)

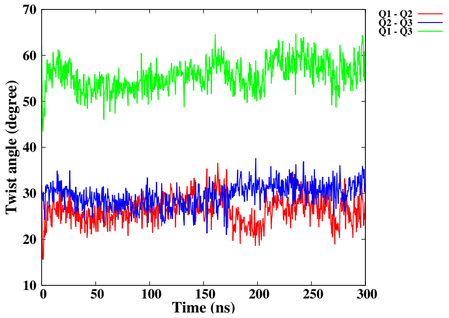


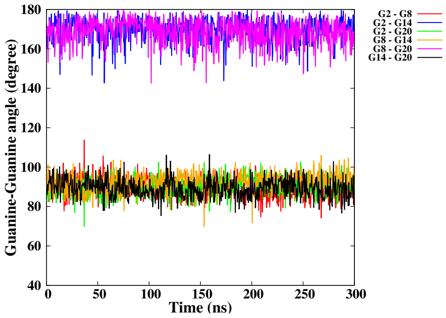


(E)

(F)

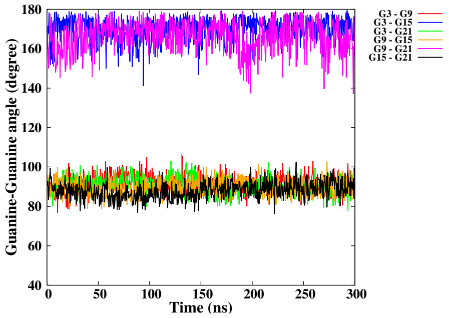


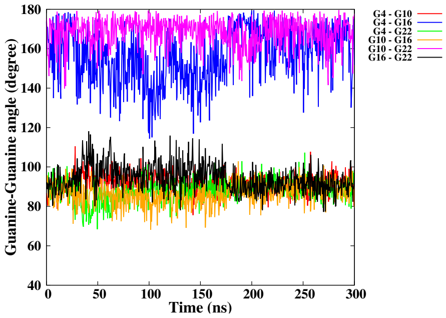


(G)

(H)

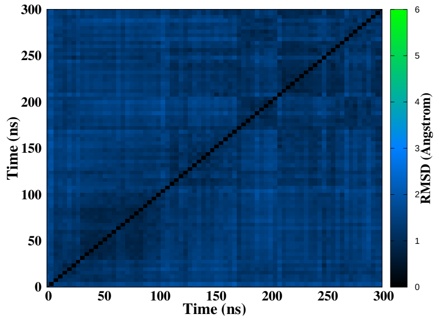


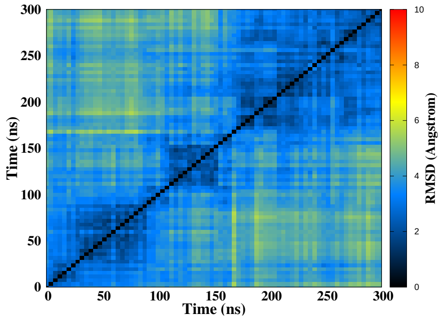


(I)

(J)

Figure S36 – Simulation of the NC damage at position 14-15-16, run 2. Snapshots of the initial and final MD simulation (A-B). RMSD of guanines forming the tetrads (C), distance between the tetrads (D), twist angle (E), angle between guanines for the first (F), second (G) and the third (H) tetrad, 2D-RMSD of guanines forming the tetrads (I) and the whole DNA (J).

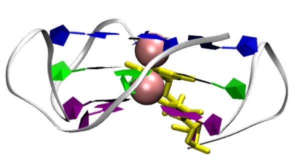

(A) 0 ns

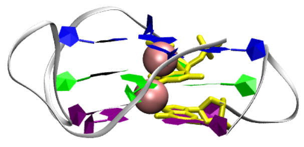

(B) 1000 ns

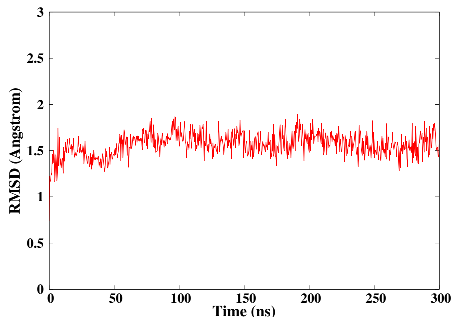


(C)

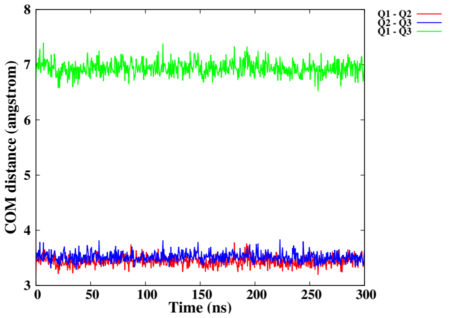


(D)

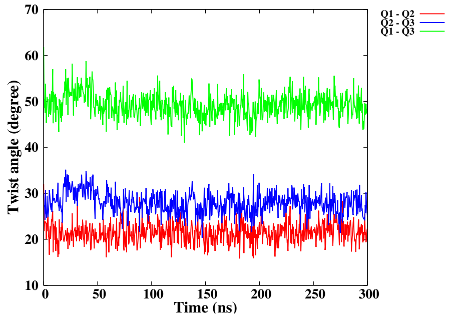


(E)

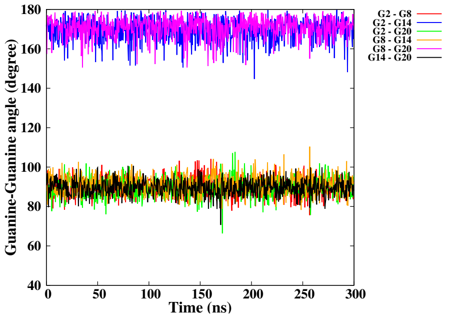


(F)

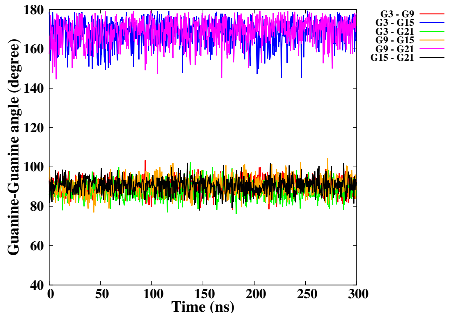


(G)

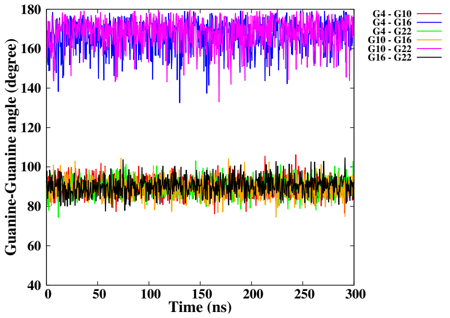


(H)

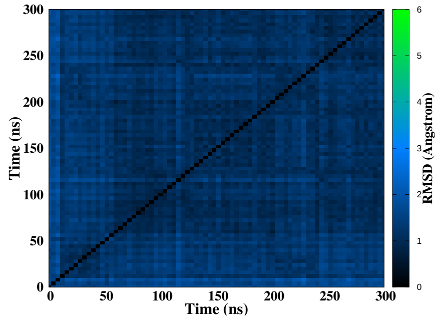


(I)

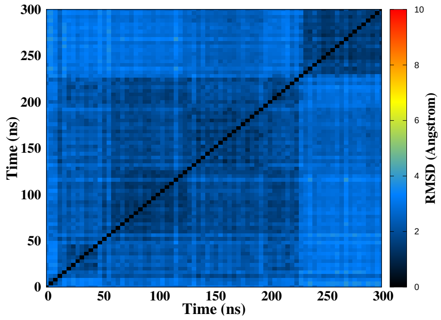


(J)

Figure S37 – Simulation of the NC damage at position 15-16, run 1. Snapshots of the initial and final MD simulation (A-B). RMSD of guanines forming the tetrads (C), distance between the tetrads (D), twist angle (E), angle between guanines for the first (F), second (G) and the third (H) tetrad, 2D-RMSD of guanines forming the tetrads (I) and the whole DNA (J).

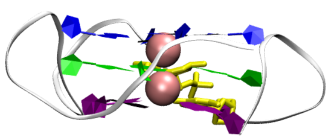

(A) 0 ns

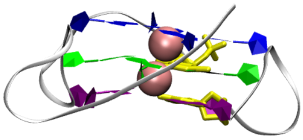

(B) 1000 ns

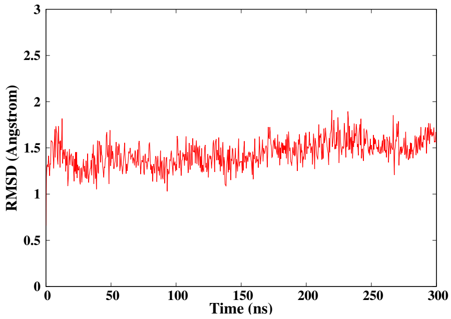


(C)

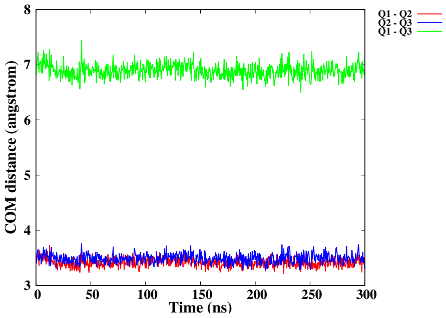


(D)

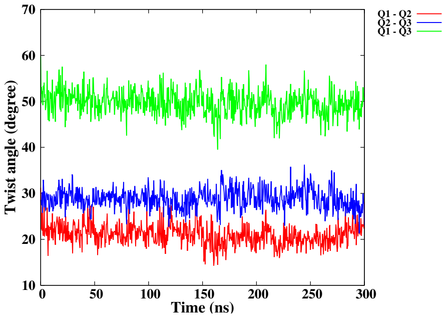


(E)

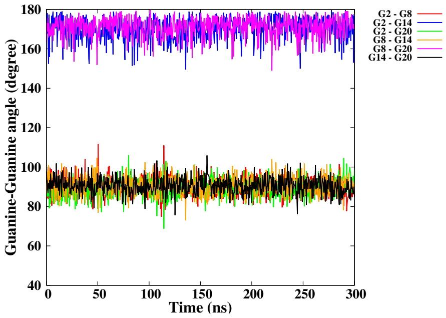


(F)

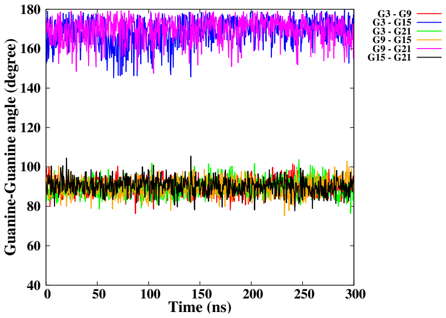


(G)

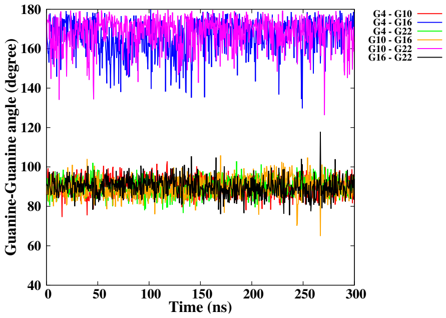


(H)

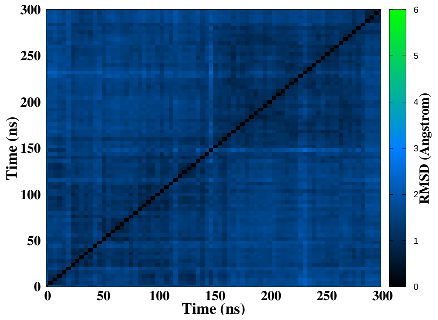


(I)

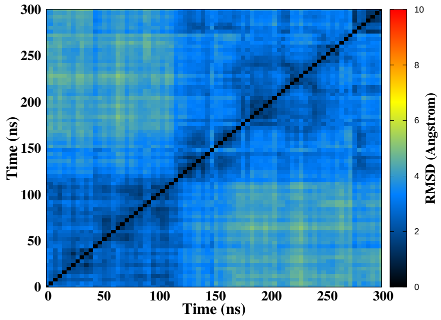


(J)

Figure S38 – Simulation of the NC damage at position 15-16, run 2. Snapshots of the initial and final MD simulation (A-B). RMSD of guanines forming the tetrads (C), distance between the tetrads (D), twist angle (E), angle between guanines for the first (F), second (G) and the third (H) tetrad, 2D-RMSD of guanines forming the tetrads (I) and the whole DNA (J).

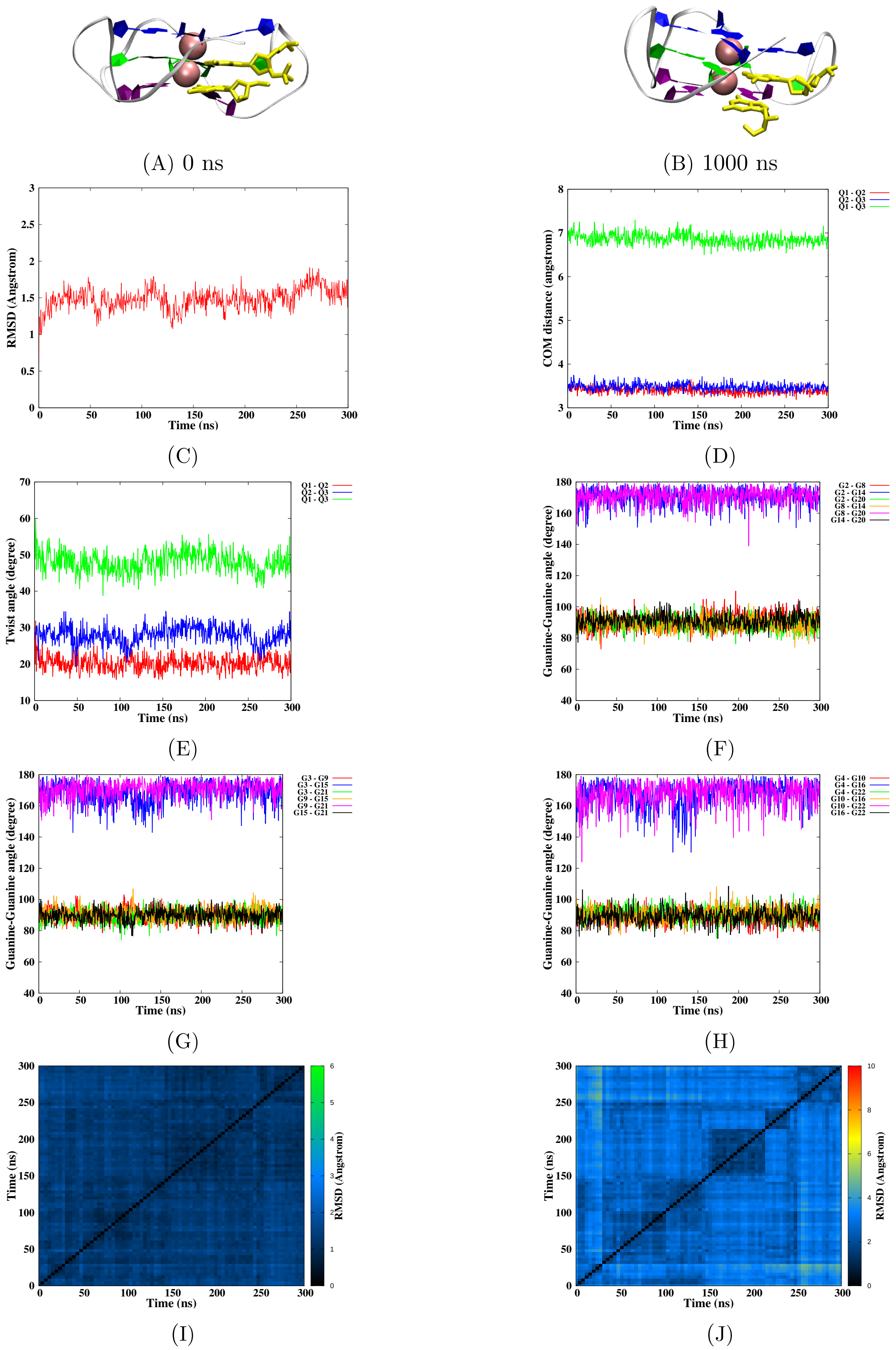


Figure S39 – Simulation of the NC damage at position 21-22, run 1. Snapshots of the initial and final MD simulation (A-B). RMSD of guanines forming the tetrads (C), distance between the tetrads (D), twist angle (E), angle between guanines for the first (F), second (G) and the third (H) tetrad, 2D-RMSD of guanines forming the tetrads (I) and the whole DNA (J).

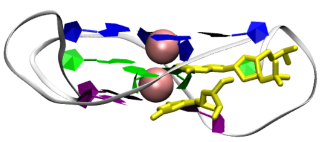

(A) 0 ns

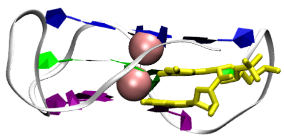

(B) 1000 ns

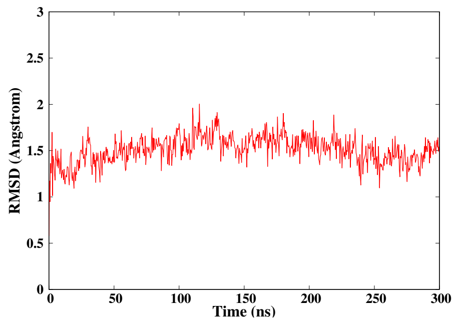


(C)

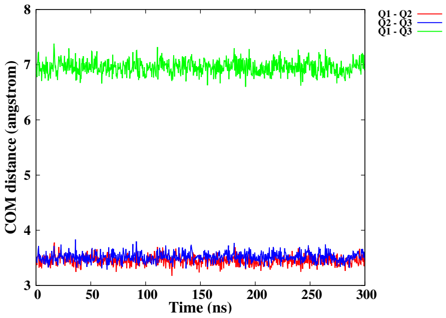


(D)

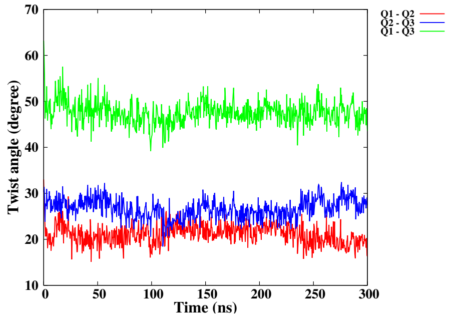


(E)

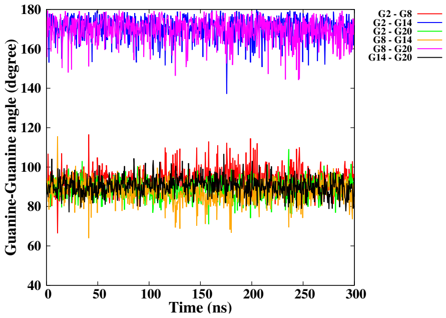


(F)

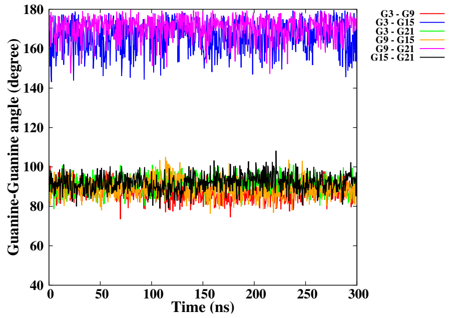


(G)

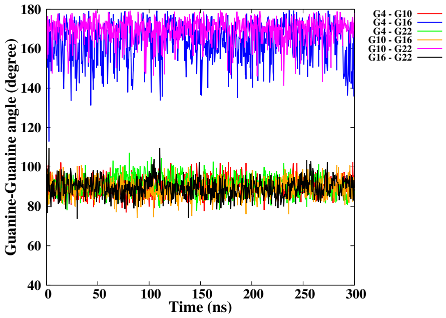


(H)

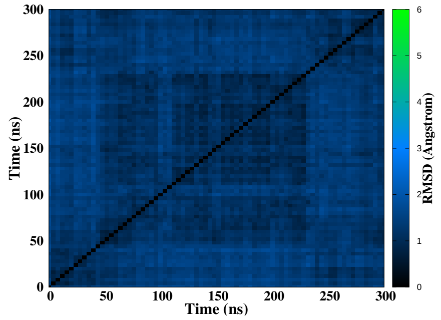


(I)

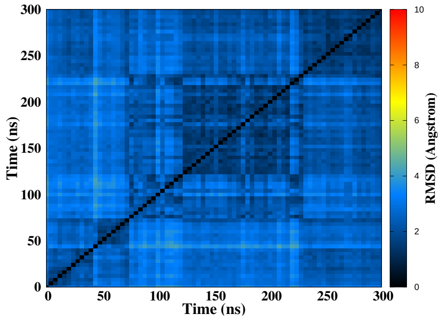


(J)

Figure S40 – Simulation of the NC damage at position 21-22, run 2. Snapshots of the initial and final MD simulation (A-B). RMSD of guanines forming the tetrads (C), distance between the tetrads (D), twist angle (E), angle between guanines for the first (F), second (G) and the third (H) tetrad, 2D-RMSD of guanines forming the tetrads (I) and the whole DNA (J).

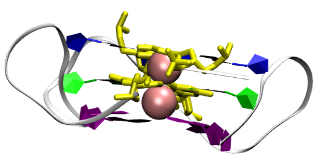

(A) 0 ns

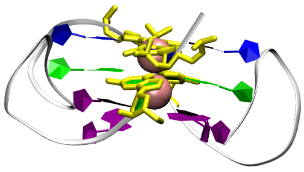

(B) 1000 ns

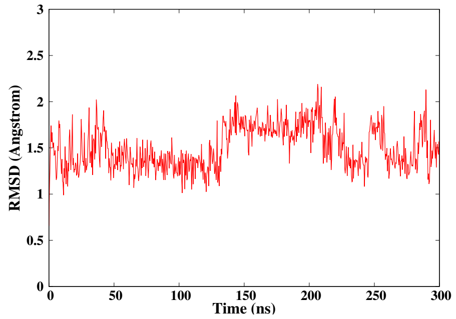


(C)

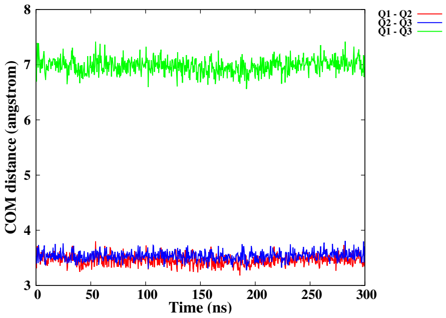


(D)

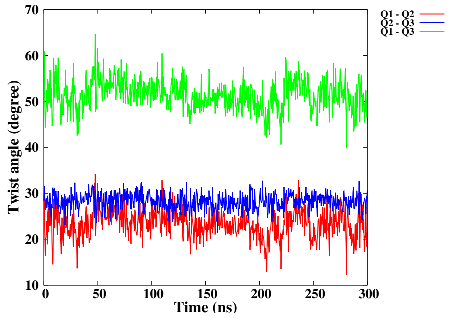


(E)

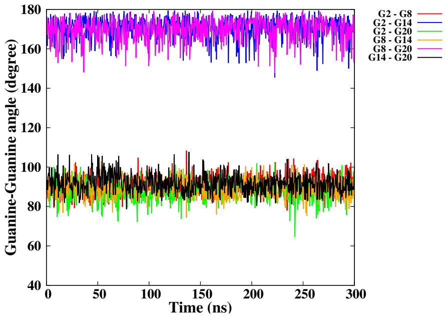


(F)

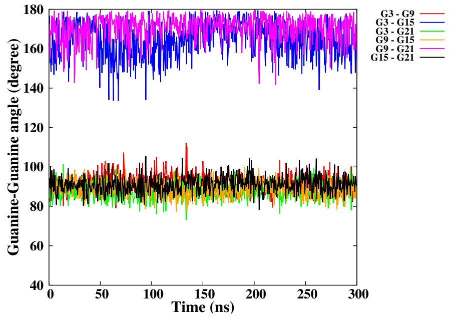


(G)

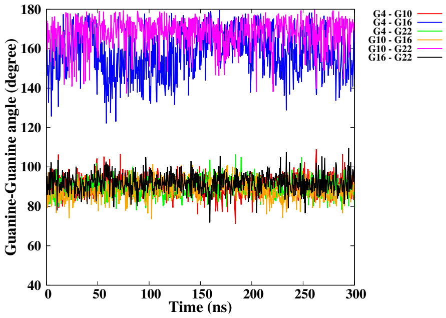


(H)

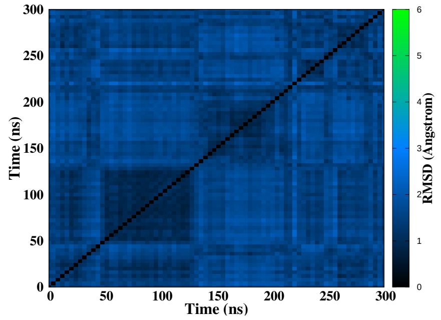


(I)

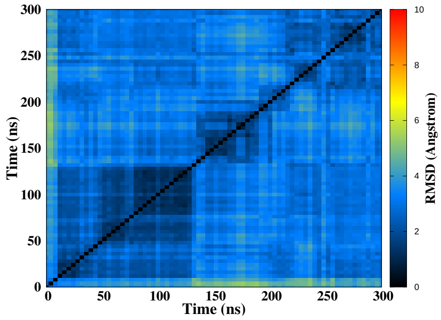


(J)

Figure S41 – Simulation of the NC damage at position 2-3/14-15, run 1. Snapshots of the initial and final MD simulation (A-B). RMSD of guanines forming the tetrads (C), distance between the tetrads (D), twist angle (E), angle between guanines for the first (F), second (G) and the third (H) tetrad, 2D-RMSD of guanines forming the tetrads (I) and the whole DNA (J).

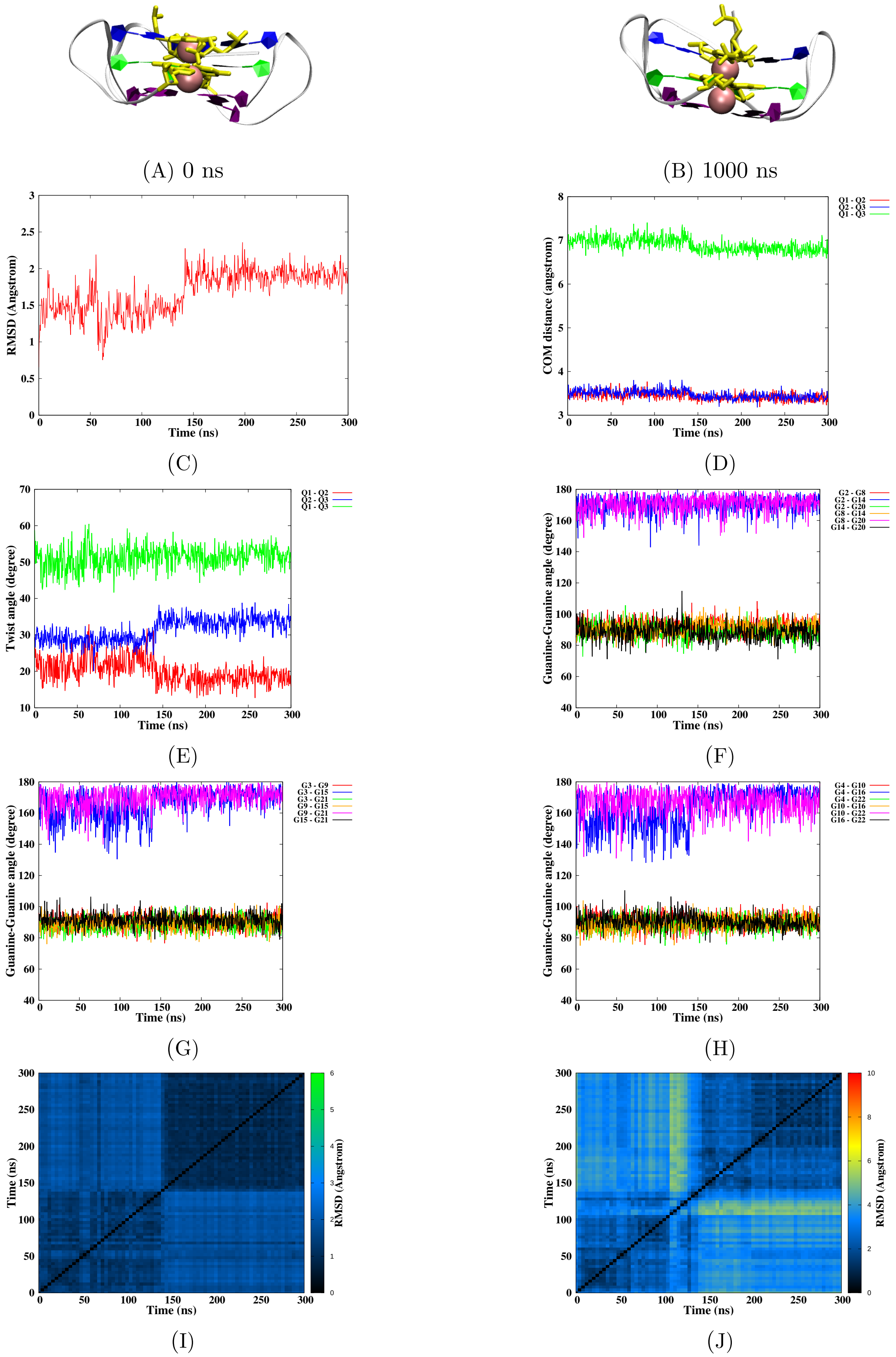


Figure S42 – Simulation of the NC damage at position 2-3/14-15, run 2. Snapshots of the initial and final MD simulation (A-B). RMSD of guanines forming the tetrads (C), distance between the tetrads (D), twist angle (E), angle between guanines for the first (F), second (G) and the third (H) tetrad, 2D-RMSD of guanines forming the tetrads (I) and the whole DNA (J).

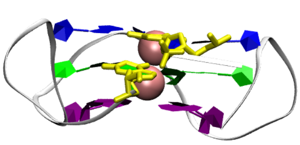

(A) 0 ns

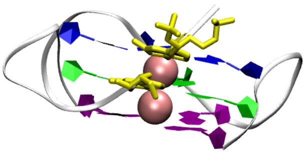

(B) 1000 ns

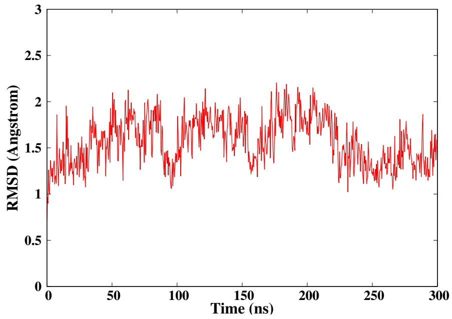


(C)

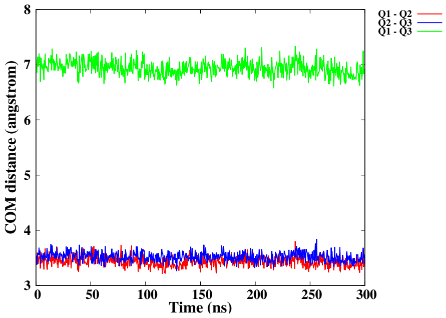


(D)

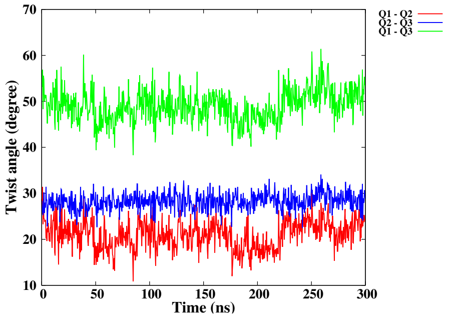


(E)

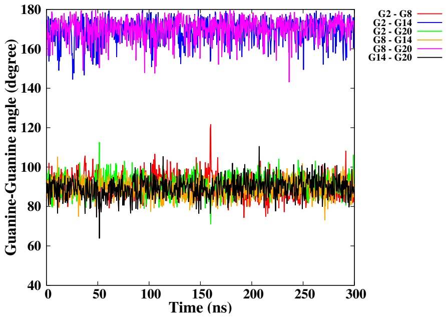


(F)

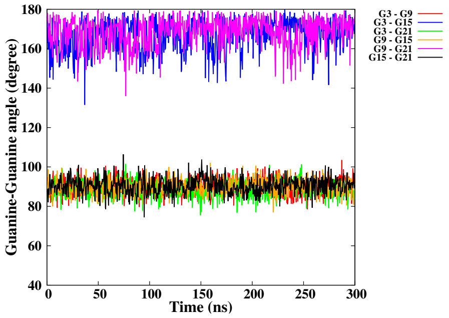


(G)

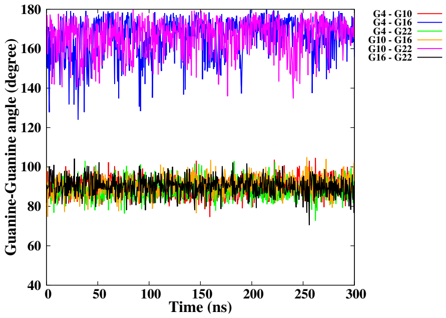


(H)

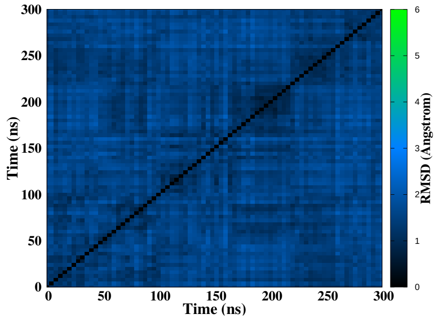


(I)

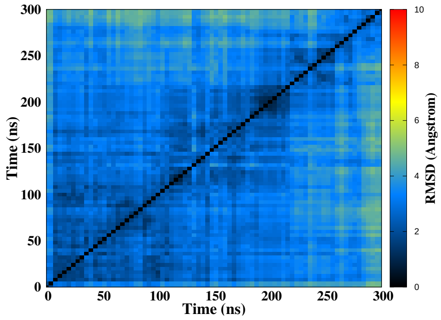


(J)

Figure S43 – Simulation of the NC damage at position 2-3, run 1. Snapshots of the initial and final MD simulation (A-B). RMSD of guanines forming the tetrads (C), distance between the tetrads (D), twist angle (E), angle between guanines for the first (F), second (G) and the third (H) tetrad, 2D-RMSD of guanines forming the tetrads (I) and the whole DNA (J).

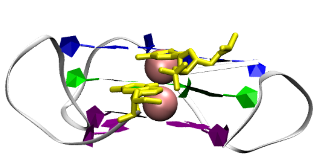

(A) 0 ns

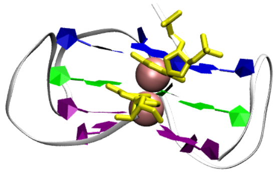

(B) 1000 ns

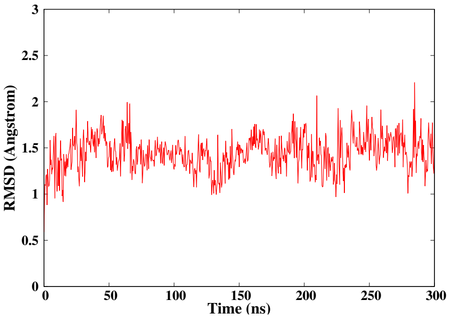


(C)

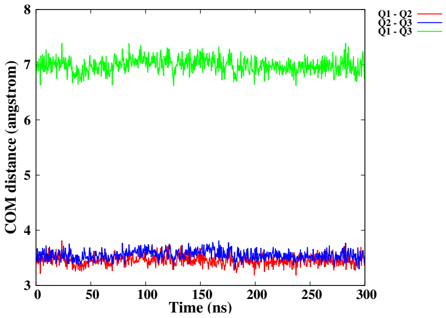


(D)

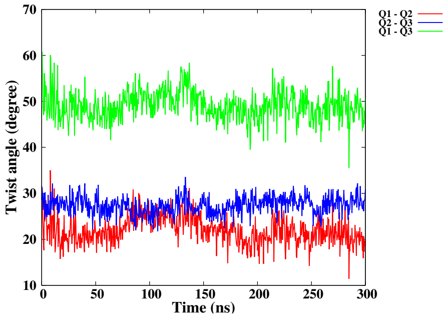


(E)

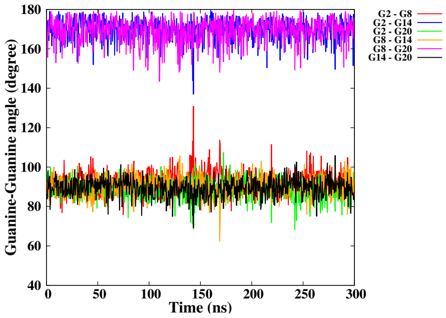


(F)

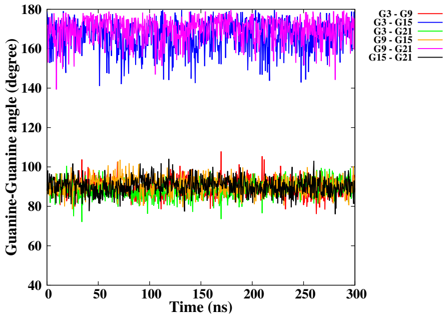


(G)

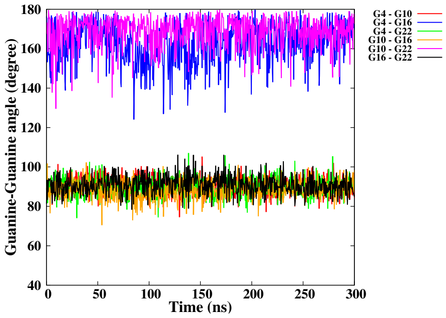


(H)

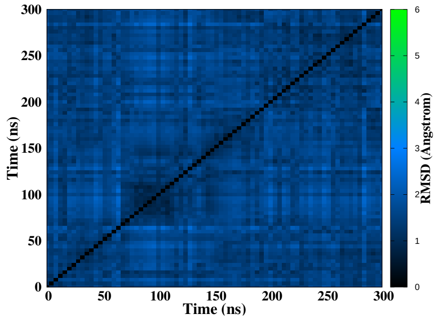


(I)

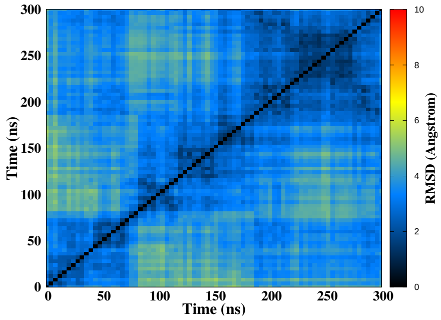


(J)

Figure S44 – Simulation of the NC damage at position 2-3, run 2. Snapshots of the initial and final MD simulation (A-B). RMSD of guanines forming the tetrads (C), distance between the tetrads (D), twist angle (E), angle between guanines for the first (F), second (G) and the third (H) tetrad, 2D-RMSD of guanines forming the tetrads (I) and the whole DNA (J).

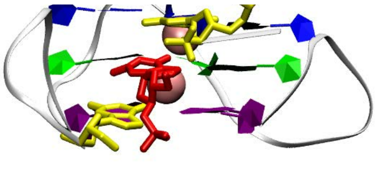

(A) 0 ns

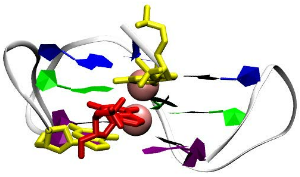

(B) 1000 ns

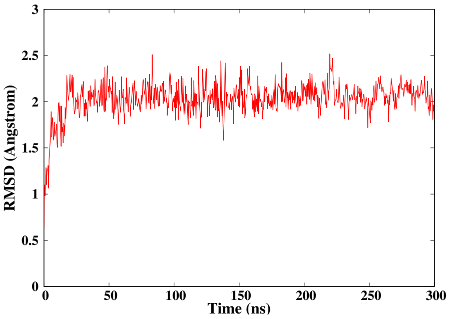


(C)

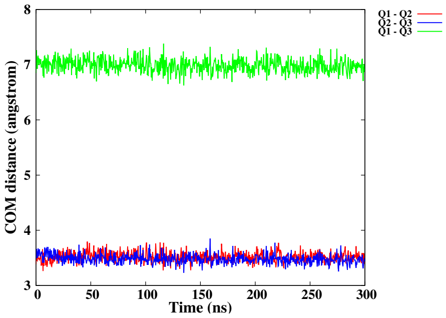


(D)

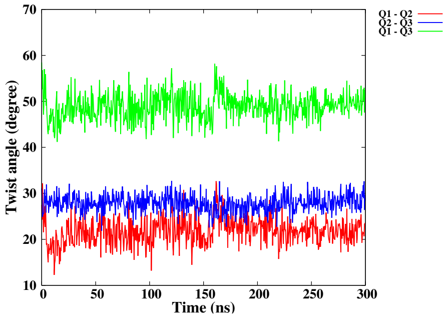


(E)

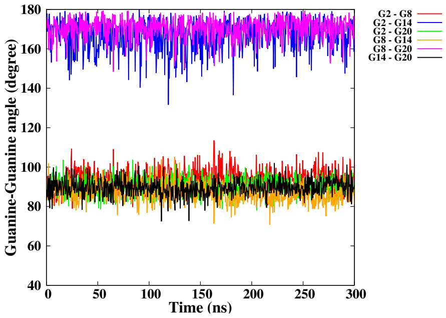


(F)

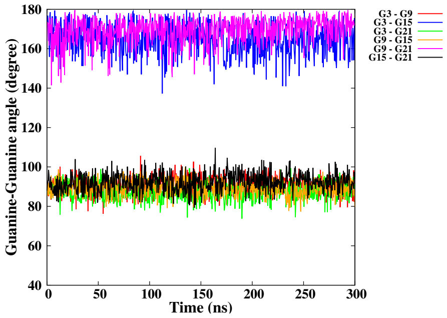


(G)

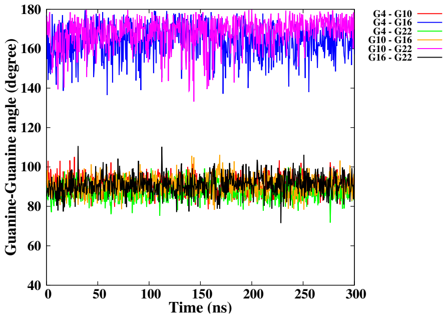


(H)

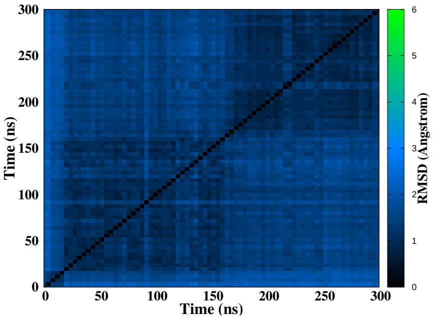


(I)

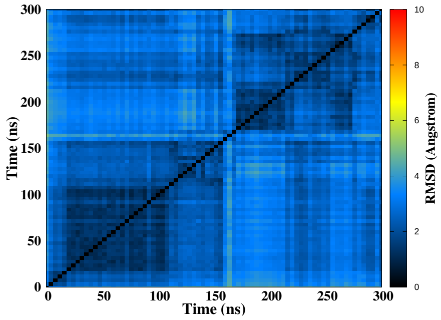


(J)

Figure S45 – Simulation of the NC damage at position 2-3-4, run 1. Snapshots of the initial and final MD simulation (A-B). RMSD of guanines forming the tetrads (C), distance between the tetrads (D), twist angle (E), angle between guanines for the first (F), second (G) and the third (H) tetrad, 2D-RMSD of guanines forming the tetrads (I) and the whole DNA (J).

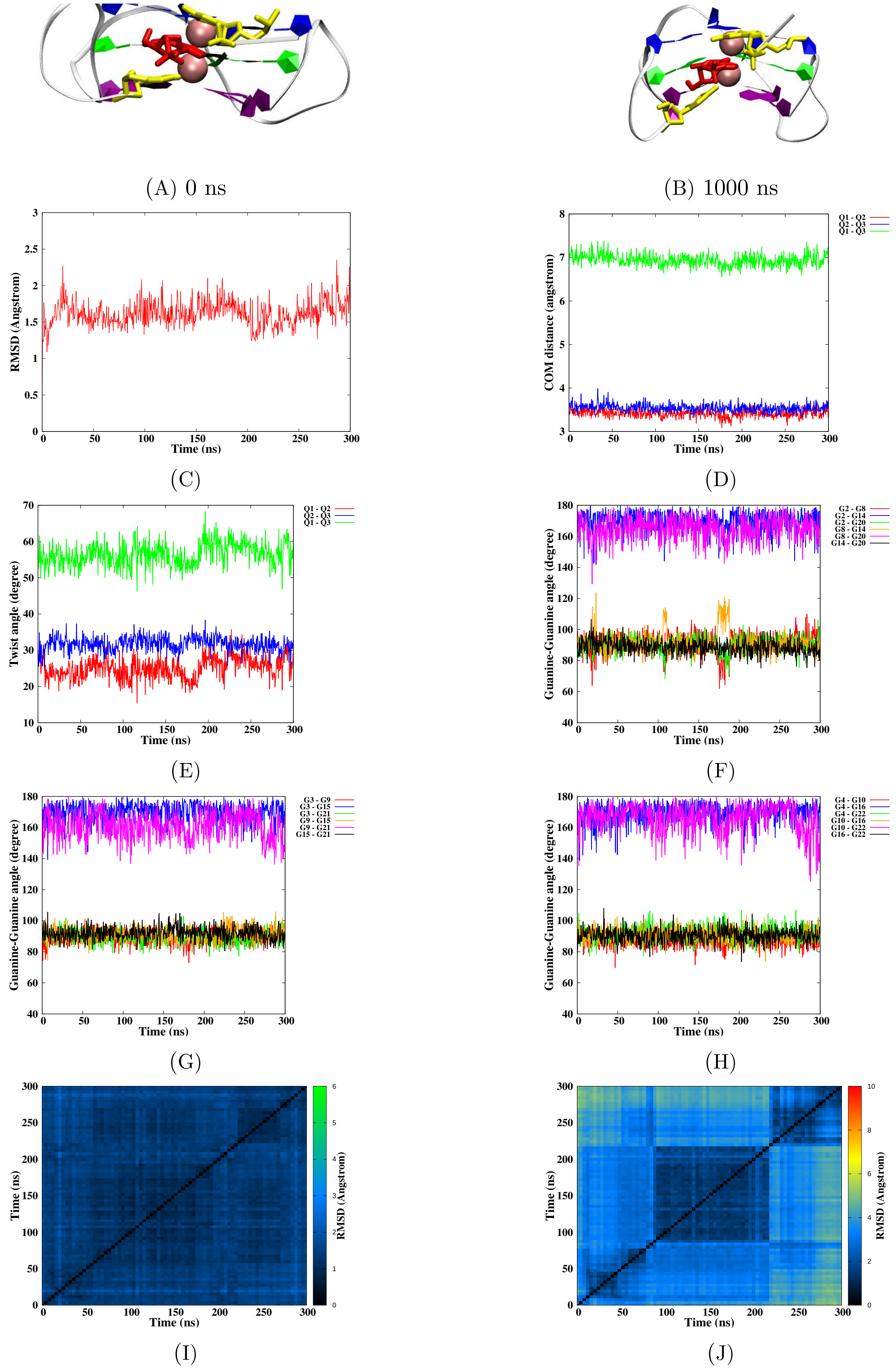


Figure S46 – Simulation of the NC damage at position 2-3-4, run 2. Snapshots of the initial and final MD simulation (A-B). RMSD of guanines forming the tetrads (C), distance between the tetrads (D), twist angle (E), angle between guanines for the first (F), second (G) and the third (H) tetrad, 2D-RMSD of guanines forming the tetrads (I) and the whole DNA (J).

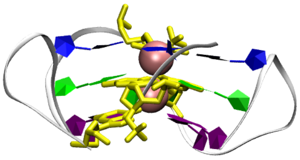

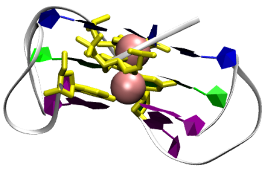

(A) 0 ns

(B) 1000 ns

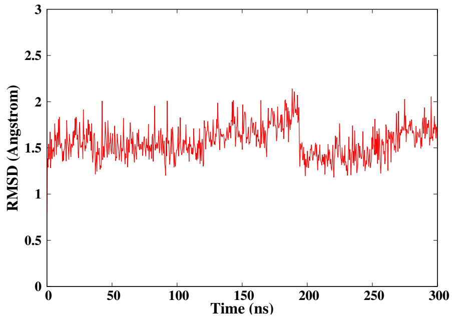


(C)

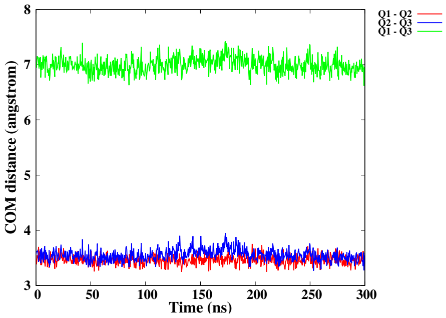


(D)

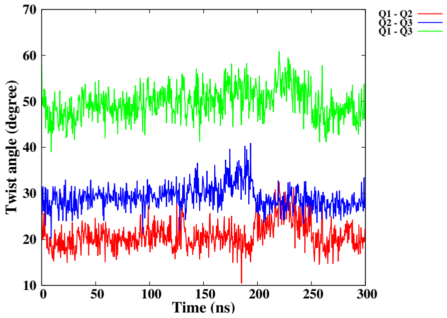


(E)

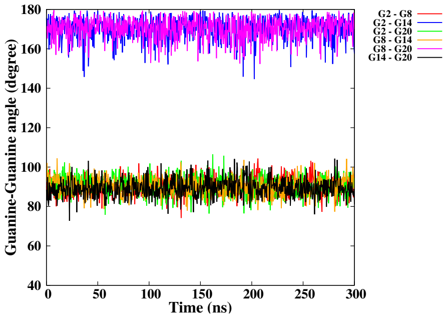


(F)

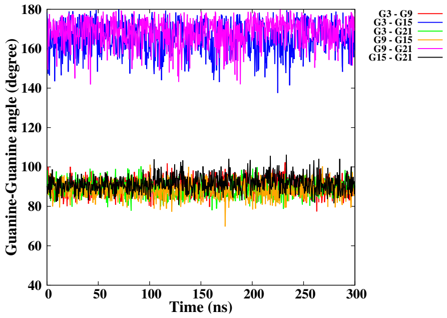


(G)

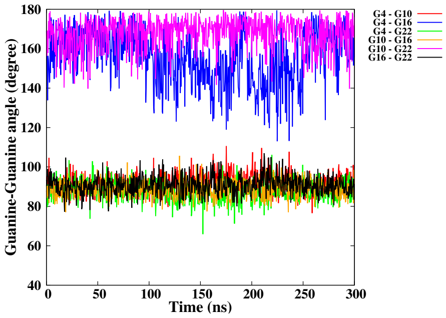


(H)

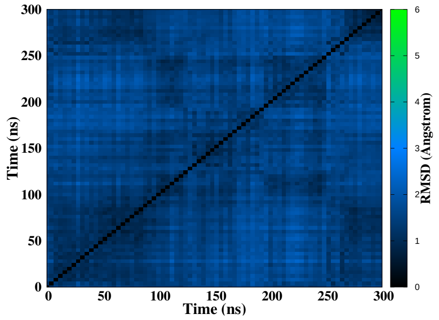


(I)

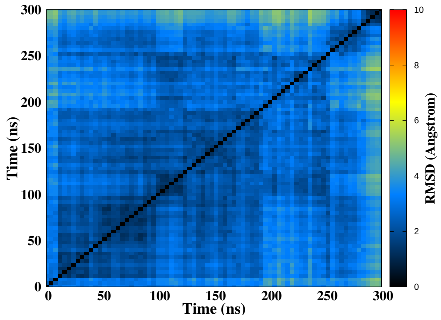


(J)

Figure S47 – Simulation of the NC damage at position 3-4/14-15, run 1. Snapshots of the initial and final MD simulation (A-B). RMSD of guanines forming the tetrads (C), distance between the tetrads (D), twist angle (E), angle between guanines for the first (F), second (G) and the third (H) tetrad, 2D-RMSD of guanines forming the tetrads (I) and the whole DNA (J).

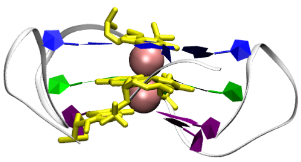

(A) 0 ns

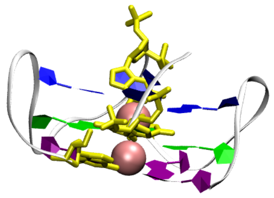

(B) 1000 ns

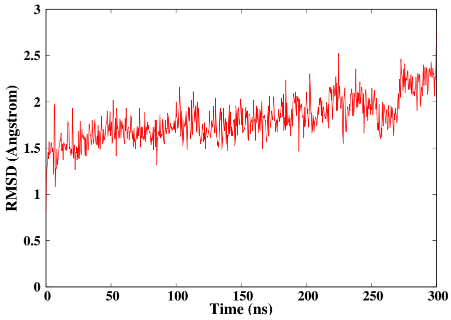


(C)

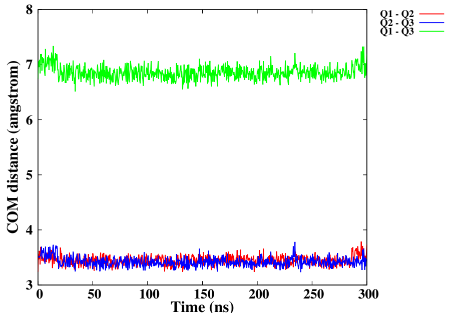


(D)

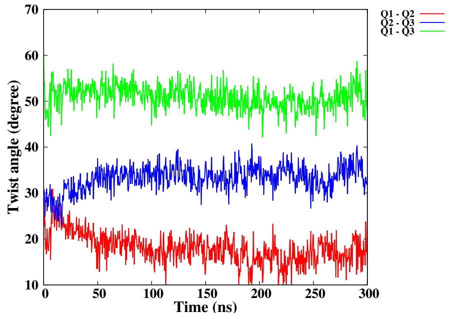


(E)

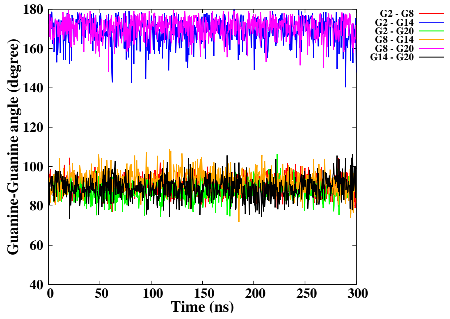


(F)

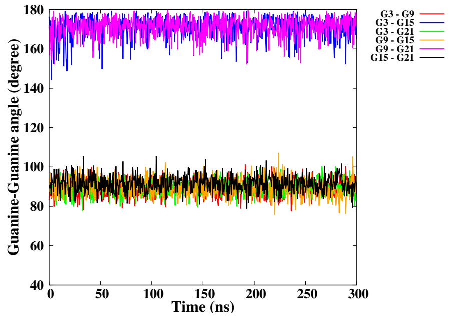


(G)

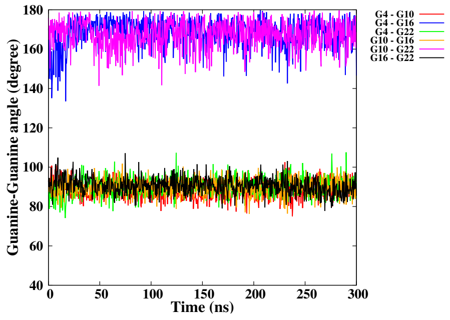


(H)

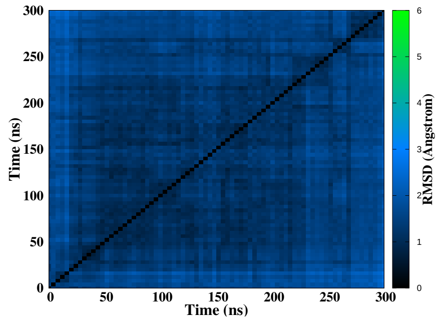


(I)

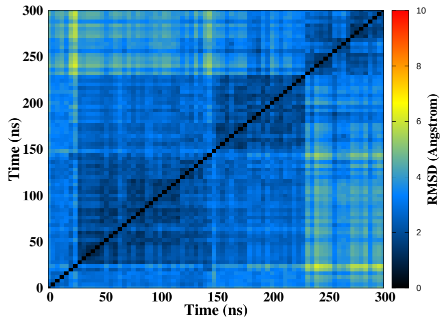


(J)

Figure S48 – Simulation of the NC damage at position 3-4/14-15, run 2. Snapshots of the initial and final MD simulation (A-B). RMSD of guanines forming the tetrads (C), distance between the tetrads (D), twist angle (E), angle between guanines for the first (F), second (G) and the third (H) tetrad, 2D-RMSD of guanines forming the tetrads (I) and the whole DNA (J).

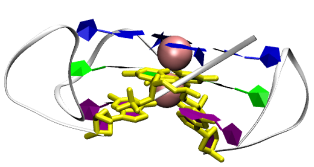

(A) 0 ns

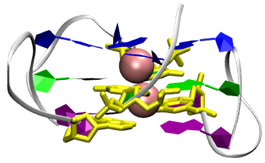

(B) 1000 ns

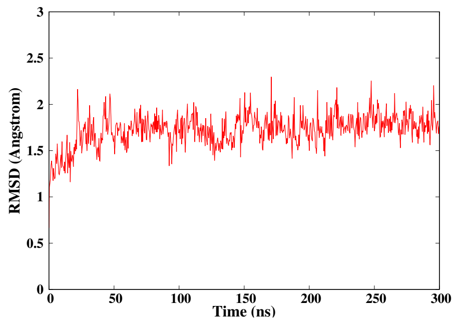


(C)

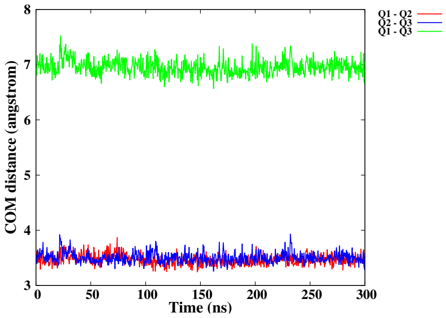


(D)

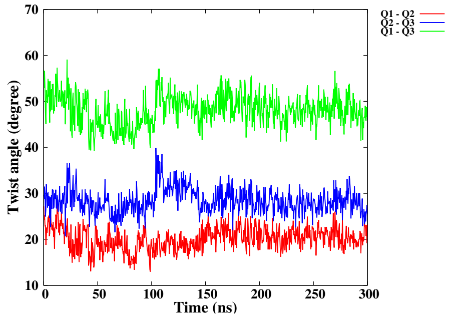


(E)

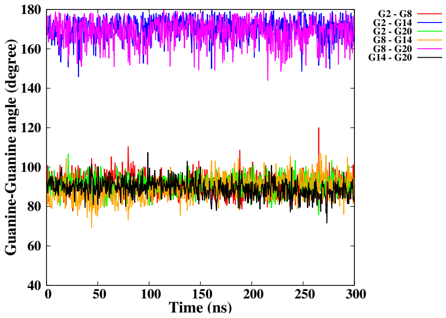


(F)

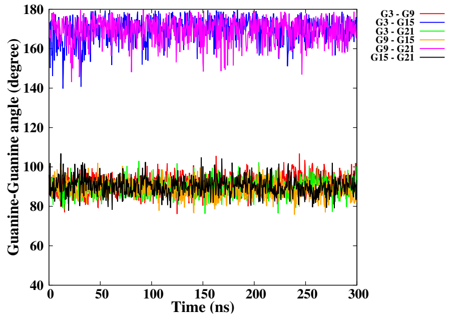


(G)

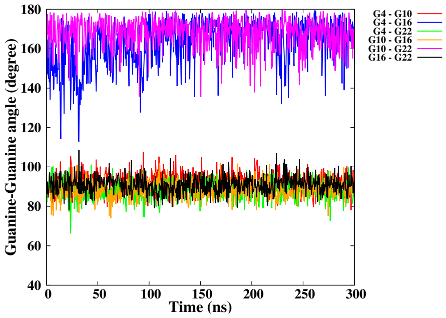


(H)

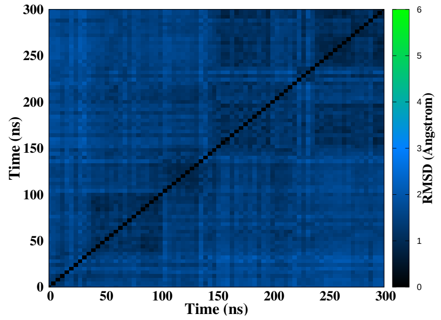


(I)

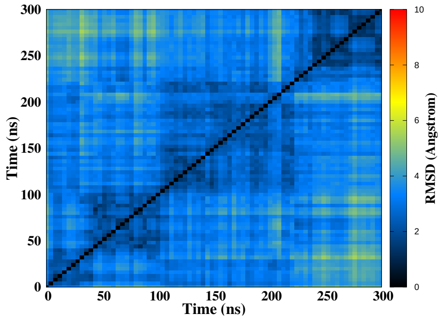


(J)

Figure S49 – Simulation of the NC damage at position 3-4/15-16, run 1. Snapshots of the initial and final MD simulation (A-B). RMSD of guanines forming the tetrads (C), distance between the tetrads (D), twist angle (E), angle between guanines for the first (F), second (G) and the third (H) tetrad, 2D-RMSD of guanines forming the tetrads (I) and the whole DNA (J).

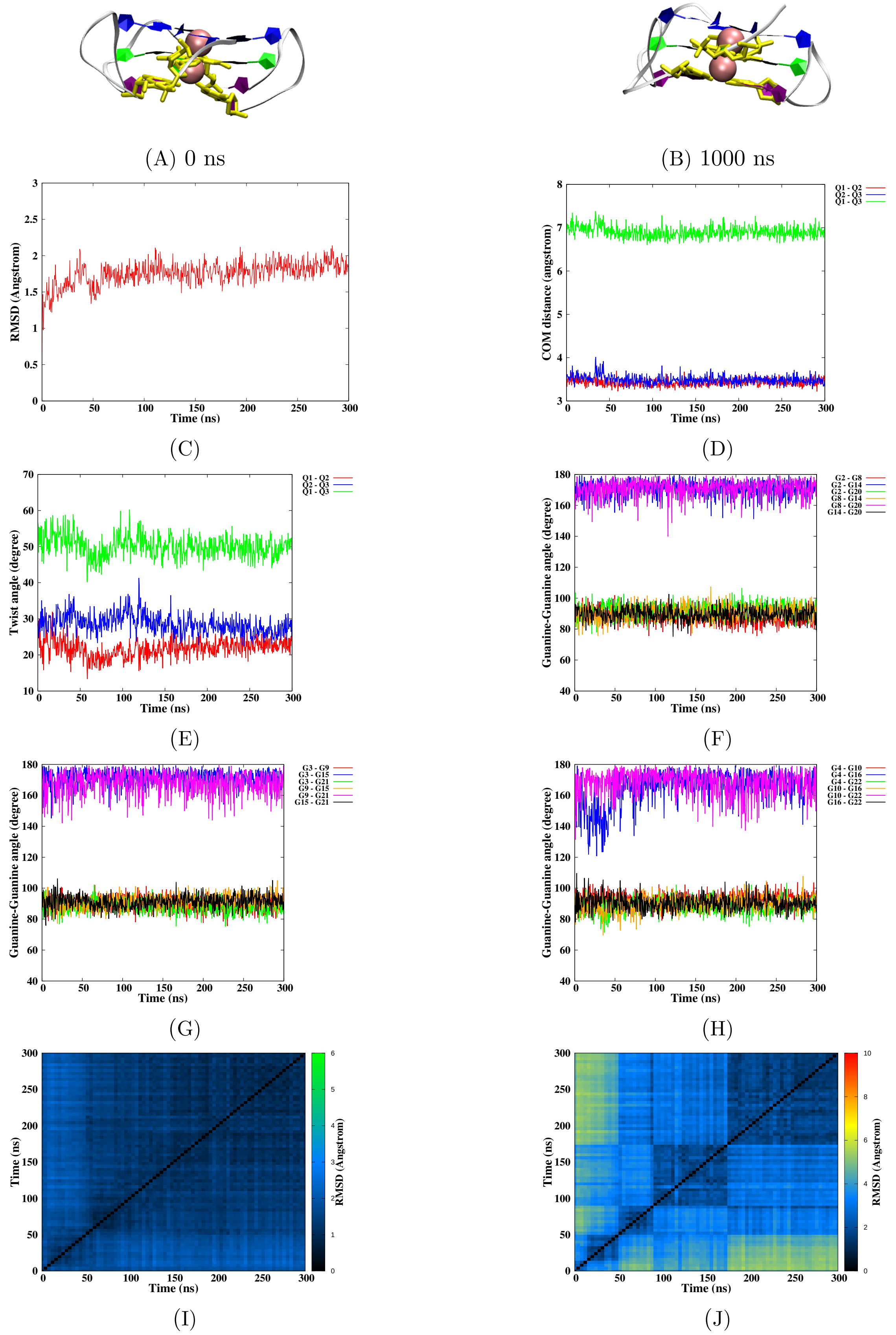


Figure S50 – Simulation of the NC damage at position 3-4/15-16, run 2. Snapshots of the initial and final MD simulation (A-B). RMSD of guanines forming the tetrads (C), distance between the tetrads (D), twist angle (E), angle between guanines for the first (F), second (G) and the third (H) tetrad, 2D-RMSD of guanines forming the tetrads (I) and the whole DNA (J).

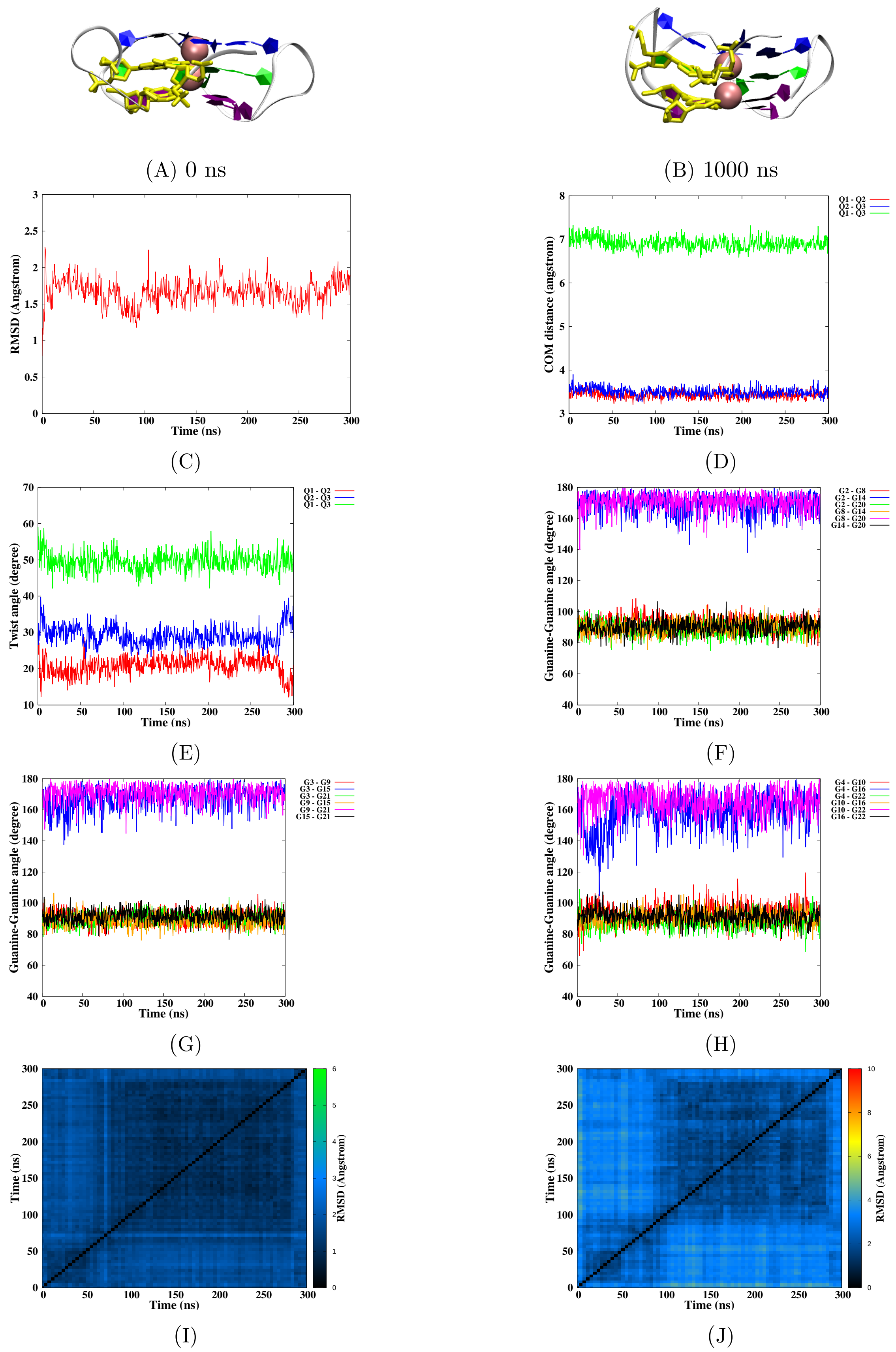


Figure S51 – Simulation of the NC damage at position 3-4/9-10, run 1. Snapshots of the initial and final MD simulation (A-B). RMSD of guanines forming the tetrads (C), distance between the tetrads (D), twist angle (E), angle between guanines for the first (F), second (G) and the third (H) tetrad, 2D-RMSD of guanines forming the tetrads (I) and the whole DNA (J).

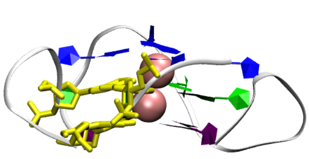

(A) 0 ns

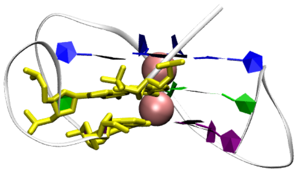

(B) 1000 ns

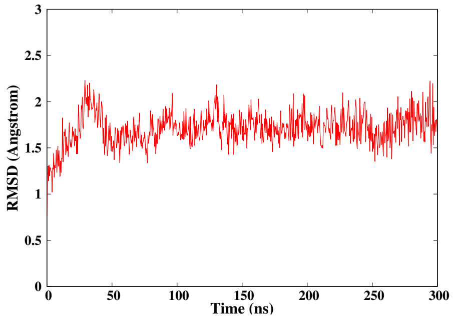


(C)

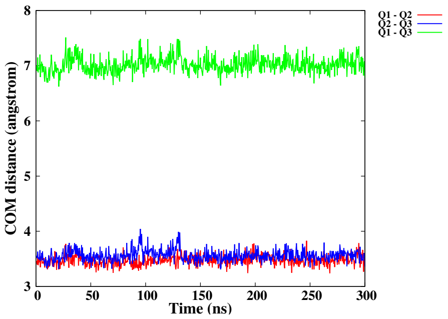


(D)

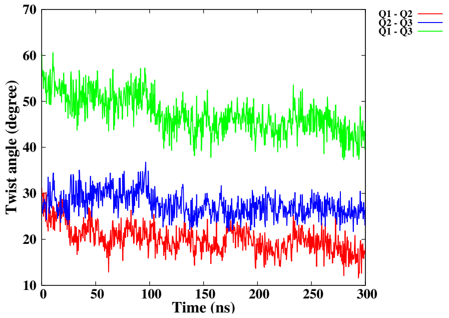


(E)

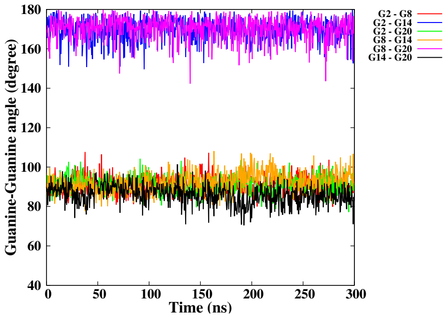


(F)

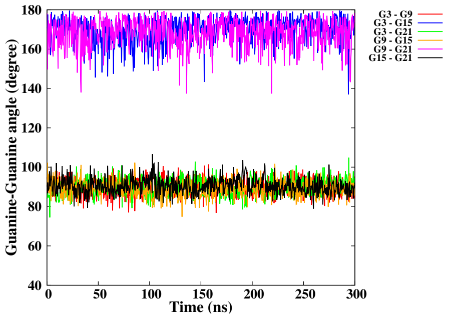


(G)

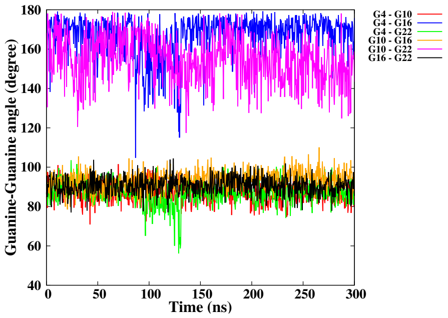


(H)

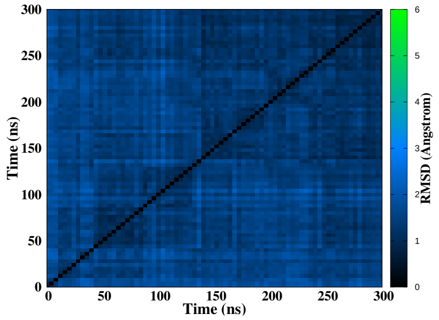


(I)

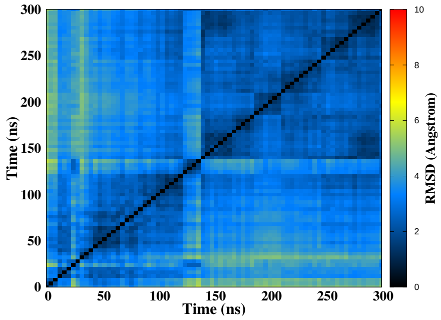


(J)

Figure S52 – Simulation of the NC damage at position 3-4/9-10, run 2. Snapshots of the initial and final MD simulation (A-B). RMSD of guanines forming the tetrads (C), distance between the tetrads (D), twist angle (E), angle between guanines for the first (F), second (G) and the third (H) tetrad, 2D-RMSD of guanines forming the tetrads (I) and the whole DNA (J).

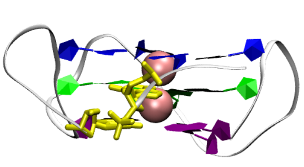

(A) 0 ns

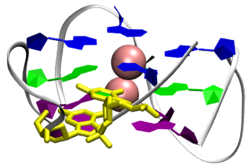

(B) 1000 ns

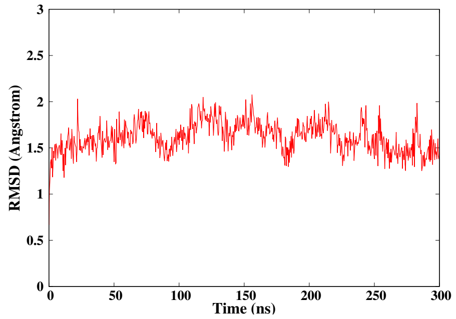


(C)

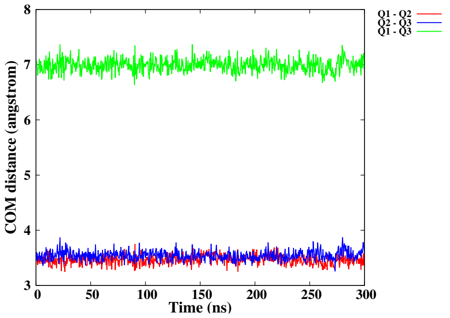


(D)

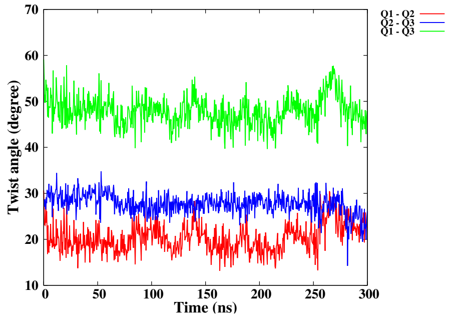


(E)

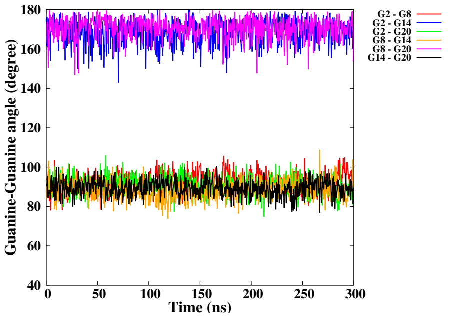


(F)

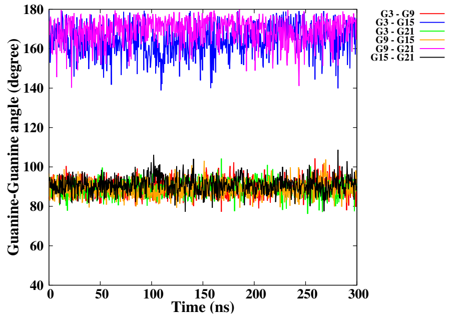


(G)

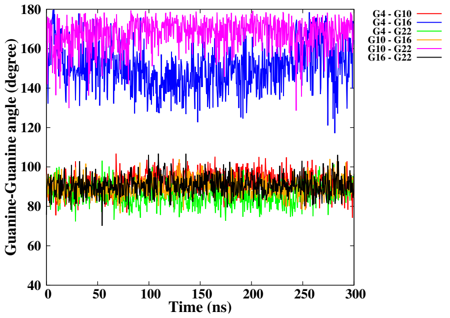


(H)

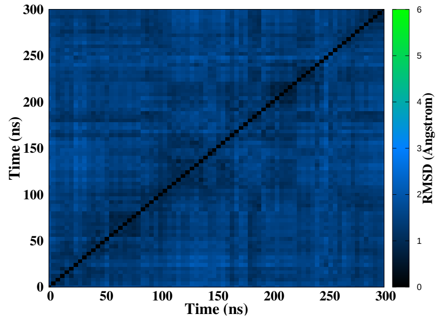


(I)

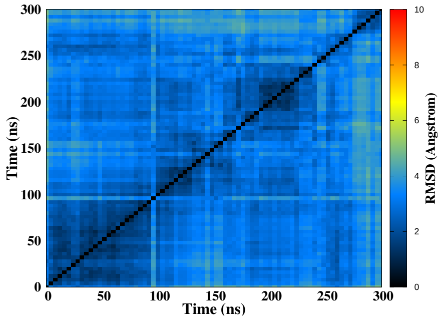


(J)

Figure S53 – Simulation of the NC damage at position 3-4, run 1. Snapshots of the initial and final MD simulation (A-B). RMSD of guanines forming the tetrads (C), distance between the tetrads (D), twist angle (E), angle between guanines for the first (F), second (G) and the third (H) tetrad, 2D-RMSD of guanines forming the tetrads (I) and the whole DNA (J).

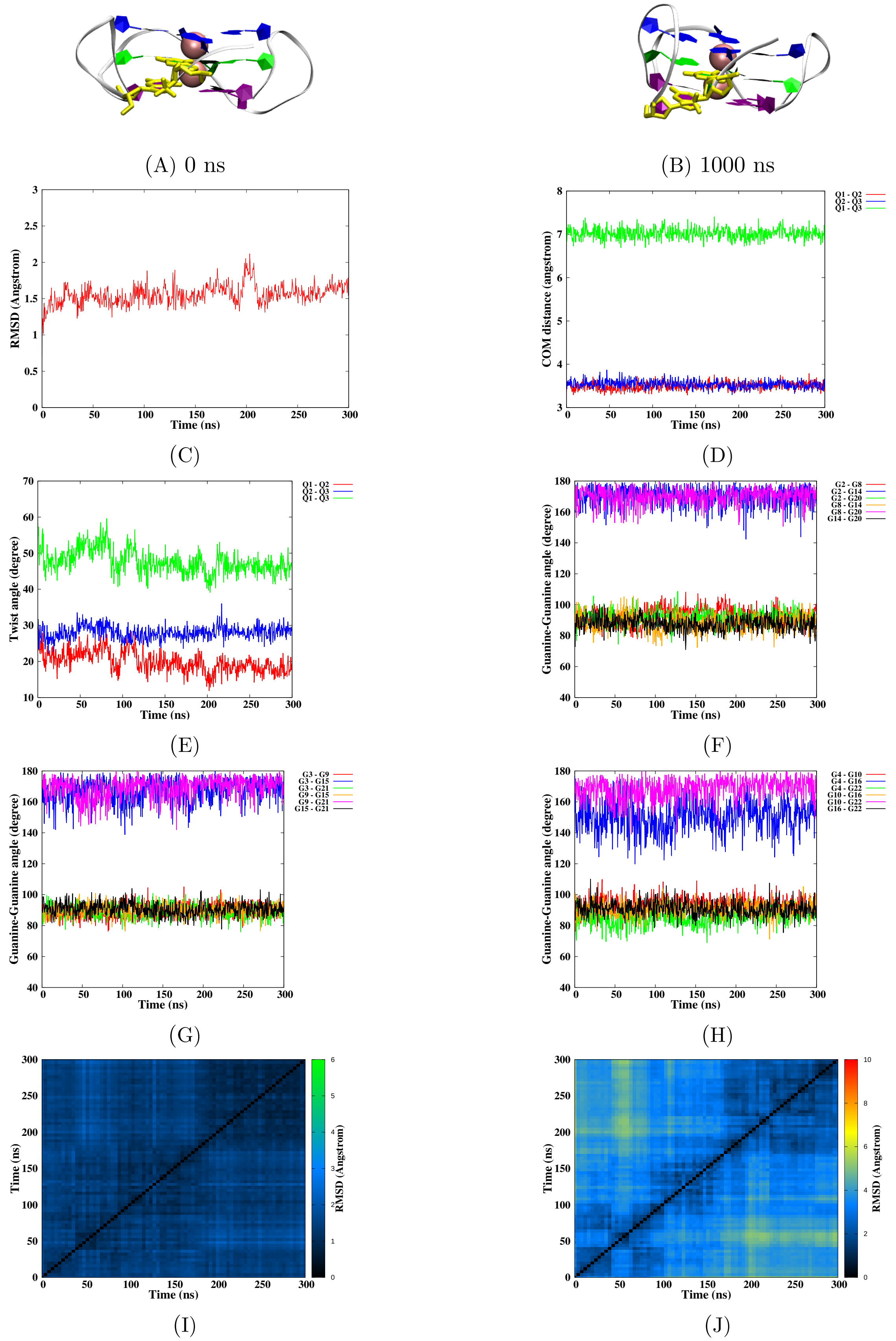


Figure S54 – Simulation of the NC damage at position 3-4, run 2. Snapshots of the initial and final MD simulation (A-B). RMSD of guanines forming the tetrads (C), distance between the tetrads (D), twist angle (E), angle between guanines for the first (F), second (G) and the third (H) tetrad, 2D-RMSD of guanines forming the tetrads (I) and the whole DNA (J).

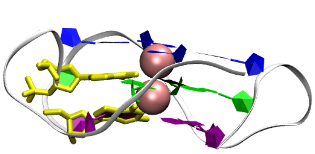

(A) 0 ns

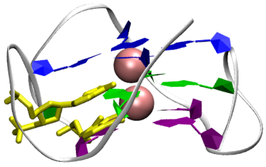

(B) 1000 ns

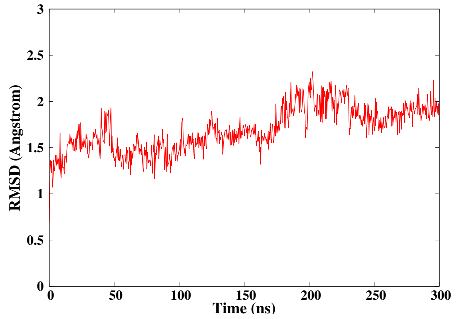


(C)

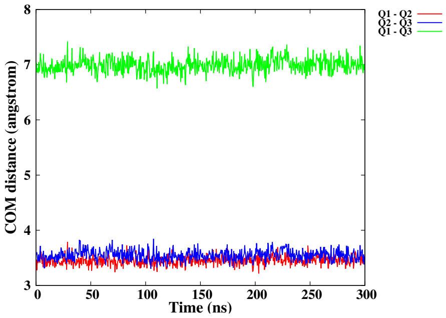


(D)

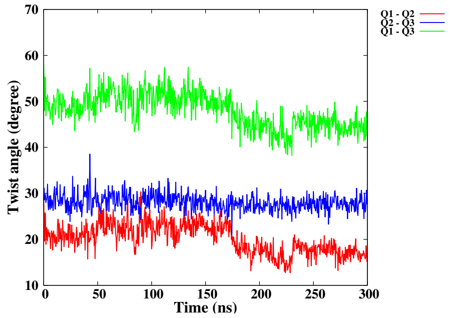


(E)

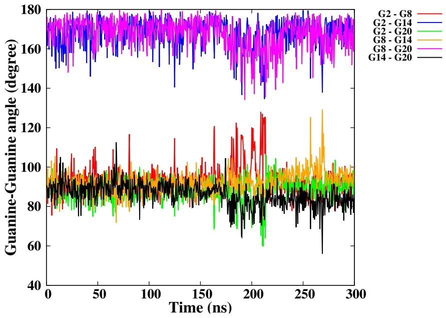


(F)

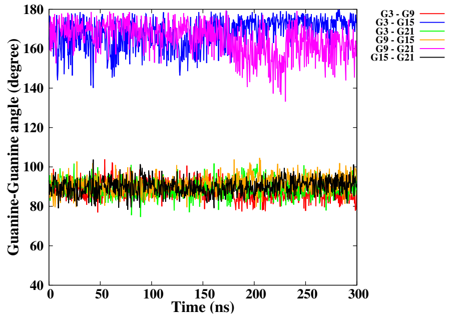


(G)

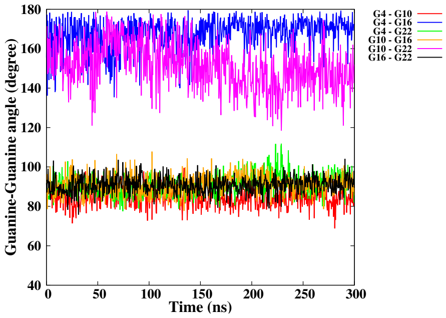


(H)

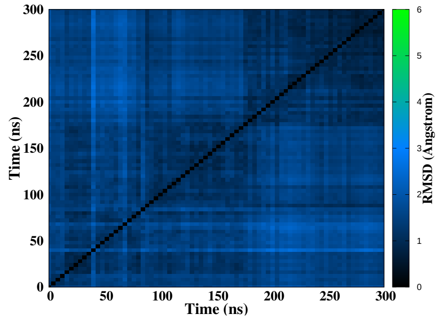


(I)

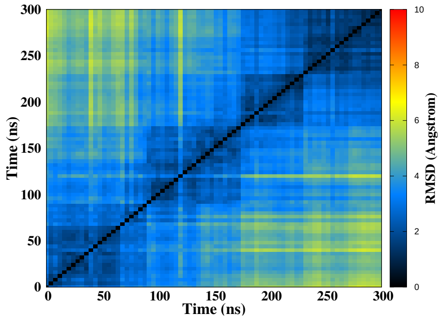


(J)

Figure S55 – Simulation of the NC damage at position 9-10, run 1. Snapshots of the initial and final MD simulation (A-B). RMSD of guanines forming the tetrads (C), distance between the tetrads (D), twist angle (E), angle between guanines for the first (F), second (G) and the third (H) tetrad, 2D-RMSD of guanines forming the tetrads (I) and the whole DNA (J).

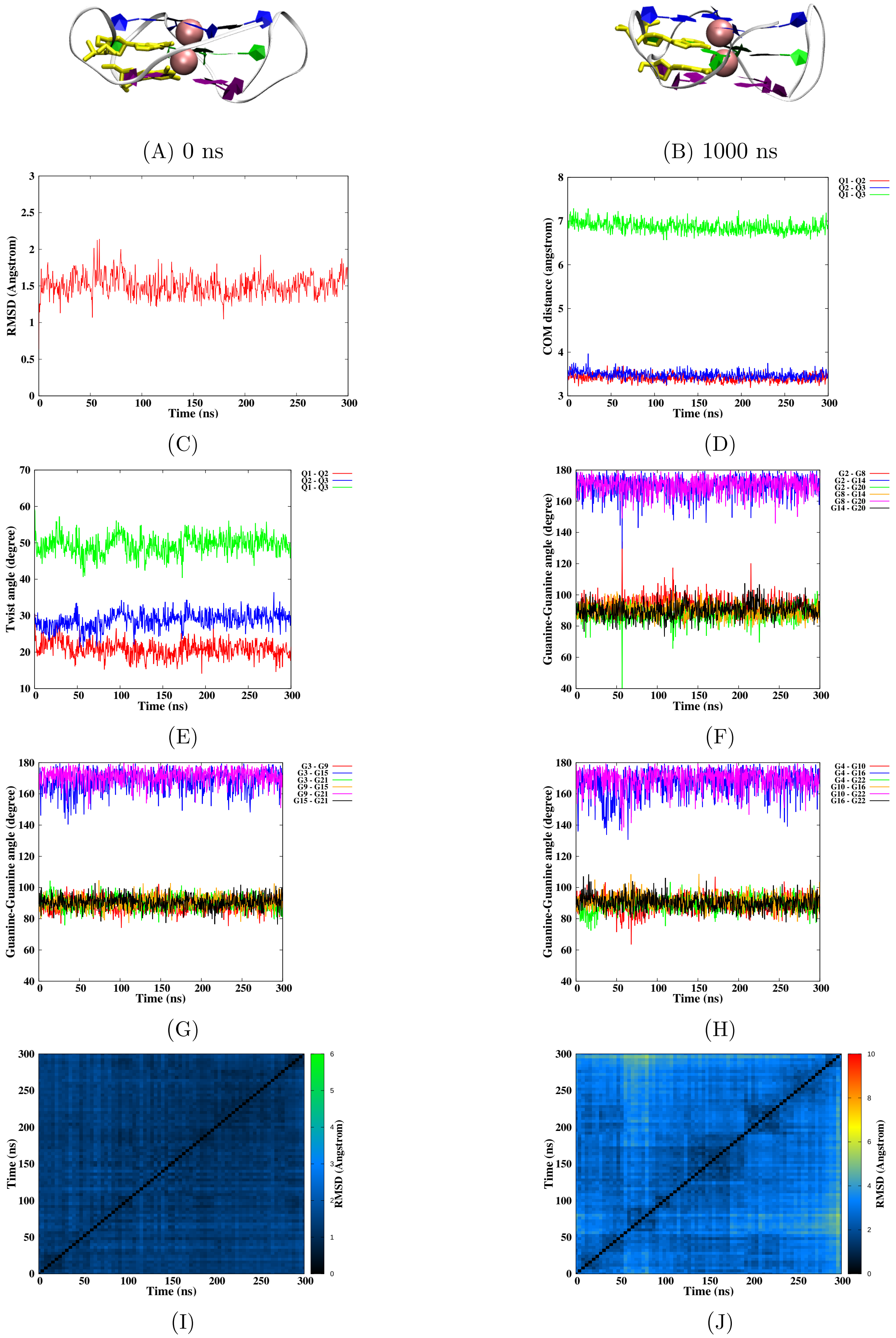


Figure S56 – Simulation of the NC damage at position 9-10, run 2. Snapshots of the initial and final MD simulation (A-B). RMSD of guanines forming the tetrads (C), distance between the tetrads (D), twist angle (E), angle between guanines for the first (F), second (G) and the third (H) tetrad, 2D-RMSD of guanines forming the tetrads (I) and the whole DNA (J).

| Strand break Type | Positions | Run 1 $\Psi_{Q1-Q2}$ | Run 1 $\Psi_{Q2-Q3}$ | Run 1 $\Psi_{Q1-Q3}$ | Run 2 $\Psi_{Q1-Q2}$ | Run 2 $\Psi_{Q2-Q3}$ | Run 2 $\Psi_{Q1-Q3}$ |
|---|---|---|---|---|---|---|---|
| Native | | | | | | | |
| No Strand Breaks | | 19.19 ± 2.22 | 28.55 ± 1.93 | 47.57 ± 2.62 | 19.35 ± 2.45 | 27.85 ± 2.02 | 47.02 ± 2.92 |
| Loops - Canonical strand breaks | | | | | | | |
| **Single SB** | 12-13 | 19.14 ± 2.13 | 28.22 ± 1.69 | 47.18 ± 2.70 | 18.39 ± 2.41 | 27.58 ± 1.87 | 46.05 ± 3.03 |
| | 6-7 | 18.39 ± 2.41 | 28.19 ± 1.69 | 46.40 ± 2.89 | 18.32 ± 2.42 | 27.72 ± 1.80 | 45.86 ± 2.91 |
| **Double SB** | 6-7/12-13 | 18.48 ± 2.62 | 27.41 ± 1.82 | 45.72 ± 2.87 | 18.42 ± 2.11 | 27.19 ± 2.39 | 45.41 ± 3.43 |
| Tetrads - Canonical strand breaks | | | | | | | |
| **Single SB** | 2-3 | 20.89 ± 2.57 | 27.79 ± 2.11 | 48.48 ± 2.77 | 20.15 ± 3.29 | 27.97 ± 2.08 | 47.87 ± 3.40 |
| | 3-4 | 21.44 ± 1.92 | 29.43 ± 2.46 | 50.64 ± 2.76 | 20.39 ± 1.97 | 27.70 ± 1.88 | 47.89 ± 2.50 |
| | 9-10 | 21.48 ± 2.12 | 27.20 ± 1.72 | 48.49 ± 2.56 | 21.63 ± 2.18 | 26.86 ± 1.98 | 48.30 ± 2.89 |
| | 14-15 | 19.50 ± 2.45 | 30.64 ± 2.00 | 49.95 ± 2.62 | 26.23 ± 6.11 | 27.76 ± 2.40 | 50.44 ± 5.24 |
| | 15-16 | 22.24 ± 2.26 | 27.75 ± 1.68 | 49.77 ± 2.53 | 21.93 ± 2.33 | 27.04 ± 1.85 | 48.78 ± 2.58 |
| | 21-22 | 19.87 ± 2.51 | 27.85 ± 2.19 | 47.50 ± 3.11 | 22.17 ± 2.96 | 27.26 ± 2.28 | 49.22 ± 3.32 |
| **Double SB** | 2-3/14-15 | 17.85 ± 3.37 | 31.99 ± 2.27 | 49.55 ± 3.49 | 17.37 ± 2.94 | 31.84 ± 1.84 | 49.03 ± 2.92 |
| | 2-3-4 | 20.68 ± 2.85 | 29.50 ± 2.41 | 49.93 ± 3.54 | 20.17 ± 2.25 | 27.90 ± 2.19 | 47.78 ± 3.11 |
| | 3-4/9-10 | 21.76 ± 2.44 | 27.03 ± 1.80 | 48.58 ± 2.79 | 20.72 ± 1.97 | 28.38 ± 2.15 | 48.92 ± 2.68 |
| | 3-4/14-15 | 22.19 ± 3.06 | 26.98 ± 2.50 | 48.93 ± 3.59 | 21.11 ± 2.67 | 28.56 ± 2.13 | 49.46 ± 2.81 |
| | 3-4/15-16 | 23.36 ± 2.52 | 26.22 ± 2.97 | 49.35 ± 3.21 | 20.86 ± 2.22 | 23.23 ± 2.57 | 43.84 ± 3.41 |
| | 14-15-16 | 19.79 ± 2.25 | 27.52 ± 2.46 | 47.09 ± 3.03 | 21.48 ± 2.52 | 27.68 ± 1.99 | 48.93 ± 3.20 |
| Tetrads - Non-Canonical strand breaks | | | | | | | |
| **Single SB** | 2-3 | 21.03 ± 3.16 | 28.27 ± 1.85 | 49.05 ± 3.49 | 22.02 ± 2.93 | 27.36 ± 1.94 | 49.12 ± 3.17 |
| | 3-4 | 20.46 ± 3.10 | 27.71 ± 2.24 | 47.93 ± 3.17 | 19.93 ± 2.96 | 28.02 ± 1.82 | 47.65 ± 3.37 |
| | 9-10 | 20.14 ± 3.18 | 28.05 ± 1.78 | 47.97 ± 3.77 | 20.86 ± 2.22 | 28.76 ± 2.22 | 49.38 ± 2.68 |
| | 14-15 | 22.37 ± 2.78 | 28.73 ± 1.95 | 50.84 ± 3.09 | 20.63 ± 2.98 | 29.47 ± 2.14 | 49.78 ± 3.27 |
| | 15-16 | 21.49 ± 2.18 | 27.94 ± 2.34 | 49.22 ± 2.66 | 21.04 ± 2.30 | 28.86 ± 2.18 | 49.69 ± 2.76 |
| | 21-22 | 20.24 ± 2.04 | 27.99 ± 2.49 | 48.01 ± 2.86 | 21.13 ± 2.16 | 26.64 ± 2.19 | 47.56 ± 2.54 |
| **Double SB** | 2-3/14-15 | 23.48 ± 3.09 | 28.21 ± 1.77 | 51.40 ± 3.31 | 20.14 ± 3.40 | 31.37 ± 3.16 | 51.27 ± 2.98 |
| | 2-3-4 | 21.89 ± 2.84 | 27.84 ± 1.87 | 48.96 ± 2.87 | 25.09 ± 2.95 | 31.72 ± 1.97 | 56.49 ± 3.22 |
| | 3-4/9-10 | 20.78 ± 2.45 | 29.14 ± 2.68 | 49.59 ± 2.55 | 20.11 ± 3.05 | 27.55 ± 2.67 | 47.30 ± 4.19 |
| | 3-4/14-15 | 20.89 ± 3.02 | 29.16 ± 2.59 | 49.69 ± 3.49 | 18.00 ± 3.25 | 33.19 ± 2.75 | 50.89 ± 2.61 |
| | 3-4/15-16 | 20.14 ± 2.51 | 28.43 ± 2.72 | 48.16 ± 3.42 | 21.84 ± 2.48 | 28.61 ± 2.69 | 50.14 ± 2.98 |
| | 14-15-16 | 23.30 ± 2.66 | 31.51 ± 2.89 | 54.49 ± 3.56 | 26.36 ± 2.94 | 29.59 ± 2.56 | 55.67 ± 3.30 |

**Table S1 :** Average and standard deviation of the twist angle (Ψ) parameter for all simulated systems. The Ψ values are reporter for each couple of quartets find into the G4. The red text refers to the simulation in which the G-quadruplex was destabilized.

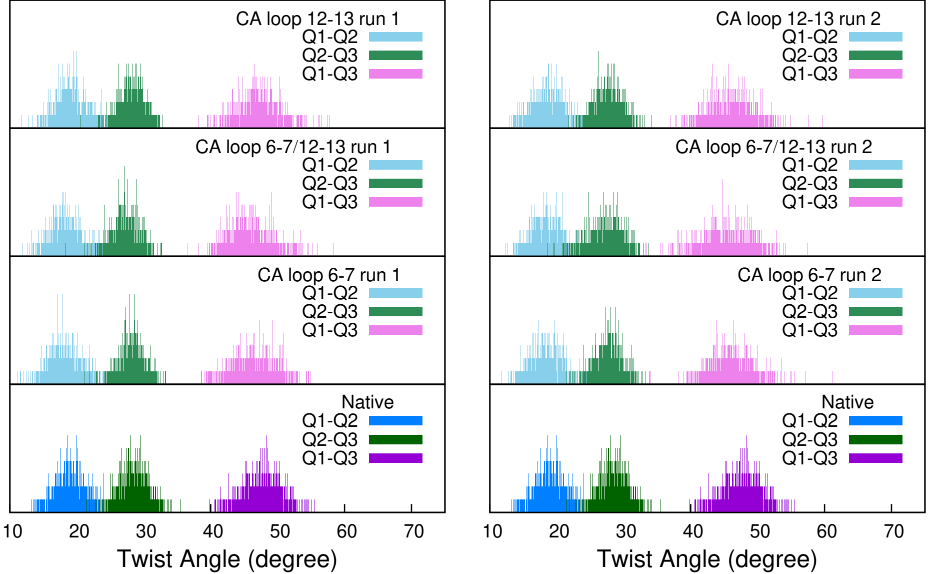


Figure S57 - Frequence distribution of the twist angles of the loops strand breaks systems.

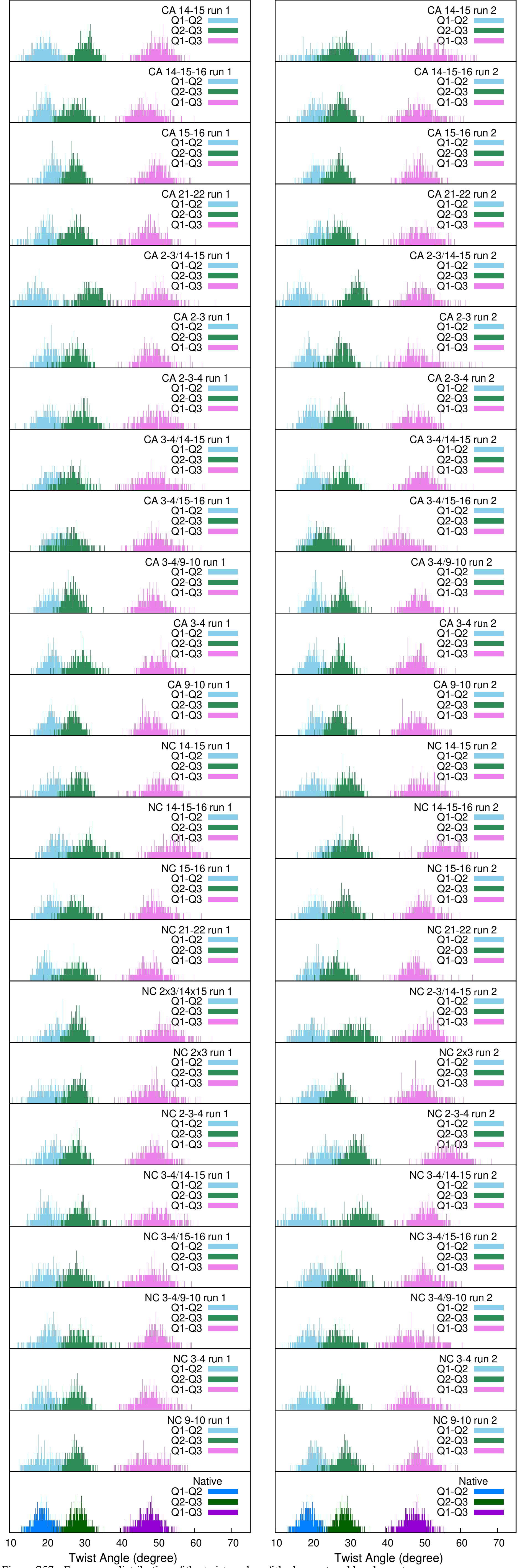


Figure S57 - Frequence distribution of the twist angles of the loops strand breaks systems.

| Bonded nucleotides | Non-bonded nucleotides |
|---|---|
| 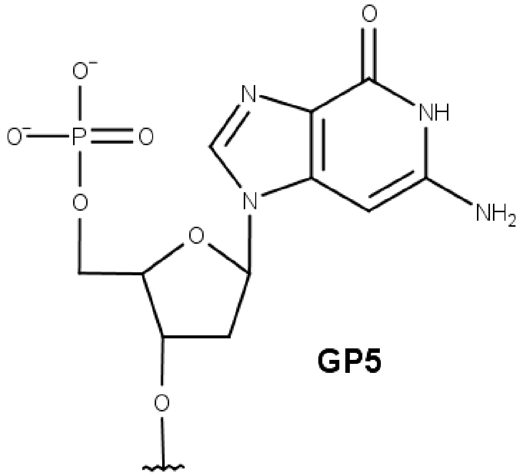  **GP5**<br>*Modified from guanosine (DG). Canonical type.* | 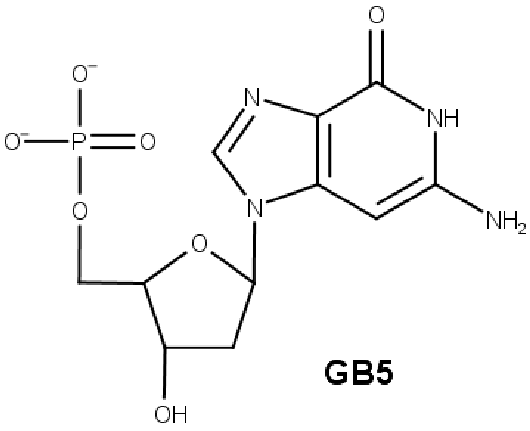  **GB5**<br>*Modified from guanosine (DG). Canonical type.* |
| 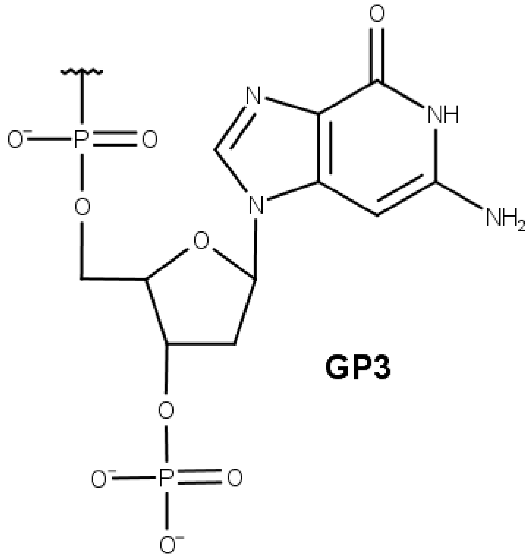 **GP3**<br>*Modified from guanosine (DG). Non-canonical type.* | 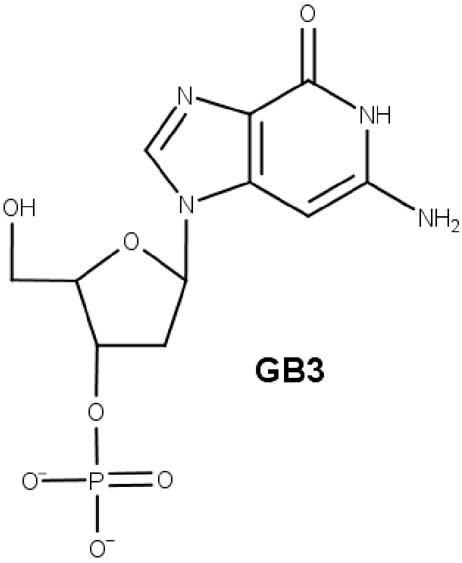  **GB3**<br>*Modified from guanosine (DG). Non-canonical type.* |
| 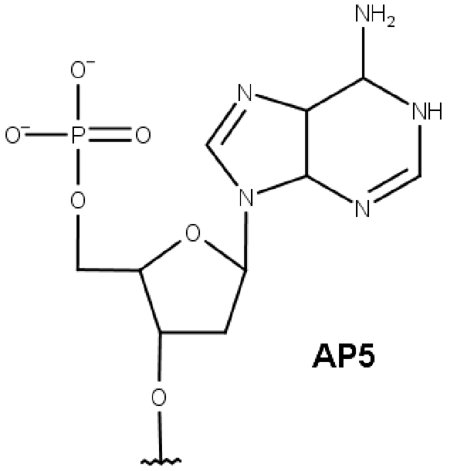 **AP5**<br>*Modified from adenosine (DA). Canonical type.* | / |

**Table S2 –** Chemical structures of modified nucleotides, corresponding to a modification of the guanosine (GP3, GB3, GP5, GB5) or the adenosine (AP5). In our nomenclature, the end number mean the position of the phosphate group, in 3' or in 5'. Non-bonded nucleotides are involved in vertical double strand breaks.
